# Gut complement C1q and C3 interact with *Clostridioides difficile* spores via CdeM and contribute to pathogenesis

**DOI:** 10.64898/2026.09.24.753972

**Authors:** Marjorie Pizarro-Guajardo, Christian Brito-Silva, Pablo Castro-Córdova, Nicolás Montes-Bravo, Eduardo Callegari, Fernando Gil, Daniel Paredes-Sabja

**Affiliations:** Department of Biology, Texas A&M University, College Station, Texas, USA; ANID – Millennium Science Initiative Program - Millennium Nucleus in the Biology of the Intestinal Microbiota, Santiago, Chile; IMPACT, Center of Interventional Medicine for Precision and Advanced Cellular Therapy, Santiago, Chile; Laboratory of Nano-Regenerative Medicine, Centro de Investigación e Innovación Biomédica (CiiB), Faculty of Medicine, Universidad de los Andes, Santiago, Chile; Division of Biomedical and Translational Sciences, Sanford School of Medicine, University of South Dakota.; Microbiota-Host Interactions & Clostridia Research Group, Center for Biomedical Research and Innovation (CIIB), Universidad de los Andes, Santiago 7620001, Chile; School of Medicine, Faculty of Medicine, Universidad de los Andes, Santiago 7620001, Chile

**Keywords:** C. difficile, gut complement, C1q, C3, CdeM, exosporium

## Abstract

*Clostridioides difficile* infection (CDI) is characterized by toxin-mediated epithelial injury, intestinal inflammation, and a high frequency of disease recurrence. Although complement components are produced locally within the gastrointestinal tract, their contribution to *C. difficile* pathogenesis remains poorly understood. Here, we investigated the interaction of the complement proteins C1q and C3 with *C. difficile* spores and their contribution to spore–host interactions and disease outcome. We found that C1q and C3 are accessible within the healthy ileal mucosa and that TcdB intoxication differentially remodels their spatial availability, decreasing accessible C1q while increasing accessible C3 at higher toxin concentrations. *C. difficile* spores associated with both complement components in vivo and directly interacted with purified human C1q and C3 in vitro, with immunogold electron microscopy localizing these interactions to the outer exosporium. Although the collagen-like BclA proteins modulated complement binding under some conditions, they were not required for C1q or C3 association. Far-Western analysis coupled to mass spectrometry identified the exosporium proteins CotE and CdeM as candidate complement-interacting proteins, and purified CdeM inclusion bodies directly associated with both C1q and C3. Exposure of spores to human serum promoted their interaction with intestinal epithelial cells, whereas depletion of either C1q or C3 markedly reduced spore internalization and reconstitution with the corresponding purified protein partially restored entry. C3-derived species, including C3dg, were selectively associated with spores but not vegetative cells, without reducing spore viability. Finally, C1q deficiency did not alter the initial onset of CDI but accelerated recovery and markedly attenuated recurrent disease following vancomycin treatment. Together, these findings identify intestinal complement as a previously unrecognized component of the *C. difficile* spore–host interface and reveal C1q and C3 as modulators of spore epithelial entry, persistence, and recurrent disease.

**Author summary:** *Clostridioides difficile* is a major cause of antibiotic-associated diarrhea, and recurrent infections remain difficult to prevent. Disease recurrence depends in part on highly resistant spores that persist in the intestine and can interact with the intestinal lining. Here, we show that two components of the complement immune system, C1q and C3, are present and accessible in the intestinal mucosa and can associate directly with *C. difficile* spores. We identified the spore surface protein CdeM as one molecular component capable of interacting with both C1q and C3. These complement proteins also enhanced the ability of spores to enter intestinal epithelial cells, suggesting that a host defense pathway can be exploited by the pathogen to promote interactions with the intestinal barrier. C3 was deposited and processed on spores but did not reduce their viability. Importantly, mice lacking C1q recovered more rapidly from infection and developed markedly less recurrent disease after antibiotic treatment. These findings reveal an unexpected role for intestinal complement in *C. difficile* pathogenesis and suggest that complement–spore interactions may contribute to persistence and recurrence.

## Introduction

*Clostridioides difficile* is a Gram-positive, strict anaerobe, opportunistic pathogenic bacterium spore-former that causes *C. difficile* infection (CDI) that cause antibiotics associated diarrhea [1, 2]. The clinical symptoms vary from mild diarrhea to pseudomembranous colitis, and in most fatal cases, to toxic megacolon and death [3–8]. The mortality rate of CDI is ∼5 %, but in several outbreaks, mortality may increase to 20% [1]. Although the current standard of care treatments consisting of vancomycin, metronidazole, and recently fidaxomicin are effective against the first episode of CDI, recurrence of the disease after a first, second and third episode may reach up to 20%, 40%, and 60% respectively [9, 10]. During antibiotic therapy, the intestinal microbiota diversity becomes depleted, reducing colonization resistance against *C. difficile* [4–6], allowing ingested or endogenous dormant *C. difficile* spores to germinate and colonize intestinal mucosal surfaces and secrete two major toxins TcdA and TcdB also an auxiliary toxin (CdtAB), which provoke epithelial damage leading to local inflammation and disruption of the epithelial barrier and the clinical manifestation of the disease [11–14]. *C. difficile* initiates a sporulation cycle leading to the formation of new metabolically dormant spores [15, 16], considered the main factor of R-CDI [15]. Recent work supports this notion, as removal of *C. difficile* spore-during CDI by anti-spore chicken antibodies, or by blocking spore-formation during CDI, reduces recurrence [17, 18]. Our group has also demonstrated that C. difficile spores enter the intestinal mucosa, and that inhibiting spore entry reduces R-CDI [19].

The mechanisms through which *C. difficile* spores interact with the intestinal mucosa to persist are poorly understood. Initial studies demonstrated that *C. difficile* spores had high binding to intestinal epithelial cells (IECs) *in vitro* [20]. *C. difficile* spores also interact with high affinity to components of the intestinal mucosa, including mucin and extracellular matrix proteins, fibronectin and vitronectin [21]. Notably, both, Recently, Castro-Córdova et al. (2021) demonstrated that *C. difficile* spores gain entry into IECs via fibronectin-α_5_β_1_and vitronectin-α_v_β_1_in a BclA3-dependent manner and that spore-entry into the intestinal barrier contributes to disease recurrence in a mouse model of the disease [19]. Moreover, *C. difficile* spores interact with E-cadherin, an adherens junction protein, through the exosporium proteins CdeM and CotE, and this interaction contributes to spore adherence to and internalization into intestinal epithelial cells [22, 23].

In addition to host cellular surface molecules, components of the innate immune system can also be hijacked by pathogens to invade host cells or evade the host immune system. The complement system consists of more than fifty secreted proteins and membrane receptors that work in a highly coordinated manner as a first line of defense against infection of host tissues by microbes [24]. In the blood, complement activation is initiated through the classical, lectin, or alternative pathways, all of which converge at the level of C3 cleavage. Proteolytic activation of C3 generates the soluble anaphylotoxin C3a and the opsonin C3b, whose exposed thioester permits covalent deposition on microbial surfaces. Surface-bound C3b and its proteolytic derivatives promote opsonophagocytic recognition by complement receptors expressed on neutrophiles and macrophages, facilitating microbial uptake and clearance. Continued complement activation also generates C5 convertase and initiates the terminal pathway, culminating in assembly of the C5b-9 membrane attack complex /MAC), which can directly lyse susceptible Gram-negative bacteria. Traditionally, the complement has been considered a predominantly hepatocyte-derived, plasma-resident innate immune system, with circulating components distributed throughout the vascular and interstitial compartments [25]. However, studies have shown that complement system is also produced outside of the liver, by immune, intestinal epithelial and lung epithelial cells [26–31]. Complement cascade genes are also upregulated in rectal biopsies from patience with inflammatory bowel disease relative to healthy individuals [32]. During CDI, *in vivo* transcriptional profiling experiments demonstrate that the complement components C1q and C3 are upregulated in response to *C. difficile* TcdA and TcdB [33].

The complement system is targeted and exploited by several pathogens as a mechanism of immune modulation and host-cell interactions. *Bacillus anthracis* spores activate the classical complement pathway [34] and complement components C3 and C3b associate with the spore surface [35]. Both C1q and C3 enhance spore uptake by macrophages, while deposition of C3-derived fragments on the spore surface is dependent is dependent on C1q and the exosporium protein BclA and occurs independently of IgG [34]. *B. anthracis* spores also directly recruit complement factor H (CFH) through its spore-surface protein BclA, a mechanism that limits complement activation and attenuates antibody responses against spores in vivo, suggesting that BclA contributes to immune evasion during *B. anthracis* infection [36]. In addition, *B. anthracis* exploits C1q to promote spore entry into intestinal epithelial cells through a BclA-C1q-α_2_β_1_ integrin dependent mechanism [37].

In contrast, comparatively little is known about the interactions of C. difficile with the complement system. One study where identified CD2831, a member of the Microbial Surface Components Recognizing Adhesive Matrix Molecules, as a C1q binding protein that interacts with the collagen-like region of human C1q [38]. This interaction has been proposed to interfere with activation of classical complement pathway, suggesting a dual function for CD2831 in adhesion to collagen-rich host tissues and complement mediated immune evasion. Intestinal C1q has also been detected in a murine model of chemically induced diarrhea and intestinal inflammation, in which increased tissue injury was accompanied by elevated C1q levels in serum and intestinal tissues [39]. These observations support the possibility that, during CDI, epithelial barrier disruption and pseudo membrane-associated inflammation may increase the exposure of accessibility of complement components within the intestinal mucosa, hence, creating conditions that favor interactions between *C. difficile* spores and C1q [40].

In this study, we define a previously unrecognized interface between intestinal complement and *C. difficile* spores. We show that C1q and C3 are accessible within the healthy intestinal mucosa and that their distribution is remodeled by TcdB-induced epithelial injury. *C. difficile* spores associate with both complement components in vivo and bind purified C1q and C3 through the spore exosporium, with the cysteine-rich morphogenetic protein CdeM as a major complement-interacting factor. We further demonstrate that C1q and C3 enhance spore internalization into intestinal epithelial cells, whereas C3 is deposited and processed on the spore surface without compromising spore viability. Finally, C1q deficiency alters recovery and recurrent disease after CDI, supporting a previously unrecognized role for intestinal complement in *C. difficile* persistence and pathogenesis.

## Results

### C1q and C3 are accessible in the healthy ileum mucosa

Although the liver is the major source of circulating complement proteins, complement components are also synthesized locally within the intestine [41]. C3 is produced by intestinal stromal and epithelial cells and is present in the intestinal lumen, whereas macrophages constitute a major local source of C1q [42, 43]. However, whether C1q and C3 are accessible at the luminal interface of the healthy ileal epithelium, where they could encounter *C. difficile*, remains unclear. We therefore examined the distribution and accessibility of C1q and C3 in the healthy ileal mucosa. To distinguish accessible from total complement, intact ileal tissue was first immunostained under non-permeabilizing conditions to detect accessible C1q or C3 and was subsequently permeabilized and stained to detect the corresponding total complement pool.

Three-dimensional confocal microscopy revealed a heterogeneous distribution of C1q throughout the ileal mucosa (Fig. 1A). Quantification showed that 14.8% of intestinal epithelial cells were C1q-positive, whereas 85.2% showed no detectable C1q staining (Fig. 1C). Among C1q-positive cells, 44.5% contained C1q that was accessible before permeabilization, whereas 55.5% contained C1q detectable only after permeabilization (Fig. 1D).

**Fig. 1.**
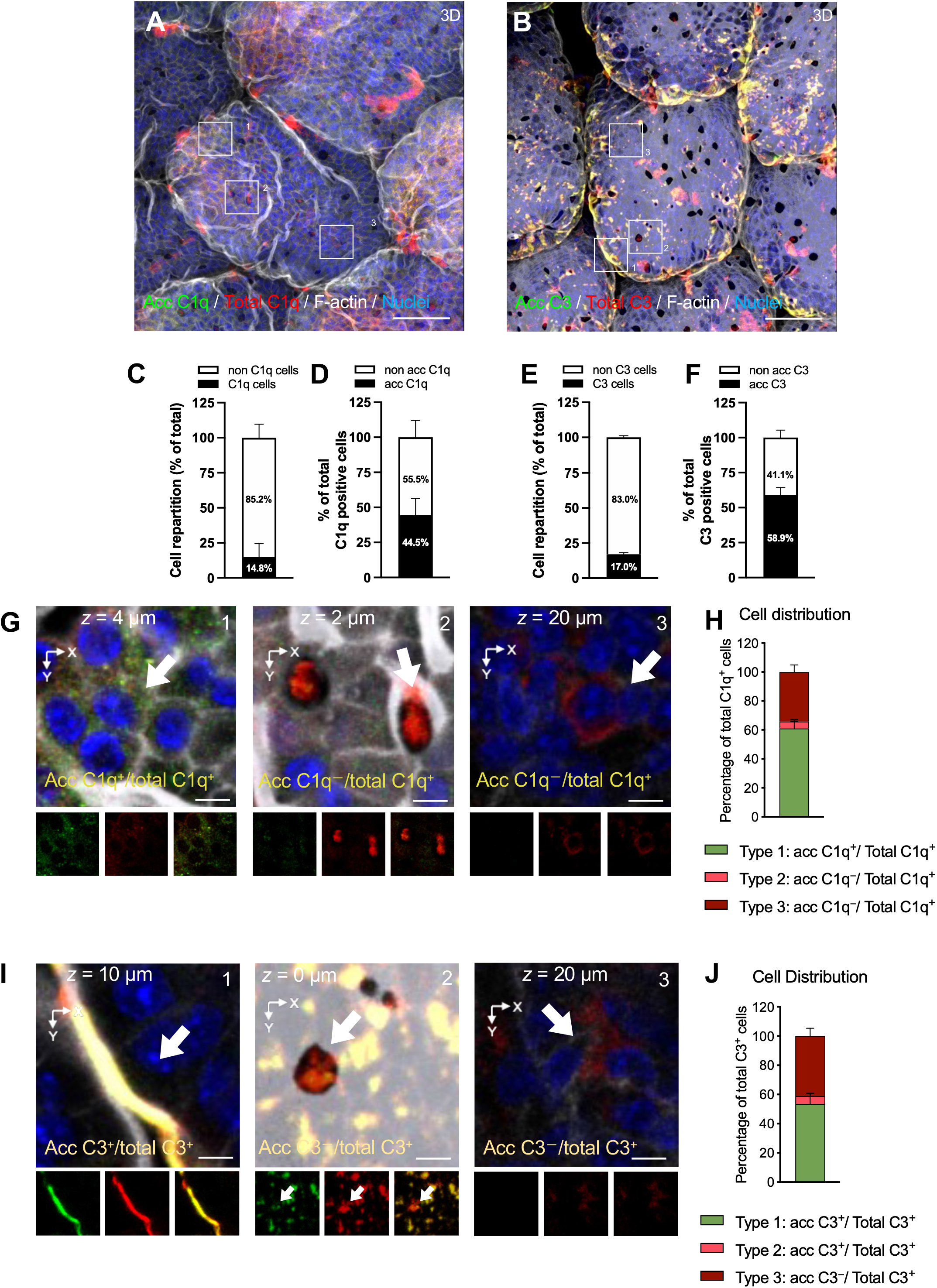
C1q and C3 are accessible in healthy ileum mucosa. **A, B,** Representative three-dimensional confocal micrographs of fixed whole-mount ileal mucosa from healthy mice (*n* = 3) immunostained for **A**, accessible C1q (acc C1q) and total C1q, or **B**, accessible C3 (acc C3) and total C3. Accessible C1q and C3 are shown in green, total C1q and C3 in red, F-actin in gray, and nuclei in blue. Fluorophore colors were digitally reassigned for visualization. Numbered boxes indicate representative regions corresponding to the three cellular staining patterns shown at higher magnification in **G** and **I**. **C,** Distribution of ileal epithelial cells according to total C1q immunoreactivity. **D,** Distribution of C1q-positive cells according to the presence or absence of accessible C1q. **E,** Distribution of ileal epithelial cells according to total C3 immunoreactivity. **F,** Distribution of C3-positive cells according to the presence or absence of accessible C3. A total of 448 C1q-positive cells from 12 villi and 373 C3-positive cells from 7 villi were analyzed. **G,** Representative high-magnification confocal images illustrating the three C1q staining patterns identified in the ileal mucosa: **type 1**, epithelial cells exhibiting accessible and total C1q (acc C1q /total C1q); **type 2**, cells lacking detectable accessible C1q but positive for total C1q after permeabilization (acc C1q /total C1q); and **type 3**, cells located beneath the villus surface that were negative for accessible C1q but positive for total C1q following permeabilization (acc C1q /total C1q). Arrows indicate representative cells or regions corresponding to each staining pattern. Images below each panel show the individual accessible and total C1q fluorescence channels and the merged image. The indicated *z*-position corresponds to the optical plane shown. **H,** Relative distribution of the three C1q-positive cellular staining patterns. **I,** Representative high-magnification confocal images illustrating the corresponding three C3 staining patterns: **type 1**, cells exhibiting accessible and total C3 (acc C3 /total C3); **type 2**, cells lacking detectable accessible C3 but positive for total C3 after permeabilization (acc C3 /total C3); and **type 3**, cells located beneath the villus surface that were negative for accessible C3 but positive for total C3 following permeabilization (acc C3 /total C3). Arrows indicate representative cells or regions corresponding to each staining pattern. Images below each panel show the individual accessible and total C3 fluorescence channels and the merged image. **J,** Relative distribution of the three C3-positive cellular staining patterns. Data in **C–F, H, and J** are presented as mean ± SEM. Scale bars: **A, B, 50** μ**m; G, I, 5** μ**m.**

Closer examination of C1q-positive cells revealed three recurrent spatial patterns (Fig. 1G). Type 1 cells displayed a defined apical F-actin architecture and contained both accessible and total C1q (acc C1q^+^/total C1q^+^). Type 2 cells contained total C1q but little or no detectable accessible C1q at the epithelial surface (acc C1q^−^/total C1q^+^). Type 3 cells were located beneath the villus surface and similarly lacked accessible C1q while remaining positive for total C1q after permeabilization (acc C1q^−^/total C1q^+^). Quantification of these morphologies showed that Type 1 cells represented 61.0% of the C1q-positive population, whereas Type 2 and Type 3 cells represented 4.8% and 34.1%, respectively (Fig. 1H). Thus, although C1q was present in several spatial compartments within the villus, the predominant C1q-positive morphology corresponded to cells in which C1q was accessible at the epithelial interface.

C3 exhibited a related distribution (Fig. 1B). Overall, 17.0% of intestinal epithelial cells were C3-positive, whereas 83.0% showed no detectable C3 staining (Fig. 1E). Among C3-positive cells, 58.9% contained accessible C3, whereas 41.1% contained C3 detectable only following permeabilization (Fig. 1F). Three spatial patterns were again evident (Fig. 1I). Type 1 cells contained both accessible and total C3 (acc C3^+^/total C3^+^); Type 2 cells were negative for accessible C3 but positive for total C3 (acc C3^−^/total C3^+^); and Type 3 cells located beneath the villus surface were negative for accessible C3 but positive for total C3 following permeabilization. Type 1 cells accounted for 53.5% of the C3-positive population, whereas Type 2 and Type 3 cells represented 5.4% and 41.1%, respectively (Fig. 1J).

Together, these results demonstrate that C1q and C3 are heterogeneously distributed within the healthy ileal mucosa and occupy both accessible and non-accessible tissue compartments. Importantly, the predominant C1q-and C3-positive epithelial morphology contained complement that was accessible without permeabilization, identifying discrete sites within the healthy ileal mucosal interface where these complement components could potentially encounter luminal microorganisms, including *C. difficile*.

### TcdB differentially alters accessible and total C1q and C3 in the ileal mucosa

Having established that C1q and C3 are accessible within the healthy ileal mucosa, we next asked whether toxin-induced epithelial injury alters this complement landscape. Because TcdB disrupts intestinal epithelial architecture during CDI [33], ileal loops were exposed to increasing amounts of TcdB (0.1, 0.5, 1, and 5 μg), and accessible and total C1q and C3 were evaluated by confocal microscopy. Increasing TcdB exposure produced marked alterations in epithelial architecture that were accompanied by changes in the distribution of both complement components (Fig. 2A,D).

**Fig. 2.**
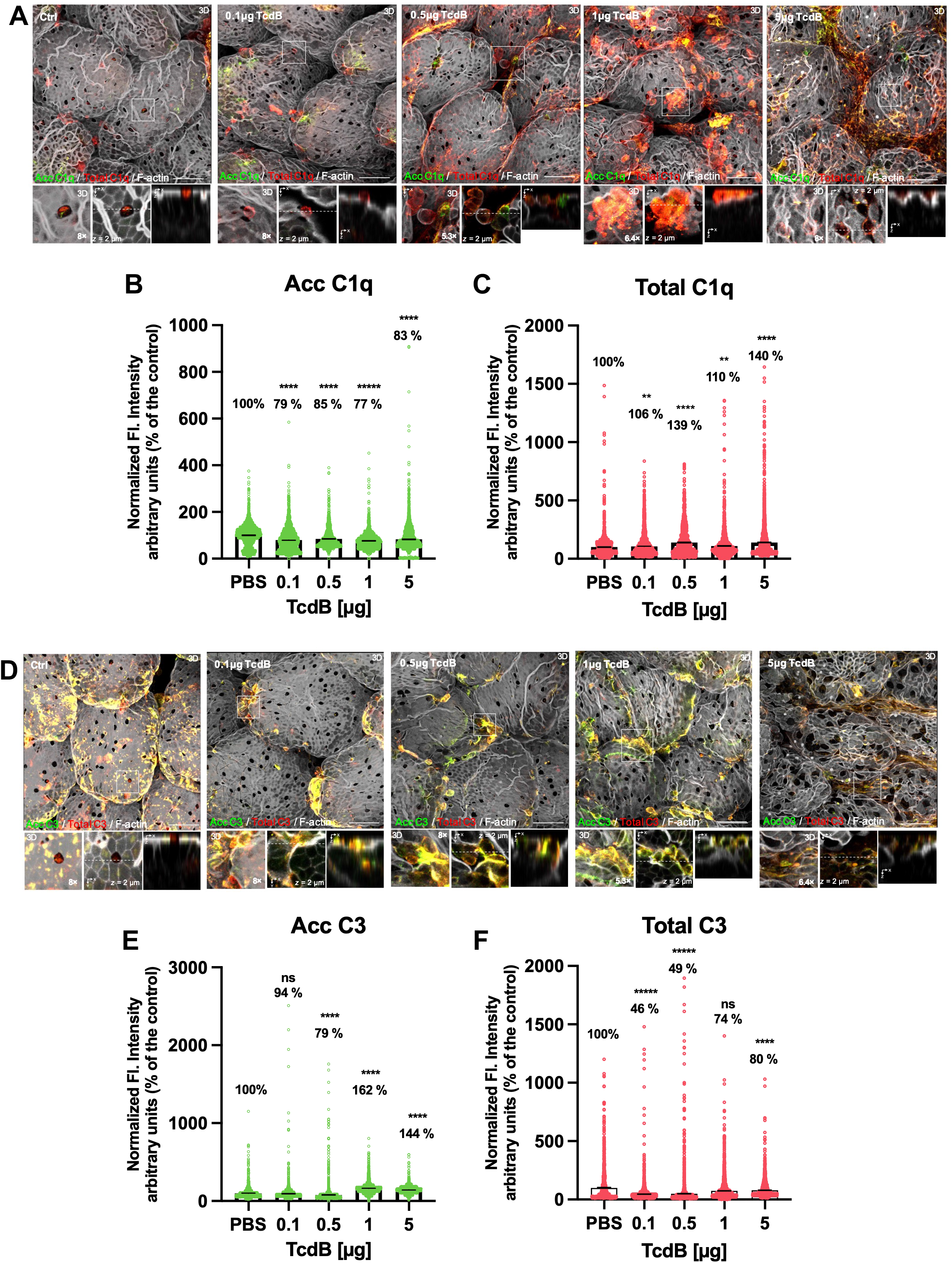
TcdB differentially alters accessible and total C1q and C3 in the ileal mucosa. Ligated ileal loops were exposed to PBS (control) or increasing amounts of *Clostridioides difficile* toxin B (TcdB; 0.1, 0.5, 1, or 5 μg) and analyzed for accessible (acc) and total C1q or C3. **A,** Representative three-dimensional confocal micrographs of ileal mucosa immunostained for accessible C1q (green) and total C1q (red); F-actin is shown in gray. Boxed regions are shown at higher magnification below each image together with orthogonal views illustrating the spatial distribution of C1q within the epithelial architecture. **B,** Quantification of accessible C1q fluorescence intensity. **C,** Quantification of total C1q fluorescence intensity. **D,** Representative three-dimensional confocal micrographs of ileal mucosa immunostained for accessible C3 (green) and total C3 (red); F-actin is shown in gray. Boxed regions are shown at higher magnification below each image together with orthogonal views. **E,** Quantification of accessible C3 fluorescence intensity. **F,** Quantification of total C3 fluorescence intensity. For **B, C, E,** and **F**, fluorescence intensity was normalized to the corresponding PBS-treated control, which was set to 100%. Percentages above each condition indicate the mean fluorescence intensity relative to PBS. Accessible C1q decreased following TcdB exposure, whereas total C1q was maintained or increased. Accessible C3 exhibited a dose-dependent biphasic response, with reduced signal at 0.5 μg and increased signal at 1 and 5 μg TcdB, whereas total C3 was reduced following TcdB exposure. Individual points represent pixels/cells, and violin plots show the distribution of normalized fluorescence intensity values. Statistical significance was determined using one-way ANOVA with Sidak’s multiple-comparisons test; ns, not significant; **P < 0.01; ****P < 0.0001. Scale bars are indicated in the micrographs.

Accessible C1q was significantly reduced at all TcdB doses, reaching 79%, 85%, 77%, and 83% of the PBS control at 0.1, 0.5, 1, and 5 μg TcdB, respectively (Fig. 2B). In contrast, total C1q fluorescence was maintained or increased, reaching 106%, 139%, 110%, and 140% of control at the corresponding doses (Fig. 2C). Thus, TcdB produced opposing effects on accessible and total C1q, with reduced accessible C1q despite preservation or accumulation of the total tissue-associated C1q pool.

C3 exhibited a distinct response. Accessible C3 remained comparable to control at 0.1 μg TcdB (94%), decreased to 79% at 0.5 μg, and increased markedly at the higher toxin doses, reaching 162% and 144% of control at 1 and 5 μg TcdB, respectively (Fig. 2E). Conversely, total C3 fluorescence was reduced to 46% and 49% of control at 0.1 and 0.5 μg TcdB, respectively, and remained below control levels at 1 and 5 μg TcdB (74% and 80%, respectively), although the difference at 1 μg was not statistically significant (Fig. 2F).

Together, these findings demonstrate that TcdB does not uniformly increase or decrease complement within the ileal mucosa, but instead differentially alters the accessible and total pools of C1q and C3. Whereas accessible C1q decreased despite preservation or increased total C1q, higher TcdB doses increased accessible C3 despite reduced total C3. These results indicate that TcdB-induced epithelial disruption alters the spatial availability of complement components at the mucosal interface, potentially modifying the complement environment encountered by C. difficile during intestinal injury.

### *C. difficile* spores associates to accessible C1q and C3 in healthy intestinal mucosa *in vivo*

Having established that C1q and C3 are accessible within the healthy ileal mucosa and that TcdB alters their mucosal distribution, we next asked whether *C. difficile* spores encounter and associate with these accessible complement components in vivo. To address this question, *C. difficile* R20291 spores were introduced into ligated mouse ileal loops and incubated for 5 h. Ileal tissues were subsequently immunostained under non-permeabilizing conditions to detect accessible C1q or C3 and analyzed by three-dimensional confocal microscopy (Fig. 3A,E).

**Fig. 3.**
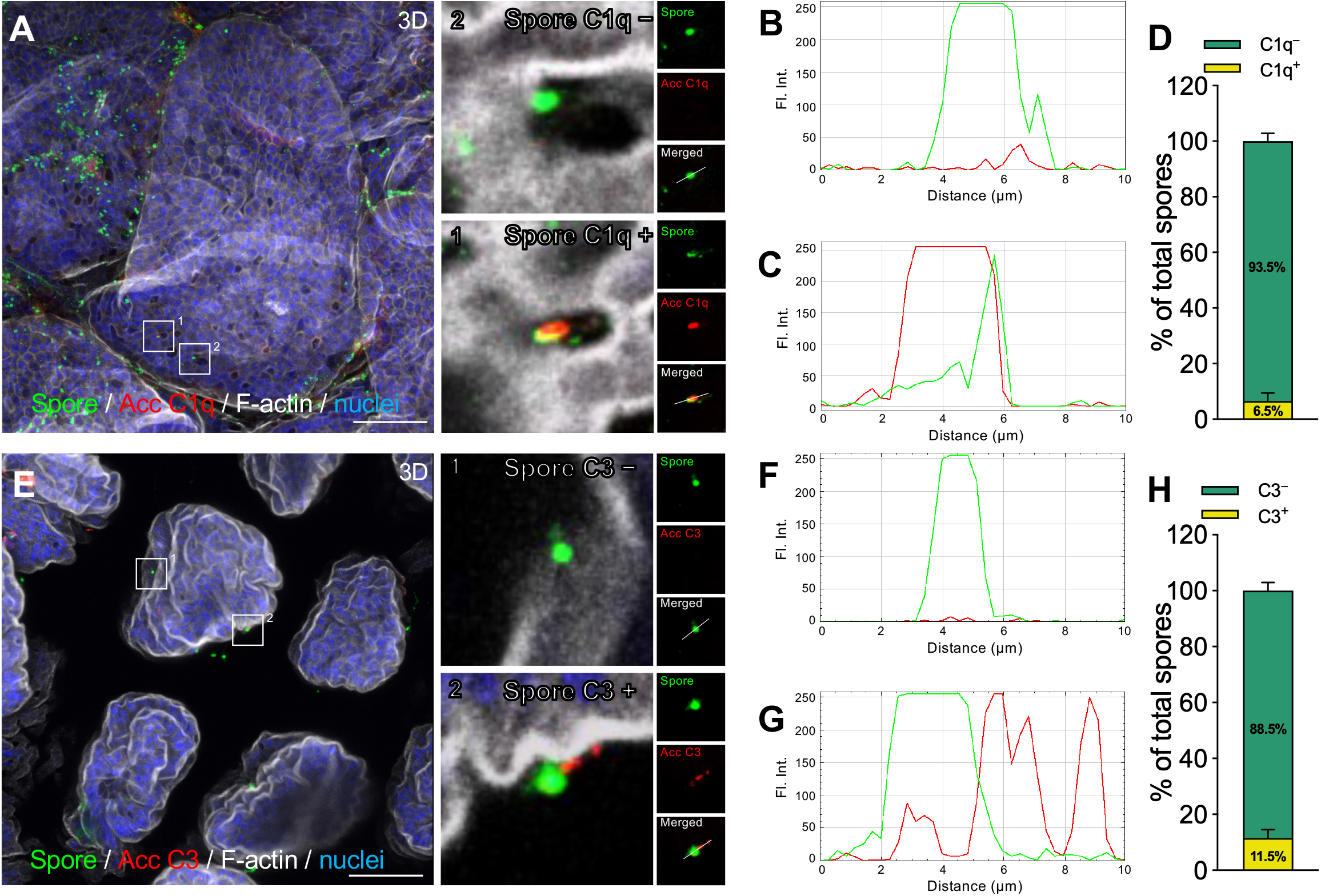
C1q and C3 associate with *C. difficile* spores in the ileal mucosa. The ileum of C57BL/6 mice (n = 5) was inoculated with 5 × 10^8 *C. difficile* R20291 spores for 5 h using a ligated ileal loop model. **A,** Representative three-dimensional confocal projection of fixed whole-mount ileal mucosa immunostained for *C. difficile* spores (green), accessible C1q (acc C1q; red), F-actin (gray), and nuclei (blue). Enlarged regions on the right show representative spores classified as non-associated with C1q (C1q−) or associated with C1q (C1q+). **B, C,** Fluorescence-intensity line profiles of representative C1q− (**B**) and C1q+ (**C**) spores. The green line represents spore fluorescence intensity and the red line represents accessible C1q fluorescence intensity. **D,** Distribution of spores classified as C1q-associated or non-C1q-associated. **E,** Representative three-dimensional confocal projection of fixed whole-mount ileal mucosa immunostained for *C. difficile* spores (green), accessible C3 (acc C3; red), F-actin (gray), and nuclei (blue). Enlarged regions on the right show representative spores classified as non-associated with C3 (C3−) or associated with C3 (C3+). **F, G,** Fluorescence-intensity line profiles of representative C3− (**F**) and C3+ (**G**) spores. The green line represents spore fluorescence intensity and the red line represents accessible C3 fluorescence intensity. **H,** Distribution of spores classified as C3-associated or non-C3-associated. Fluorophore colors were digitally reassigned for visualization. An area of 84,628 µm² was analyzed in each mouse. Data are presented as mean ± SEM. Scale bar, 50 µm.

Confocal imaging revealed *C. difficile* spores distributed along the ileal mucosa, including a subset associated with accessible C1q (Fig. 3A). Higher-magnification images and fluorescence intensity line profiles distinguished spores lacking detectable C1q association, in which the spore fluorescence signal occurred without a corresponding accessible C1q signal (Fig. 3A,B), from C1q-associated spores, in which accessible C1q fluorescence was detected at the position of the spore (Fig. 3A,C). Quantification showed that 6.5% of the total spores were associated with accessible C1q, whereas 93.5% showed no detectable C1q association (Fig. 3D).

A similar pattern was observed for accessible C3 (Fig. 3E). Higher-magnification images and fluorescence intensity profiles identified spores without detectable C3 association (Fig. 3E,F) and spores associated with accessible C3, characterized by C3 fluorescence spatially coincident with or immediately adjacent to the spore signal (Fig. 3E,G). Quantification showed that 11.5% of the total spores were associated with accessible C3, whereas 88.5% showed no detectable C3 association (Fig. 3H).

Together, these results demonstrate that a subset of *C. difficile* spores associates with accessible C1q and C3 within the ileal mucosa *in vivo*. These findings establish that the accessible complement pools identified at the intestinal mucosal interface can encounter *C. difficile* spores and prompted us to next determine whether C1q and C3 can directly bind to the spore surface.

### Human C1q and C3 bind to *C. difficile* spores *in vitro*

Because *C. difficile* spores associated with accessible C1q and C3 in the ileal mucosa in vivo, we next asked whether these complement proteins can directly interact with the spore surface. Solid-phase binding assays using purified human C1q demonstrated concentration-dependent binding to spores of both *C. difficile* strains 630Δ*ermB* and R20291, with little binding to the BSA control (Fig. 4A). Consistent with these results, immunofluorescence microscopy showed C1q-associated fluorescence on R20291 spores following incubation with purified C1q (Fig. 4B). Quantification of individual spores showed a significant increase in C1q-associated fluorescence relative to the control, with the mean fluorescence increasing from 100 to approximately 362% of control (Fig. 4C). However, C1q association was heterogeneous across the spore population: approximately 45% of spores were C1q-associated, whereas the remaining ∼55% showed little or no detectable C1q signal (Fig. 4D).

**Fig. 4.**
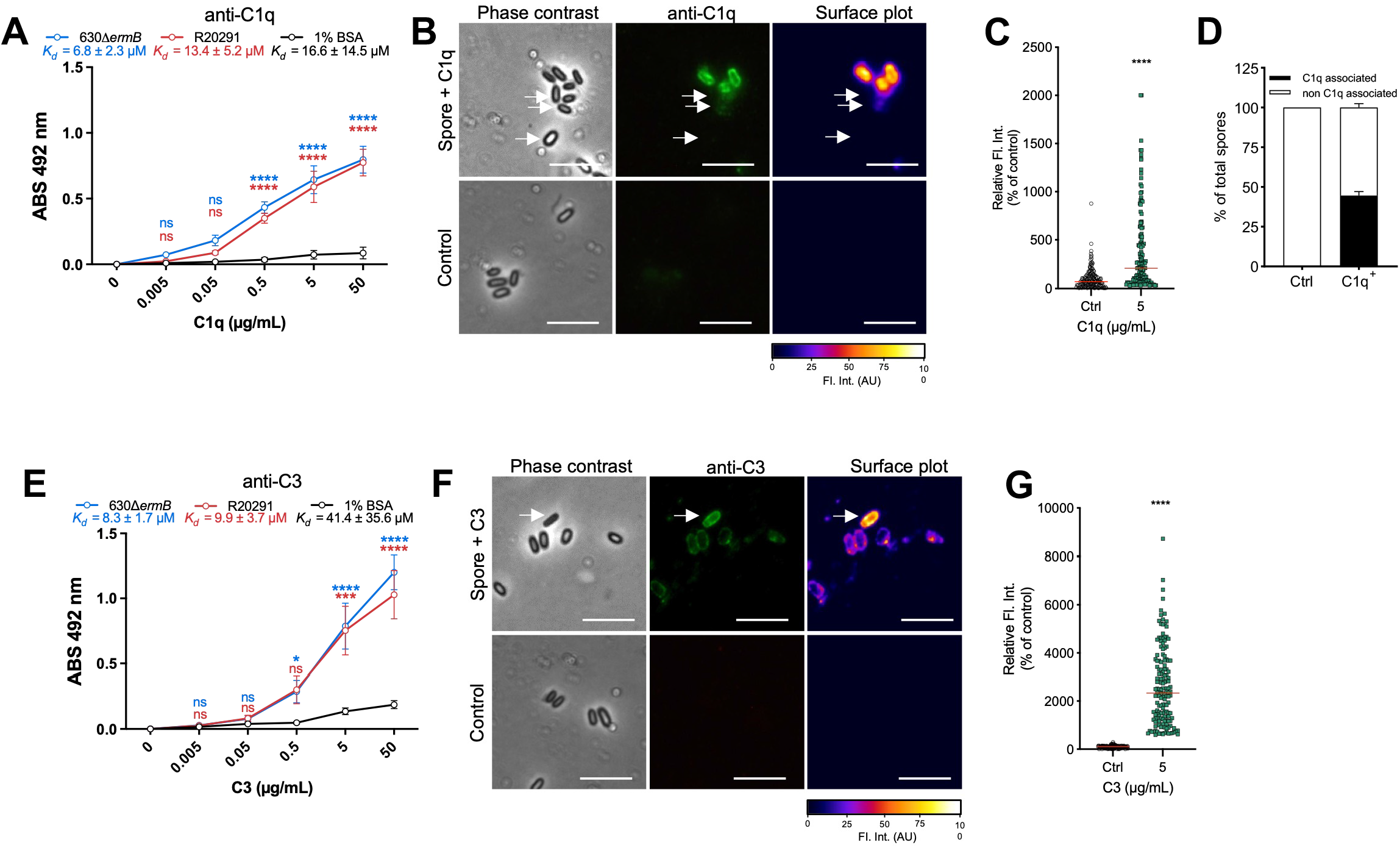
Purified human C1q and C3 bind to *C. difficile* spores. **A,** Solid-phase binding assay of purified human C1q to *C. difficile* spores. Wells coated with 1.6 × 10^7^ spores of strains 630Δ*ermB* or R20291, or with 1% BSA as a negative control, were incubated with increasing concentrations of purified human C1q. Bound C1q was detected by immunoassay, and absorbance was measured at 492 nm. Apparent dissociation constants (*K*d) calculated from the binding curves are indicated. Data represent the mean ± SEM of four independent wells from two independent experiments. **B,** Representative phase-contrast, anti-C1q immunofluorescence, and fluorescence-intensity surface plots of R20291 spores incubated with purified C1q or control buffer. Arrows indicate representative C1q-associated and non-C1q-associated spores. **C,** Quantification of C1q-associated fluorescence intensity in individual R20291 spores incubated without C1q (Ctrl) or with 5 μg/mL purified C1q. Each point represents an individual spore. **D,** Distribution of spores classified as C1q-associated or non-C1q-associated following incubation in the absence (Ctrl) or presence of purified C1q. **E,** Solid-phase binding assay of purified human C3 to *C. difficile* spores. Wells coated with 1.6 × 10^7^ spores of strains 630Δ*ermB* or R20291, or with 1% BSA as a negative control, were incubated with increasing concentrations of purified human C3. Bound C3 was detected by immunoassay, and absorbance was measured at 492 nm. Apparent *K*d values calculated from the binding curves are indicated. Data represent the mean ± SEM of four independent wells from two independent experiments. **F,** Representative phase-contrast, anti-C3 immunofluorescence, and fluorescence-intensity surface plots of R20291 spores incubated with purified C3 or control buffer. Arrows indicate representative C3-associated spores. **G,** Quantification of C3-associated fluorescence intensity in individual R20291 spores incubated without C3 (Ctrl) or with 5 μg/mL purified C3. Each point represents an individual spore. Scale bars in **B** and **F**, 5 μm. Statistical significance in **A** and **E** was determined by two-way ANOVA with Sidak’s multiple-comparisons test, and in **C** and **G** by two-tailed unpaired Student’s *t*-test. ns, not significant; ****, *P* < 0.0001.

Purified C3 also exhibited concentration-dependent binding to spores of both strains in the solid-phase assay, with minimal signal detected for the BSA control (Fig. 4E). Immunofluorescence analysis revealed a markedly different pattern from that observed with C1q. Following incubation with purified C3, essentially all spores exhibited detectable C3-associated fluorescence (Fig. 4F,G). Quantification showed a pronounced increase in mean C3-associated fluorescence, from 100 in the control condition to approximately 2,6% of control following incubation with C3 (Fig. 4G). Notably, the distribution was highly heterogeneous, with a subset of spores exhibiting exceptionally high levels of C3-associated fluorescence, indicating substantial variation in the amount of C3 associated with individual spores.

Together, these results demonstrate that purified human C1q and C3 directly interact with *C. difficile* spores, but with distinct population-level patterns. C1q association was restricted to approximately half of the spore population, whereas C3 associated with essentially all spores and exhibited a broad range of fluorescence intensities. These findings suggest that C1q and C3 interact differently with the spore surface and prompted us to examine their ultrastructural localization on the exosporium.

### Purified human C1q and C3 localize to the exosporium of *C. difficile* spores

To define the ultrastructural localization of C1q and C3 on the spore surface, R20291 spores were incubated with purified human C1q or C3 and analyzed by transmission electron microscopy coupled with immunogold labeling. Immunogold particles specific for C1q were detected at the outermost region of the spore, in close proximity to the exosporium layer (Fig. 5A,B). Quantification showed that 32.5% of spores were C1q-associated, whereas 67.5% showed no detectable C1q-associated gold particles (Fig. 5C). Among the C1q-associated spores, 68% exhibited a thin exosporium and 32% a thick exosporium (Fig. 5D).

**Fig. 5.**
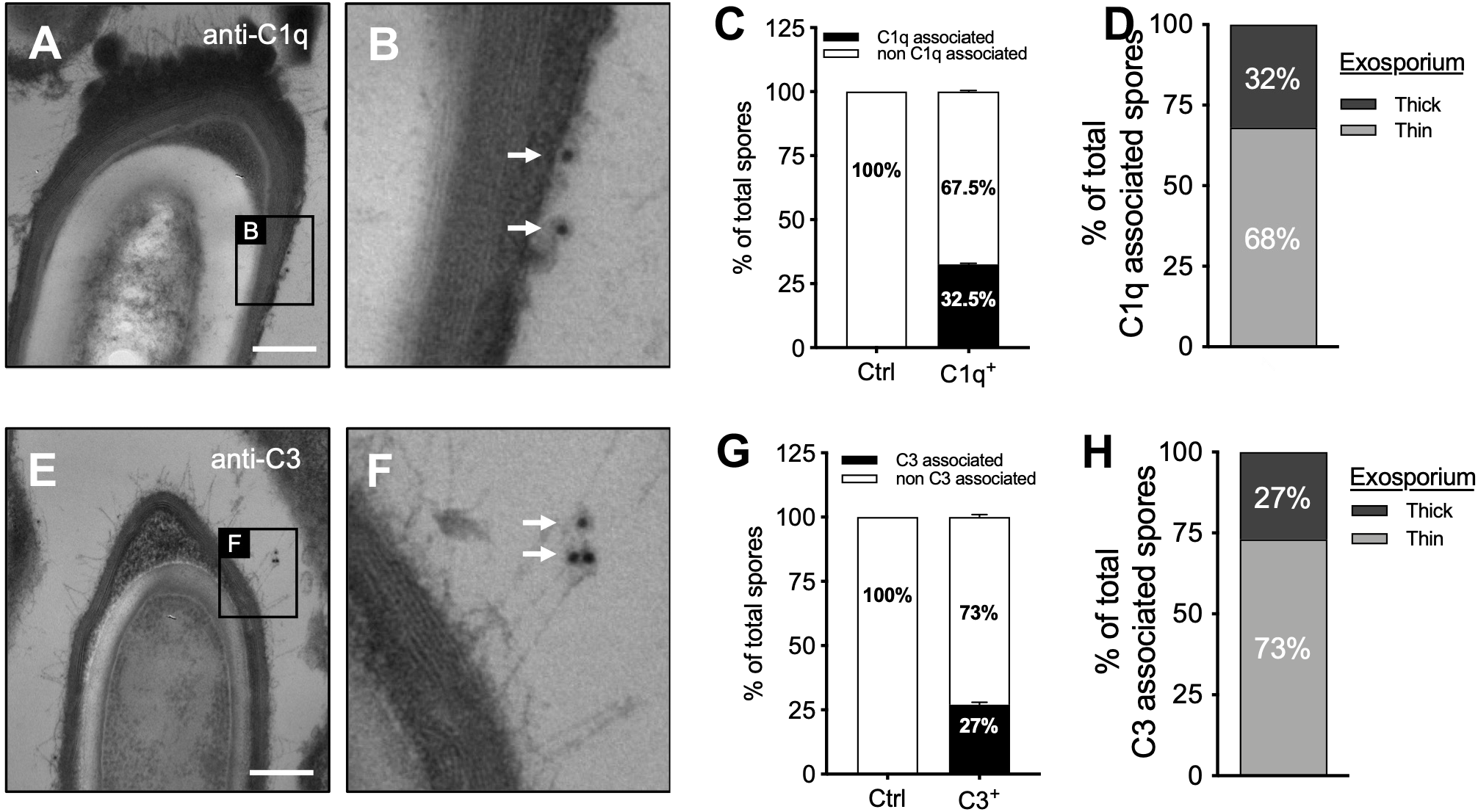
Purified human C1q and C3 associate with the exosporium of *C. difficile* spores. *C. difficile* R20291 spores were incubated with 10 μg/mL purified human C1q or C3 for 45 min at 37 °C and analyzed by transmission electron microscopy (TEM) coupled with immunogold labeling. **A,** Representative TEM image of an R20291 spore incubated with purified C1q and immunolabeled for C1q. The boxed region is shown at higher magnification in **B**. **B,** Higher-magnification image showing C1q-associated 12-nm immunogold particles at the outer spore surface. Arrows indicate representative immunogold particles. **C,** Distribution of spores classified as C1q-associated or non-C1q-associated. **D,** Distribution of exosporium morphotypes among C1q-associated spores; 68% exhibited a thin exosporium and 32% exhibited a thick exosporium. **E,** Representative TEM image of an R20291 spore incubated with purified C3 and immunolabeled for C3. The boxed region is shown at higher magnification in **F**. **F,** Higher-magnification image showing C3-associated 12-nm immunogold particles at the outer spore surface. Arrows indicate representative immunogold particles. **G,** Distribution of spores classified as C3-associated or non-C3-associated. **H,** Distribution of exosporium morphotypes among C3-associated spores. For **C** and **G**, 100 individual spores were analyzed from two independent experiments. Data are presented as mean ± SEM. Scale bars in **A** and **E**, 200 nm.

Similarly, C3-specific immunogold particles were detected at the outer spore surface, including regions associated with the hair-like projections of the exosporium (Fig. 5E,F). Quantification showed that 27% of spores were C3-associated, whereas 73% showed no detectable C3-associated gold particles (Fig. 5G). Of the C3-associated spores, 73% exhibited a thin exosporium and 27% a thick exosporium (Fig. 5H).

Together, these results provide ultrastructural evidence that purified human C1q and C3 associate with the outer exosporium of *C. difficile* spores. Both complement components were detected on spores exhibiting either thin or thick exosporium morphologies, indicating that their association is not restricted to a single exosporium morphotype.

### BclA proteins modulate, but are not required for, C1q and C3 binding to *C. difficile* spores

The exosporium glycoprotein BclA of *Bacillus anthracis* has been implicated in interactions between spores and the complement components C1q and C3 [34]. *C. difficile* encodes three BclA-like exosporium proteins, BclA1, BclA2, and BclA3, suggesting that these proteins could similarly contribute to complement association with the spore surface.

We therefore examined the contribution of the *C. difficile* BclA proteins to C1q and C3 binding using previously characterized BclA mutants in two strain backgrounds. The laboratory strain 630Δ*ermB*, which lacks the prominent hair-like exosporium extensions [5], was analyzed using previously described insertional inactivation mutants CT-*bclA1*, CT-*bclA2*, and CT-*bclA3* mutants [87], whereas the epidemic strain R20291, which exhibits prominent hair-like projections [44], was analyzed using a Δ*bclA3* deletion mutant and its complemented derivative.

Solid-phase binding assays were performed using spores from the 630Δ*ermB* background carrying C-terminal disruptions in *bclA1*, *bclA2*, or *bclA3*, as well as R20291 spores lacking *bclA3* and the corresponding complemented strain. In the 630Δ*ermB* background, purified C1q bound concentration-dependently to wild-type spores, whereas all three BclA mutant strains exhibited altered binding profiles (Fig. 6A). Differences became most apparent at the higher C1q concentrations, where CT-*bclA1*, CT-*bclA2*, and CT-*bclA3* spores exhibited significantly lower binding than wild-type spores. Nevertheless, all three mutants retained substantial C1q association. The apparent dissociation constants also differed among strains, with *K*d values of approximately 2.0 μM for wild-type spores, 2.2 μM for CT-*bclA1*, 4.1 μM for CT-*bclA2*, and 3.0 μM for CT-*bclA3*, indicating that disruption of individual BclA proteins modified but does not abolish C1q binding.

**Fig. 6.**
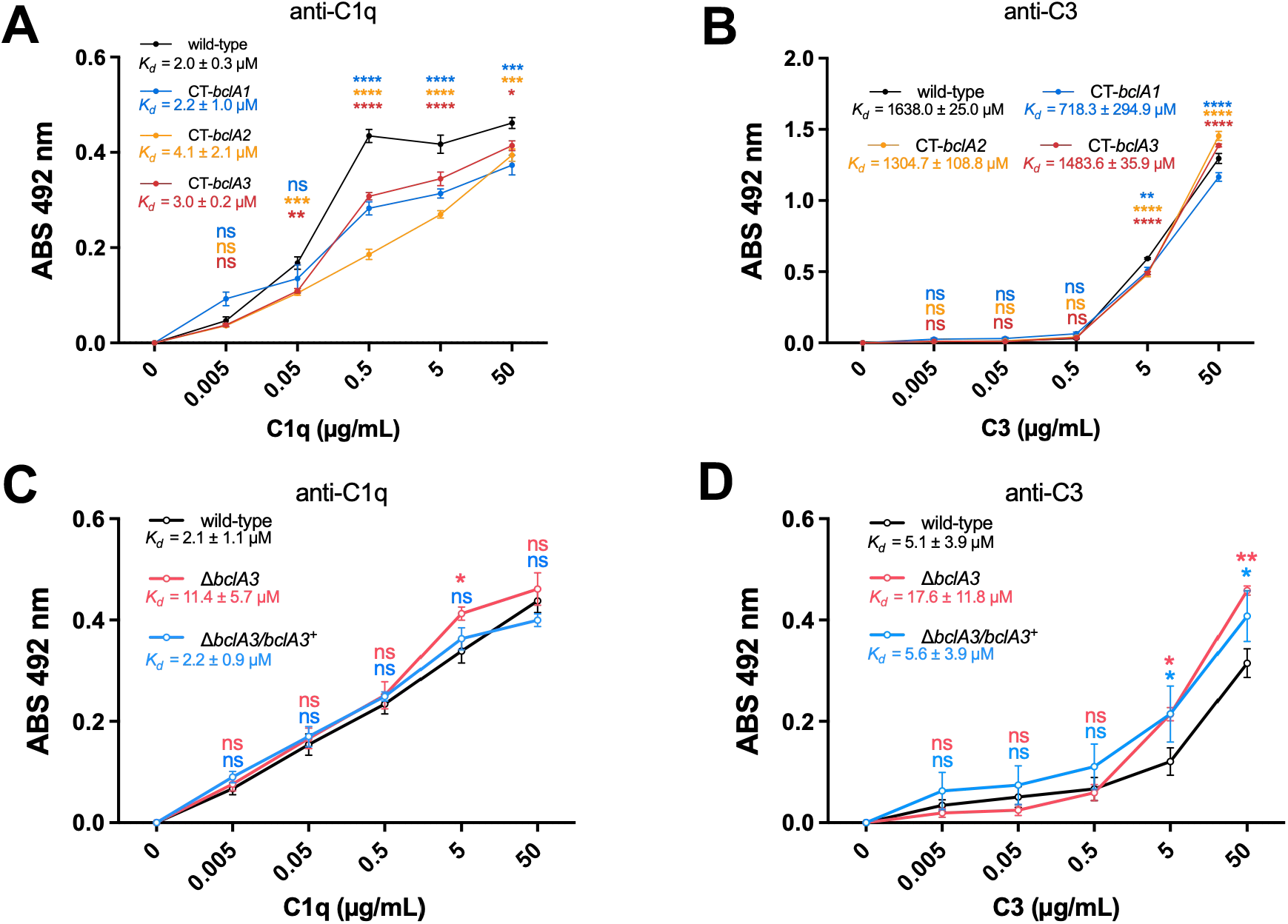
Purified human C1q and C3 bind to BclA mutant *C. difficile* spores. Solid-phase binding assays were performed to evaluate the contribution of the exosporium collagen-like BclA proteins to the interaction of purified human C1q and C3 with *C. difficile* spores. Wells coated with 1.6 × 10^7^ spores were incubated with increasing concentrations of purified human C1q or C3, washed, and bound complement protein was detected by immunoassay. **A, B,** Binding of **A,** C1q and **B,** C3 to spores of *C. difficile* strain 630Δ*ermB* and the C-terminal insertional mutants CT-*bclA1*, CT-*bclA2*, and CT-*bclA3*. **C, D,** Binding of **C,** C1q and **D,** C3 to spores of *C. difficile* R20291 Δ*pyrE*/ *pyrE*+ parental strain, the Δ*bclA3* mutant, and the complemented Δ*bclA3*/ *bclA3*+ strain. Apparent dissociation constants (*K*d) calculated from the corresponding binding curves are indicated in each panel. Data represent the mean ± SEM of six wells collected from three independent experiments. Statistical significance was determined by two-way ANOVA with Sidak’s multiple-comparisons test. Colored significance symbols indicate comparisons of the corresponding mutant or complemented strain with its respective parental strain. ns, not significant; *, *P* < 0.05; **, *P* < 0.01; ***, *P* < 0.001; ****, *P* < 0.0001.

C3 also associated concentration-dependently 630Δ*ermB* spores (Fig. 6B). Differences between wild-type and BclA mutant spores were most evident at the higher C3 concentrations, whereas little or no difference was observed at the lower concentrations. Despite these quantitative differences, all three BclA mutants retained C3 binding, indicating that none of the individual BclA proteins is essential for C3 association with the spore surface.

We next examined the contribution of BclA3 in the R20291 background, in which the hair-like exosporium projections are prominent [45, 46]. Deletion of *bclA3* had little effect on C1q binding across the concentration range tested (Fig. 6C). The Δ*bclA3* mutant and complemented strain exhibited binding profiles broadly comparable to wild-type R20291 spores, with only modest differences at individual concentrations. Thus, BclA3 is not required for C1q binding to R20291 spores.

Similarly, deletion of *bclA3* did not impair C3 binding to R20291 spores (Fig. 6D). Rather, at the higher C3 concentrations, both the Δ*bclA3* mutant and the complemented strain exhibited greater C3-associated signal than wild-type spores. Therefore, the absence of BclA3 does not reduce C3 binding and may alter the accessibility of other C3-interacting components on the spore surface. Together, these results indicate that BclA proteins can influence the magnitude of C1q and C3 binding in a strain-dependent manner but are not individually required for complement association. The persistence of substantial C1q and C3 binding in the BclA mutants suggested that additional spore-surface proteins contribute to these interactions and prompted us to identify other candidate complement-interacting spore proteins.

### Identification of C1q-and C3-interacting proteins in the *C. difficile* spore coat/exosporium

Because disruption of individual BclA proteins did not abolish C1q or C3 binding to *C. difficile* spores, we next sought to identify additional spore-surface proteins capable of interacting with these complement components. Coat/exosporium proteins extracted from R20291 spores were resolved by two-dimensional electrophoresis (2-DE). Parallel gels were either stained with Coomassie blue or transferred to nitrocellulose membranes and subjected to far-Western blotting using purified human C1q or C3 as bait (Fig. 7A). Far-Western analysis revealed several C1q-and C3-reactive spots within the coat/exosporium protein fraction. Four reproducibly detectable spots that corresponded to proteins visible in the 2-DE preparation were selected for identification by tandem mass spectrometry (Fig. 7A,B). Two complement-reactive proteins were identified: CotE (CDR20291_1282) and CdeM (CDR20291_1478), both previously localized to the outer layers of the *C. difficile* spore. CotE was identified in spot 1 with 54% sequence coverage and was detected in far-Western blots using both C1q and C3 as bait (Fig. 7A,B). CdeM was identified in three distinct spots. Spot 2, with 65% sequence coverage, reacted with both C1q and C3, whereas spots 3 and 4, with 10% and 49% sequence coverage, respectively, were detected with C3 but not C1q (Fig. 7A,B). The appearance of CdeM at multiple positions in the 2-DE analysis may reflect distinct molecular forms of the protein, although the basis for this heterogeneity was not investigated here. Together, these results identify CotE and CdeM as candidate C1q-and C3-interacting proteins within the *C. difficile* spore coat/exosporium. Notably, the repeated identification of CdeM in complement-reactive spots, including multiple C3-reactive species, prompted us to directly examine whether CdeM can associate with purified C1q and C3.

**Fig. 7.**
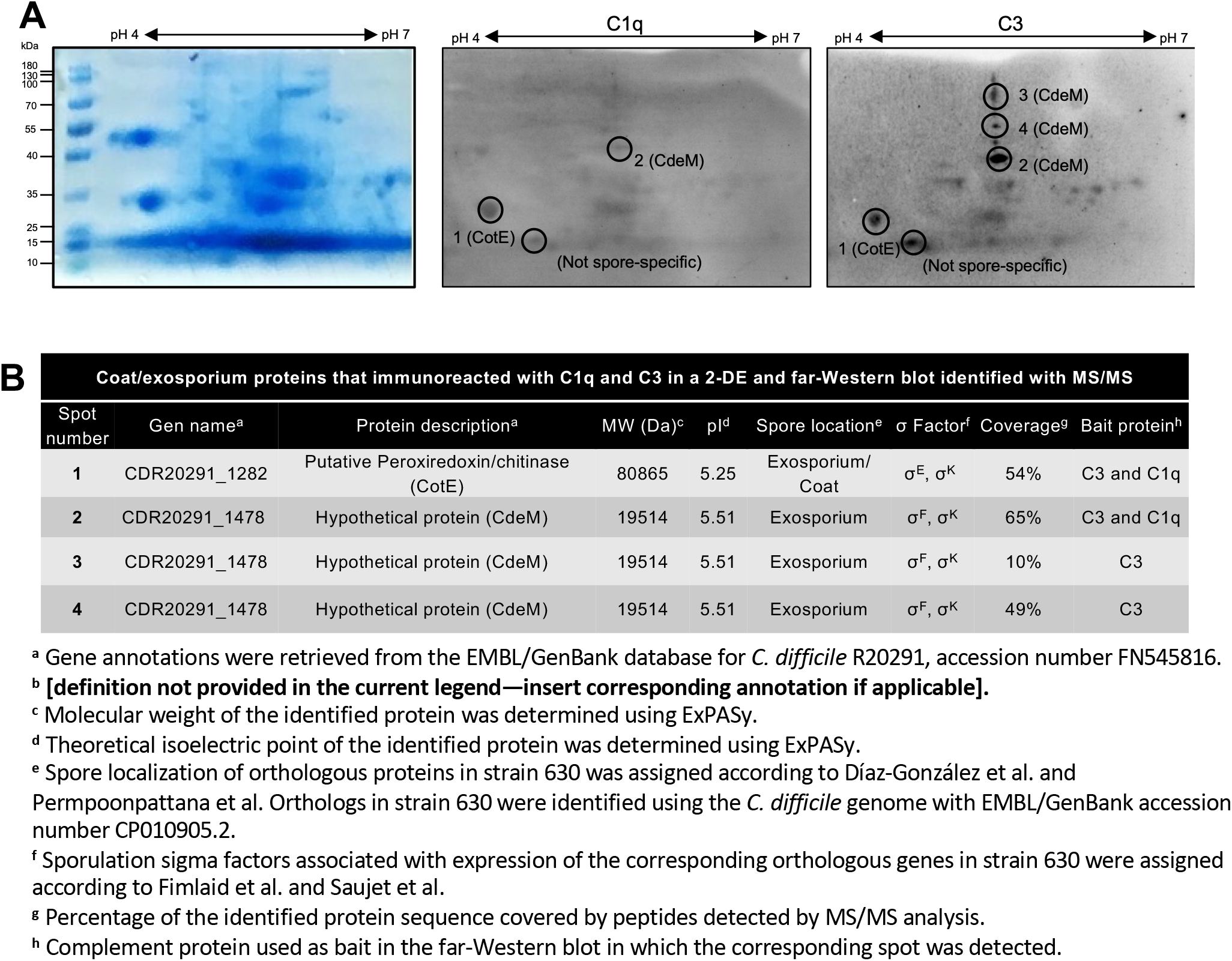
Identification of *C. difficile* spore coat/exosporium proteins associated with C1q and C3 by far-Western blotting and mass spectrometry. A, Coat/exosporium proteins extracted from *C. difficile* R20291 spores were separated by two-dimensional electrophoresis (2-DE), using isoelectric focusing over a pH 4–7 range in the first dimension followed by SDS-PAGE in the second dimension. A representative 2-DE gel was stained with Coomassie blue (left). Parallel gels were transferred to nitrocellulose membranes and subjected to far-Western blotting by incubation with 5 μg/mL purified human C1q (middle) or C3 (right), followed by immunodetection of the corresponding complement protein. Immunoreactive spots selected for protein identification are indicated by circles and numbered according to the identifications shown in **B**. Spots corresponding to CotE and CdeM are indicated. A recurrent immunoreactive spot that was not considered spore-specific is also indicated. **B,** Identification by tandem mass spectrometry (MS/MS) of proteins corresponding to selected C1q-and/or C3-reactive spots. Proteins were assigned by matching MS/MS-derived peptide sequences against the *C. difficile* R20291 protein database. The table summarizes the spot number, gene name, protein description, predicted molecular weight, theoretical isoelectric point, reported spore localization, sporulation sigma factor associated with gene expression, sequence coverage obtained by MS/MS, and the complement protein used as bait in the corresponding far-Western blot. CotE and CdeM were identified among the complement-reactive spots; CotE and one CdeM spot were detected with both C1q and C3, whereas additional CdeM spots were detected with C3.

### CdeM associates with purified human C1q and C3

Among the candidate complement-interacting proteins identified by far-Western blotting, CdeM was of particular interest because it is a cysteine-rich morphogenetic component of the *C. difficile* exosporium [47]. Because recombinant CdeM readily self-assembles into inclusion bodies (IBs) when overexpressed in *E. coli* [48], and soluble recombinant exosporium proteins are difficult to obtain [49], we used CdeM IBs as a surrogate to directly evaluate the interaction of CdeM with purified human C1q and C3.

Solid-phase binding assays demonstrated concentration-dependent association of purified C1q with both R20291 spores and CdeM IBs (Fig. 8A). CdeM IBs exhibited a lower apparent dissociation constant than intact spores (*K*d = 2.0 ± 0.9 μM and 13.8 ± 7.6 μM, respectively), consistent with a stronger apparent interaction of C1q with CdeM IBs under these assay conditions. Immunofluorescence microscopy independently confirmed C1q association with both intact spores and CdeM IBs (Fig. 8B). Quantification of individual particles showed a concentration-dependent increase in C1q-associated fluorescence on spores, with significant increases at the concentrations tested relative to the no-C1q control (Fig. 8C). CdeM IBs similarly exhibited increased C1q-associated fluorescence, with significant increases at 1 and 10 μg/mL, whereas 0.1 μg/mL did not differ significantly from control (Fig. 8D).

**Fig. 8.**
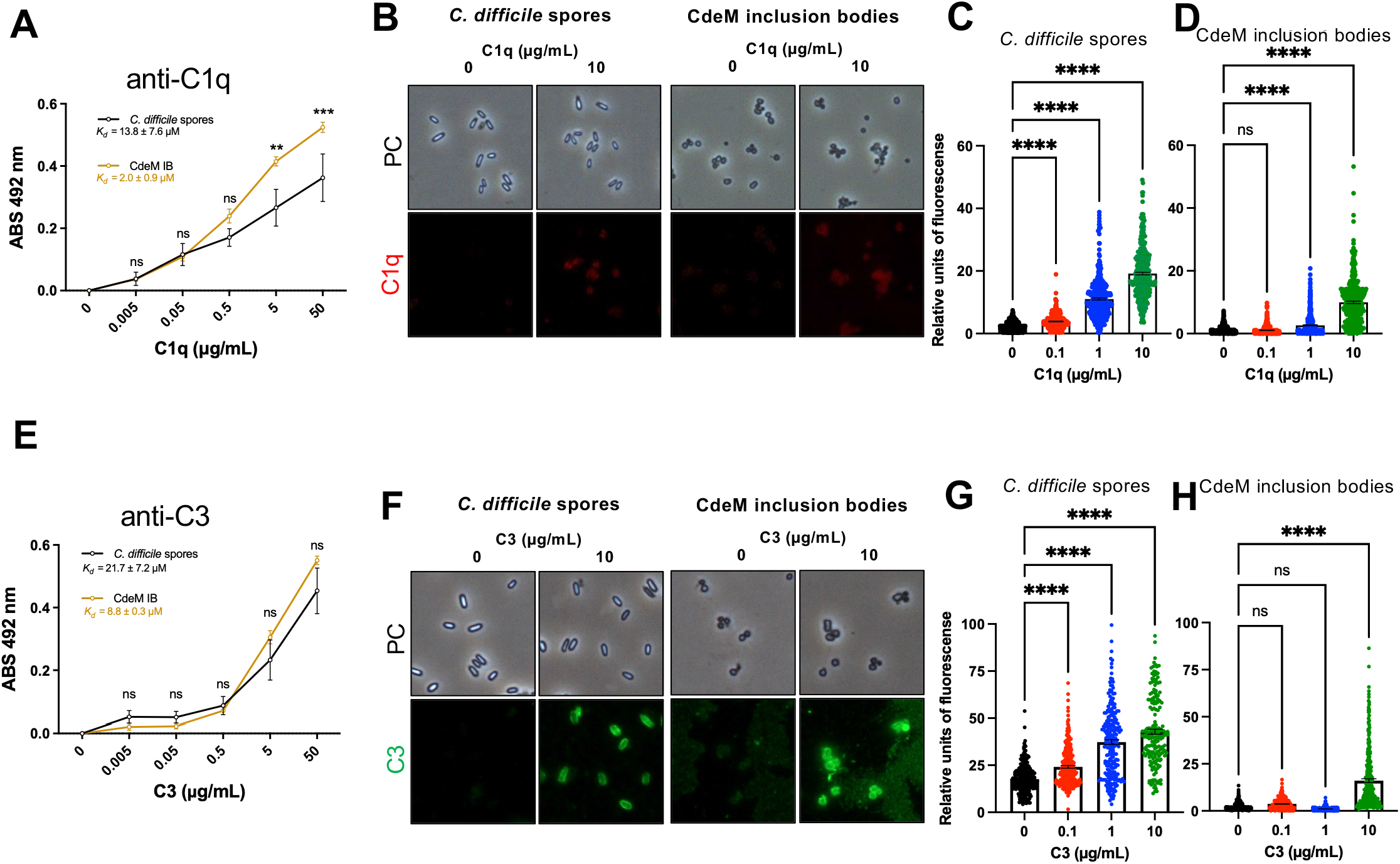
CdeM inclusion bodies associate with purified human C1q and C3. **A.** Solid-phase binding assay comparing the interaction of purified human C1q with *C. difficile* R20291 spores and CdeM inclusion bodies (IBs). Wells coated with *C. difficile* spores or CdeM IBs were incubated with increasing concentrations of purified human C1q, and associated C1q was detected by immunoassay. Apparent dissociation constants (*K*d) calculated from the binding curves are indicated. **B,** Representative phase-contrast (PC) and immunofluorescence images of *C. difficile* R20291 spores and CdeM IBs incubated in the absence or presence of 10 μg/mL purified human C1q. C1q-associated fluorescence is shown in red. **C, D,** Quantification of C1q-associated fluorescence intensity in individual **C,** *C. difficile* spores and **D,** CdeM IBs following incubation with 0, 0.1, 1, or 10 μg/mL purified C1q. Each point represents an individual spore or CdeM IB. **E,** Solid-phase binding assay comparing the interaction of purified human C3 with *C. difficile* R20291 spores and CdeM IBs. Wells coated with *C. difficile* spores or CdeM IBs were incubated with increasing concentrations of purified human C3, and associated C3 was detected by immunoassay. Apparent *K*d values calculated from the binding curves are indicated. **F,** Representative phase-contrast and immunofluorescence images of *C. difficile* R20291 spores and CdeM IBs incubated in the absence or presence of 10 μg/mL purified human C3. C3-associated fluorescence is shown in green. **G, H,** Quantification of C3-associated fluorescence intensity in individual **G,** *C. difficile* spores and **H,** CdeM IBs following incubation with 0, 0.1, 1, or 10 μg/mL purified C3. Each point represents an individual spore or CdeM IB. Solid-phase binding assays in **A** and **E** were performed in two independent experiments, and data are presented as mean ± SEM. Immunofluorescence experiments were independently performed twice; **B–D** and **F–H** show experiments performed using purified C1q or C3, respectively. Statistical significance in **A** and **E** was determined by two-way ANOVA with Sidak’s multiple-comparisons test. Statistical significance in **C, D, G,** and **H** was determined by one-way ANOVA with Dunnett’s multiple-comparisons test, comparing each complement concentration with the corresponding 0 μg/mL control. ns, not significant; *, *P* < 0.05; **, *P* < 0.01; ***, *P* < 0.001; ****, *P* < 0.0001.

Purified C3 also associated with both intact spores and CdeM IBs in the solid-phase binding assay (Fig. 8E), with apparent *K*d values of 21.7 ± 7.2 μM for spores and 8.8 ± 0.3 μM for CdeM IBs. Consistent with these findings, C3-associated fluorescence was readily detected on spores and CdeM IBs following incubation with purified C3 (Fig. 8F). C3-associated fluorescence increased significantly with increasing C3 concentration on intact spores (Fig. 8G). For CdeM IBs, C3-associated fluorescence was significantly increased at 10 μg/mL, whereas the lower concentrations did not differ significantly from the untreated control (Fig. 8H).

Together, these results validate CdeM as a C1q-and C3-interacting exosporium protein and support the far-Western/MS identification in Figure 7. The ability of isolated CdeM IBs to associate with both purified complement proteins indicates that CdeM itself can contribute to the interaction of the *C. difficile* spore surface with C1q and C3.

### CdeM inclusion bodies adhere to and internalize into intestinal epithelial cells

C1q and C3 have previously been implicated in the internalization of *Bacillus anthracis* spores by host cells [34, 35, 50]. Because CdeM was identified here as a C1q-and C3-interacting component of the *C. difficile* exosporium, we asked whether CdeM itself possesses properties that could contribute to spore interactions with intestinal epithelial cells. To address this question, CdeM inclusion bodies (IBs), which recapitulate the self-assembly properties of CdeM, were incubated with undifferentiated Caco-2 cells in the absence or presence of normal human serum (NHS). CdeM IBs efficiently adhered to Caco-2 cells in the absence of serum (Fig. 9A). Quantification showed an average of 7.6 ± 1.6 IBs per cell in DMEM, demonstrating that CdeM assemblies are sufficient to associate with intestinal epithelial cells (Fig. 9B). Pre-incubation with NHS significantly reduced adherence to 3.9 ± 1.1 IBs per cell, corresponding to an approximately 50% decrease relative to the serum-free condition (*P* < 0.01). We next asked whether CdeM IBs could gain entry into intestinal epithelial cells. Extracellular IBs were immunodetected before cell permeabilization, followed by detection of the total IB population after permeabilization, allowing internalized IBs to be distinguished from those remaining extracellular (Fig. 9C). In the absence of NHS, 8.3 ± 1.2% of cell-associated CdeM IBs were internalized, indicating that CdeM assemblies possess an intrinsic capacity to enter intestinal epithelial cells (Fig. 9D). Pre-incubation with NHS reduced internalization to 3.5 ± 0.6%, representing an approximately 58% decrease relative to the DMEM condition (*P* < 0.01). Because the epithelial surface is not uniform, we next determined where on the Caco-2 monolayer the cell-associated CdeM IBs localized. In both conditions the large majority of IBs were found at tight junctions rather than at the apical cell surface (91.3% versus 9.0% in NHS; 86.8% versus 11.3% in DMEM), and pre-incubation with NHS did not significantly alter this distribution (Fig. 9E). Thus, while NHS reduces the number of CdeM IBs that associate with intestinal epithelial cells, it does not change their preferred site of association, indicating that tight junction targeting is an intrinsic property of CdeM assemblies. Together, these results demonstrate that CdeM assemblies can both adhere to and enter intestinal epithelial cells and that both processes are reduced in the presence of NHS. Although these experiments do not establish that CdeM is required for internalization of intact *C. difficile* spores, they reveal intrinsic epithelial adherence and internalization properties of CdeM and provide a rationale to determine whether C1q and C3 similarly influence the interaction of intact *C. difficile* spores with intestinal epithelial cells.

**Fig. 9.**
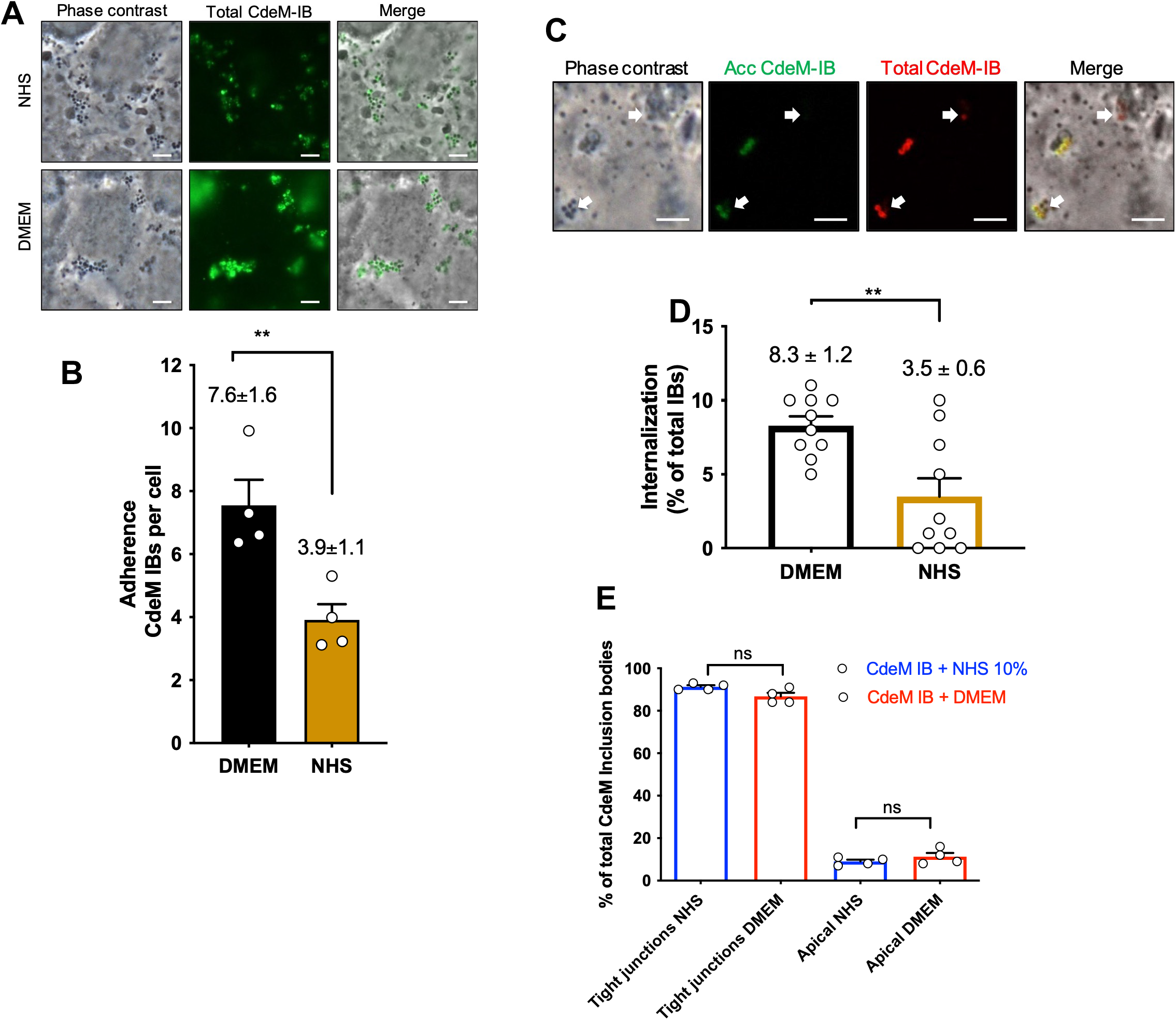
CdeM inclusion bodies adhere to and are internalized by intestinal epithelial cells. Undifferentiated Caco-2 cells were incubated for 5 h with CdeM inclusion bodies (IBs) at an MOI of 10 following pre-incubation of the IBs with either DMEM or 10% normal human serum (NHS). **A**, Representative epifluorescence micrographs of Caco-2 cells incubated with CdeM IBs pre-treated with DMEM or NHS. Total CdeM IBs were immunodetected in green. Phase-contrast, fluorescence, and merged images are shown. **B**, Quantification of CdeM IB adherence to Caco-2 cells, expressed as the number of cell-associated CdeM IBs per cell. Adherence was higher following pre-incubation in DMEM (7.6 ± 1.6 IBs per cell) than following pre-incubation with NHS (3.9 ± 1.1 IBs per cell). **C**, Representative epifluorescence micrographs illustrating CdeM IB internalization by Caco-2 cells. In non-permeabilized cells, accessible CdeM IBs (acc CdeM-IB) were immunodetected in green; cells were subsequently permeabilized and total CdeM IBs were immunodetected in red. CdeM IBs detectable by phase contrast and total staining but lacking accessible staining were classified as internalized. Arrows indicate representative CdeM IBs. **D**, Quantification of CdeM IB internalization, expressed as the percentage of total cell-associated IBs. Internalization was higher following pre-incubation in DMEM (8.3 ± 1.2%) than following pre-incubation with NHS (3.5 ± 0.6%). **E**, Distribution of cell-associated CdeM IBs on the Caco-2 monolayer, expressed as the percentage of total CdeM IBs localized at tight junctions or at the apical cell surface, following pre-incubation with 10% NHS (blue) or DMEM (red). In both conditions the majority of CdeM IBs localized at tight junctions (91.3 ± [SEM]% NHS; 86.8 ± [SEM]% DMEM) rather than at the apical surface (9.0 ± [SEM]% NHS; 11.3 ± [SEM]% DMEM), and pre-incubation with NHS did not alter this distribution. Statistical significance in E was determined by ordinary one-way ANOVA with Tukey’s multiple comparisons test; ns, not significant. Data in B, D, and E represent the mean ± SEM from three independent experiments. Statistical significance in B and D was determined using a two-tailed unpaired Student’s t-test. **, P < 0.01. Scale bars: A, 5 μm; C, 2.5 μm.

### C1q and C3 contribute to the internalization of *C. difficile* spores by intestinal epithelial cells

Because C1q and C3 associate with the *C. difficile* spore surface and have previously been implicated in the internalization of *Bacillus anthracis* spores by host cells [34, 35, 50], we next asked whether these complement components influence the interaction of intact *C. difficile* spores with intestinal epithelial cells. R20291 spores were pre-incubated for 45 min at 37°C with normal human serum (NHS), serum-free medium (SFM), C1q-depleted serum (DC1q), C3-depleted serum (DC3), or the corresponding depleted sera reconstituted with purified C1q or C3. Prior to infection, spores were washed to remove unbound complement, and Caco-2 cells were washed to remove complement components present in the culture serum. Spores were then incubated with Caco-2 intestinal epithelial cells for 4 h.

Extracellular spores were detected by immunofluorescence, whereas total cell-associated spores were identified by phase-contrast microscopy, allowing spore adherence and internalization to be quantified (Fig. 10A).

**Fig. 10.**
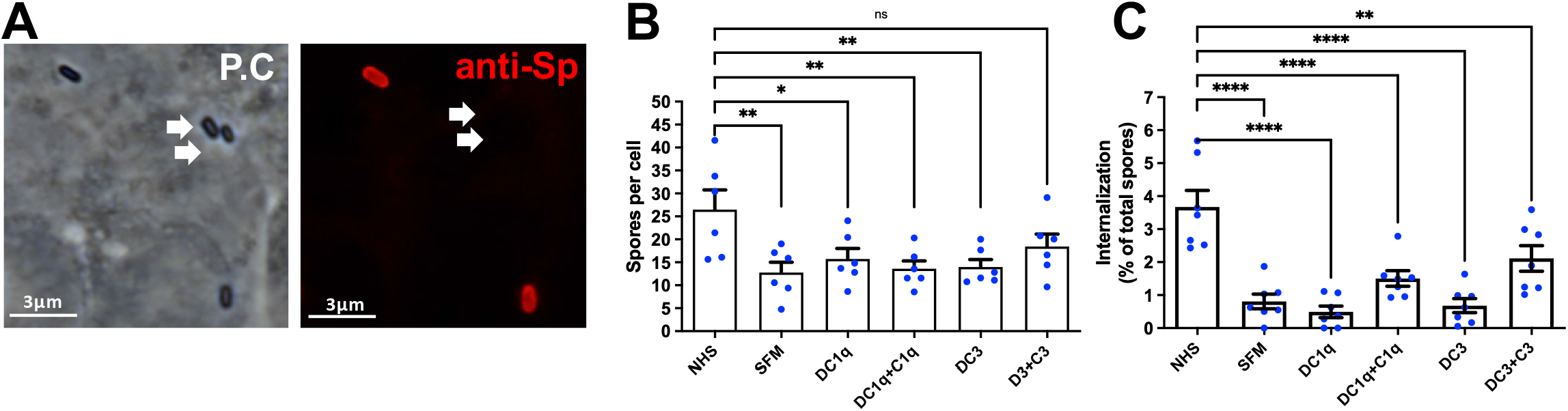
Adherence and internalization of *C. difficile* spores following exposure to C1q-and C3-containing sera. *C. difficile* R20291 spores were pre-incubated with normal human serum (NHS), serum-free medium (SFM), C1q-depleted serum (DC1q), C1q-depleted serum reconstituted with purified C1q (DC1q+C1q), C3-depleted serum (DC3), or C3-depleted serum reconstituted with purified C3 (DC3+C3), washed, and subsequently incubated with intestinal epithelial cells. **A,** Representative phase-contrast and anti-spore immunofluorescence images used to distinguish extracellular from internalized spores. Extracellular spores were immunodetected before permeabilization, whereas total spores were identified by phase-contrast microscopy. Arrows indicate representative spores. **B,** Quantification of spore adherence, expressed as the number of cell-associated spores per cell under each pre-incubation condition. **C,** Quantification of spore internalization, expressed as the percentage of total cell-associated spores under each pre-incubation condition. Individual points represent independent samples, and bars indicate the mean ± SEM. Statistical significance in **B** and **C** was determined by one-way ANOVA followed by Sidak’s multiple-comparisons test. ns, not significant; *, *P* < 0.05; **, *P* < 0.01; ****, *P* < 0.0001. Scale bars, 3 μm.

Pre-incubation with NHS markedly increased the number of spores associated with Caco-2 cells, reaching 26.5 ± 4.3 spores per cell, compared with 12.8 ± 2.2 spores per cell following incubation in SFM (Fig. 10B). Spore association was also reduced following incubation with C1q-or C3-depleted serum, reaching 15.7 ± 2.3 and 14.0 ± 1.6 spores per cell, respectively. Reconstitution of C3-depleted serum with purified C3 increased spore association to 18.4 ± 2.7 spores per cell, a value that was not significantly different from the NHS condition. In contrast, reconstitution of C1q-depleted serum with purified C1q resulted in 13.6 ± 1.7 spores per cell and did not restore the adherence phenotype toward that observed with NHS. These results indicate that serum enhances the association of *C. difficile* spores with intestinal epithelial cells, with C3 contributing more clearly than C1q to this adherence phenotype under the conditions tested.

A more pronounced effect was observed for spore internalization. Spores pre-incubated with NHS exhibited 3.67 ± 0.50% internalization, whereas internalization decreased to 0.81 ± 0.22% following incubation in SFM (Fig. 10C). Depletion of C1q or C3 similarly reduced internalization to 0.49 ± 0.17% and 0.68 ± 0.21%, respectively. Reconstitution of C1q-depleted serum with purified C1q increased internalization to 1.51 ± 0.24%, whereas reconstitution of C3-depleted serum with purified C3 increased internalization to 2.11 ± 0.39%. Although neither reconstituted condition fully reproduced the internalization observed with NHS, addition of the corresponding purified complement component shifted spore entry toward the NHS phenotype.

Together, these findings indicate that serum-associated factors enhance both the adherence and internalization of *C. difficile* spores by intestinal epithelial cells. In particular, depletion of either C1q or C3 markedly reduced spore internalization, whereas addition of the corresponding purified protein increased entry toward the levels observed with NHS. These data support a model in which association of C1q and C3 with the *C. difficile* spore surface promotes spore internalization by Caco-2 intestinal epithelial cells.

### C3 is deposited on *C. difficile* spores without compromising spore viability

Because complement activation can culminate in assembly of the membrane attack complex on susceptible bacterial surfaces, we next asked whether C3 association was specific to the *C. difficile* spore morphotype and whether exposure to complement affected spore survival. We first compared C3 association with spores and vegetative cells of strains 630Δ*ermB* and R20291 following incubation with normal human serum (NHS), C3-depleted serum (C3-Dpl), or serum-free medium (SFM).

Immunoblot analysis revealed C3-derived species associated with spores of both 630Δ*ermB* and R20291 following incubation with NHS (Fig. 11A). In both strains, prominent immunoreactive bands corresponding to the C3 β chain (∼75 kDa) and C3dg (∼40 kDa) were detected in the spore-associated fraction. In contrast, these C3-derived species were not detected in spores incubated with SFM or C3-Dpl serum. Notably, no comparable C3-associated signal was detected in vegetative cells of either strain following incubation with NHS. These results indicate that C3 deposition is preferentially associated with the spore surface and that deposited C3 undergoes proteolytic processing, as evidenced by the presence of the C3dg fragment.

**Fig. 11.**
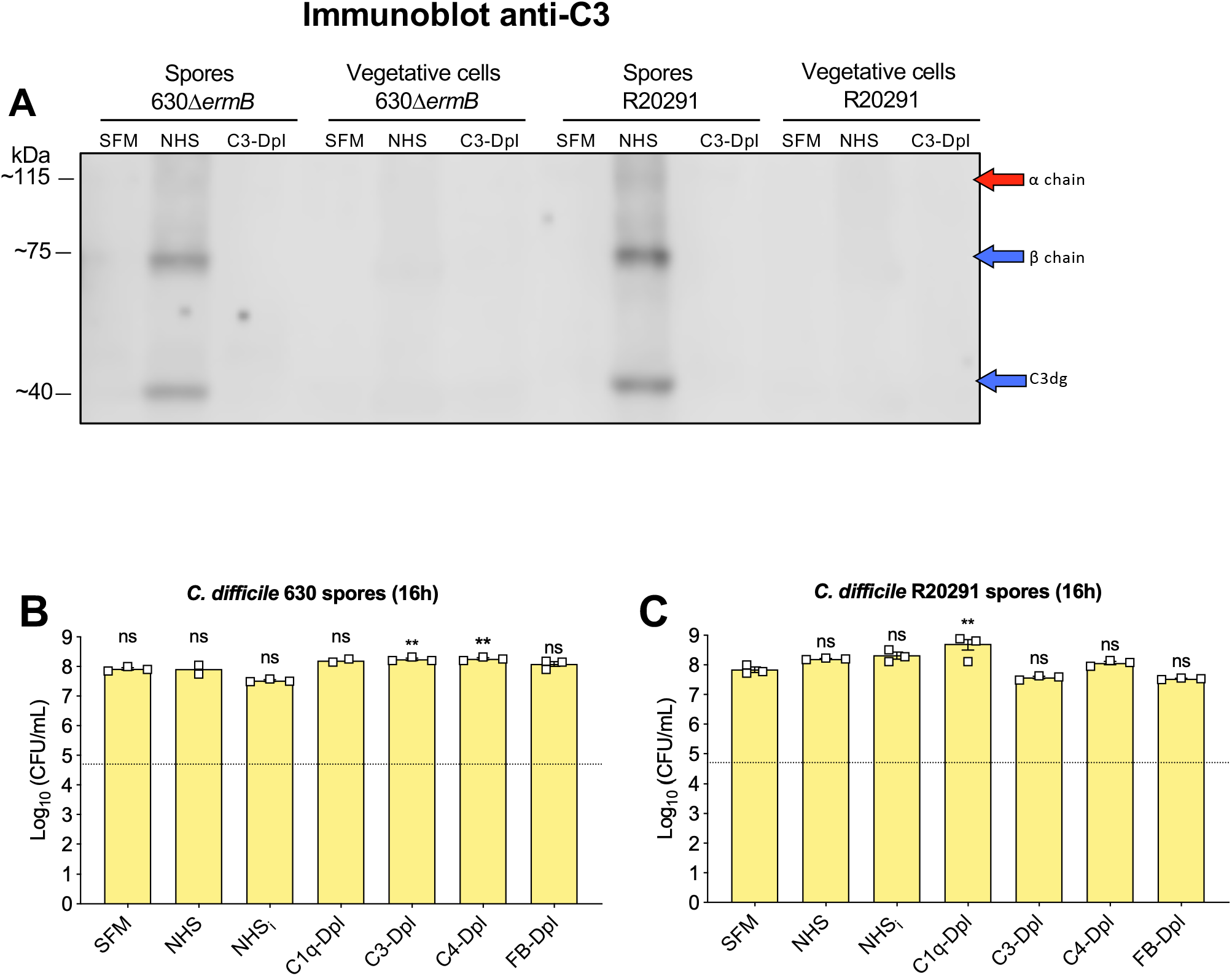
C3 association with *C. difficile* spores and vegetative cells, and effect of complement exposure on spore viability. **A,** Immunoblot analysis of C3 associated with spores or vegetative cells of *C. difficile* strains 630Δ*ermB* and R20291. Spores or vegetative cells were incubated with 10% normal human serum (NHS), 10% C3-depleted human serum (C3-Dpl), or serum-free medium (SFM), washed, and analyzed by immunoblotting with anti-C3 antibodies. Purified human C3 and NHS were included as positive controls. The positions of C3-derived species corresponding to the α chain (∼115 kDa), β chain (∼75 kDa), and C3dg (∼40 kDa) are indicated. **B, C,** Effect of complement exposure on the viability of *C. difficile* spores. Spores of **B,** strain 630Δ*ermB* and **C,** strain R20291 were incubated for 16 h with SFM, 10% NHS, heat-inactivated NHS (NHSi), C1q-depleted serum (C1q-Dpl), C3-depleted serum (C3-Dpl), C4-depleted serum (C4-Dpl), or factor B-depleted serum (FB-Dpl). Spore viability was determined by colony-forming unit (CFU) enumeration and expressed as log10 CFU/mL. Individual symbols in **B** and **C** represent independent samples, and bars indicate the mean ± SEM. The horizontal dotted line indicates the limit of detection. Statistical significance in **B** and **C** was determined separately by one-way ANOVA followed by Sidak’s multiple-comparisons test. ns, not significant; **, *P* < 0.01.

**Fig. 12.**
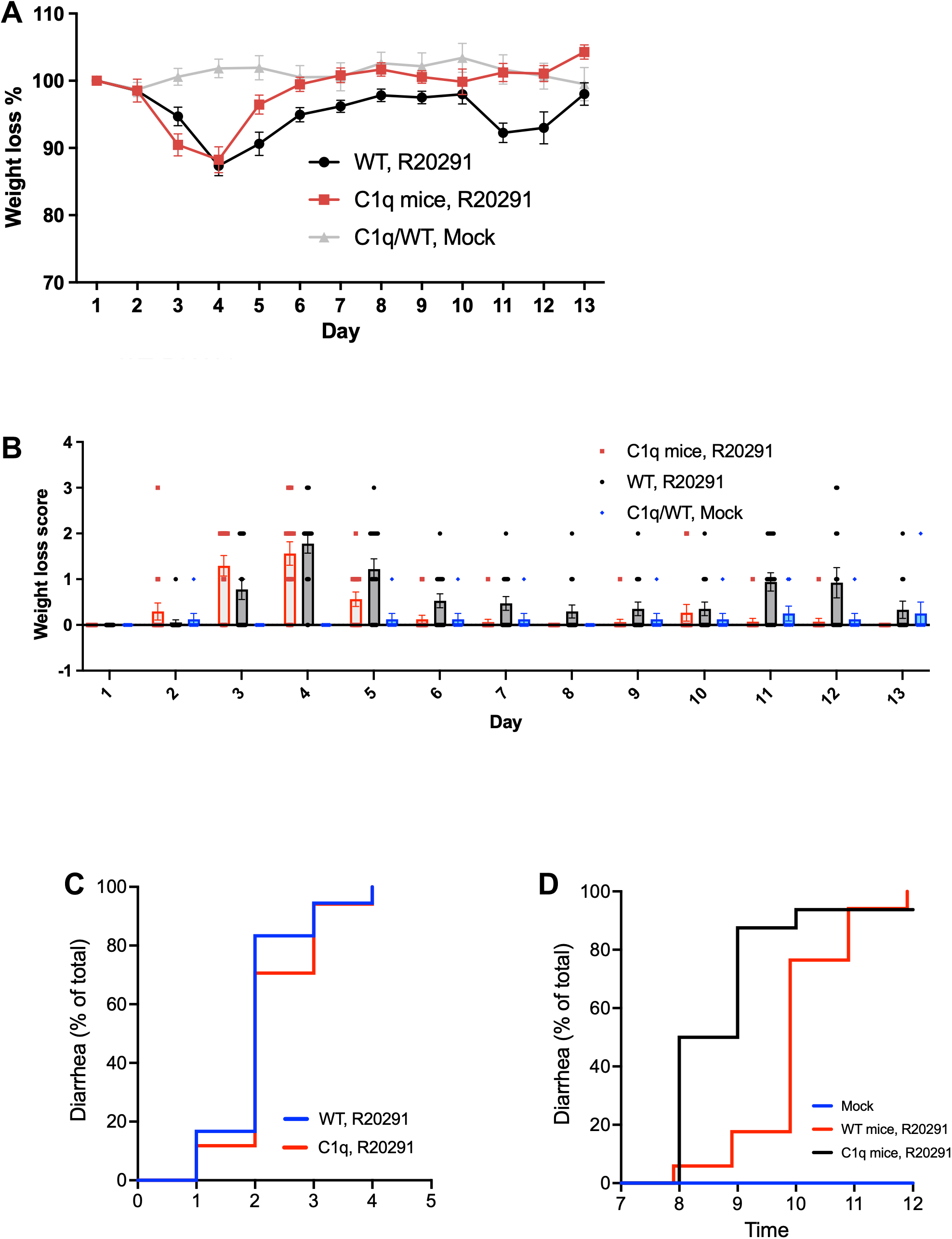
Clinical course of *C. difficile* infection in wild-type and C1q-deficient mice. Wild-type (WT) and C1q-deficient mice were infected with *Clostridioides difficile* strain R20291 and monitored for 13 days. Mock-treated WT and C1q-deficient mice were included as uninfected controls. **A,** Changes in body weight during the experimental period, expressed as percentage of initial body weight. **B,** Daily weight-loss clinical score for R20291-infected WT mice, R20291-infected C1q-deficient mice, and mock-treated controls. **C,** Cumulative percentage of R20291-infected WT and C1q-deficient mice that developed diarrhea during the early phase of infection. **D,** Cumulative percentage of mice exhibiting diarrhea during the later phase of the experiment in mock-treated controls and R20291-infected WT and C1q-deficient mice. Data represent **n = 9 mice per group from two independent experiments**. Error bars in **A** and **B** indicate the mean ± SEM.

Given the selective deposition of C3 on spores, we next determined whether complement exposure affected spore viability. Spores of 630Δ*ermB* and R20291 were incubated for 16 h with NHS, heat-inactivated NHS (NHSi), C1q-, C3-, C4-, or factor B-depleted serum, or SFM, and viable spores were quantified by CFU enumeration (Fig. 11B,C). For 630Δ*ermB*, none of the serum conditions produced a reduction in viable spore counts relative to the control conditions (Fig. 11B). Rather, modest increases were observed in some depleted-serum conditions. Similarly, R20291 spores remained fully viable following exposure to NHS or the different complement-depleted sera, with no condition producing a significant loss of viability (Fig. 11C). A modest increase in viable counts was observed under the C1q-depleted condition, but this did not indicate complement-mediated killing.

Together, these findings demonstrate that C3 selectively associates with and is processed on the surface of *C. difficile* spores rather than vegetative cells, yet complement exposure does not reduce spore viability. Thus, although spores provide a surface for C3 deposition and processing, they appear resistant to the bactericidal consequences of complement activation, consistent with their role as a highly resistant form involved in *C. difficile* persistence and transmission.

### C1q deficiency alters the clinical course of *C. difficile* infection in vivo

Because C1q associated with *C. difficile* spores and contributed to spore internalization by intestinal epithelial cells, we next asked whether C1q influences the course of CDI in vivo. Wild-type and C1q-deficient mice were infected with *C. difficile* R20291 and monitored for 13 days for changes in body weight and development of diarrhea. Mock-treated animals were included as uninfected controls (Fig. 12).

Both infected groups developed acute weight loss during the first days after infection, reaching their lowest body weights at approximately day 4 (Fig. 12A). C1q-deficient mice exhibited a pronounced early weight loss, similar in magnitude to that observed in infected wild-type mice, but recovered more rapidly thereafter. By days 6–8, C1q-deficient mice had returned to approximately their initial body weight and remained near or above baseline for the remainder of the experiment. In contrast, wild-type mice recovered more gradually and exhibited a second decrease in body weight around days 11–12 before recovering by day 13. Mock-treated animals maintained stable body weight throughout the observation period.

The clinical weight-loss scores reflected a similar pattern (Fig. 12B). Both infected groups exhibited their highest scores during the acute phase of disease, particularly around days 3–5. Scores subsequently declined in C1q-deficient mice and remained low during the later phase of the experiment. Wild-type mice, however, exhibited additional increases in clinical scores during the later observation period, coinciding with the second decrease in body weight.

The majority of mice in both infected groups developed diarrhea during the early phase of infection (Fig. 12C). By approximately day 2, diarrhea had developed in ∼83% of wild-type mice and ∼70% of C1q-deficient mice, with essentially all animals in both groups affected by days 3–4. Thus, C1q deficiency did not prevent the development of acute diarrhea, although the cumulative incidence appeared modestly delayed relative to wild-type mice.

A striking difference emerged during the later phase of infection (Fig. 12D). C1q-deficient mice developed late-onset diarrhea earlier than wild-type mice, with approximately 50% of C1q-deficient animals affected by day 8 and nearly 90% by day 9. In contrast, only a small fraction of wild-type mice exhibited diarrhea at day 8, with the cumulative incidence increasing primarily between days 9 and 11. Mock-treated animals did not develop diarrhea. Thus, although C1q-deficient mice recovered body weight more rapidly after the acute phase, they exhibited an earlier onset of diarrhea during the later phase of infection. Together, these findings indicate that C1q influences the temporal course of CDI rather than simply determining the severity of the initial acute episode. C1q deficiency was associated with rapid recovery of body weight after the initial disease phase but with earlier development of a subsequent diarrheal episode, suggesting that C1q may contribute to host processes that influence disease resolution and/or protection against later disease manifestations.

## Discussion

The high rate of recurrence of *Clostridioides difficile* infection (CDI) remains a major unresolved clinical challenge [51–53]. During the onset of infection, *C. difficile* produces the two major toxins TcdA and TcdB, which induce extensive epithelial injury and mucosal inflammatory response characterized by the production of pro-inflammatory cytokines and neutrophil-attracting chemokines, including CXCL1 and CXCL2, leading to neutrophil recruitment to the intestinal mucosa [54]. The resulting toxin-mediated disruption of the epithelial barrier is sought to create an environment suitable for *C. difficile* colonization and persistence of spore in the intestinal mucosa [23, 55, 56]. In previous work, we demonstrated that TcdA-and TcdB-mediated disruption of epithelial junctions increases the accessibility of E-cadherin and enhances *C. difficile* spore adherence to intestinal epithelial cells [19, 23]. We further demonstrated that *C. difficile* spores can enter the intestinal epithelium and persist intracellularly, and that pharmacological inhibition of spore entry during vancomycin treatment reduces recurrent CDI, supporting a role for spore–mucosa interactions in disease persistence and recurrence [19]. Hence, we sought to expand the repertoire of host factors that could contribute to interactions between *C. difficile* and the intestinal mucosa. In this context, gut complement system has been a underexplored host determinant. In this work, we identified discrete cellular types containing accessible and total C1q and C3 within the intestinal mucosa and found that TcdB intoxication markedly remodeled their distribution, with increased accessible C1q and C3 despite maintenance or decrease of total C1q and C3. We further demonstrated that *C. difficile* spores associate with accessible C1q and C3 *in vivo* in the healthy intestinal epithelium. We identified that the cysteine-rich exosporium morphogenetic protein, CdeM, acts as a spore-ligand for C1q-and C3, whereas the collagen-like proteins BclA1, BclA2, and BclA3 modulated but were not required for these interactions. Interestingly, our results demonstrate that an intact complement cascade promotes the entry of *C. difficile* spores into intestinal epithelial cells. Although C3 associated with *C. difficile* spores but not vegetative cells under the conditions tested, complement exposure did not reduce spore viability. Importantly, using C1q-deficient mice, we found that the absence of C1q did not prevent the onset of CDI but substantially altered the subsequent course of disease, including more rapid recovery from the acute clinical phenotype and an altered pattern of later disease manifestations.

The first contribution of this work is the demonstration that gut C1q and C3 are present and accessible within the healthy intestinal mucosa. Although circulating complement proteins are produced predominantly by the liver, increasing evidence demonstrates substantial extrahepatic production of complement components by both immune and nonimmune cells (ref). In the intestine, C3 has been shown to be produced locally, and recent studies under homeostatic conditions identified the subepithelial compartment as the predominant source of intestinal C3, with stromal cells representing the major C3-expressing population, followed by myeloid and epithelial cells. In agreement with these observations, our confocal imaging revealed a heterogeneous distribution of C3 within the ileal mucosa, with several distinct spatial patterns comprising both accessible and non-accessible C3 pools [42]. These findings complement previous cell-type analyses by demonstrating that intestinal C3 is not only produced by multiple cellular compartments but is also differentially accessible at the mucosal interface. Further studies using lineage-specific markers will be required to define the cellular identities associated with these distinct C3 distribution patterns. By contrast, less is known about the cellular distribution and accessibility of C1q in the intestinal mucosa; we observed a heterogeneous pattern for C1q, with at least three recurrent spatial distributions comprising accessible and non-accessible C1q pools, suggesting that C1q, like C3, is distributed among distinct cellular or tissue compartments within the healthy ileum. Importantly, TcdB intoxication markedly remodeled this intestinal complement landscape. Accessible C1q decreased despite preservation or accumulation of the total C1q pool, whereas C3 exhibited a different response, with increased accessibility at higher TcdB concentrations despite an overall reduction in total C3, changing the spatial availability of gut complement at mucosal surface. Our observation of increased C3 accessibility following TcdB intoxication is consistent with recent findings in *Citrobacter rodentium* infection in mice, in which luminal C3 levels increased significantly during infection together with increased C3 expression by stromal, myeloid, and epithelial populations [42]. Notably, Wu, et al demonstrated that C3-deficient mice exhibited increased susceptibility to *C. rodentium* [42], suggesting that locally produced mucosal C3 may play a role in host defense against other enteric pathogens such as *C. difficil*. Together, these findings suggest that intestinal infection and epithelial injury dynamically remodel the availability of complement at the luminal interface and that, during CDI, C1q and C3 may become differentially accessible to luminal *C. difficile*. However, whether these locally available complement components can influence *C. difficile* interaction with the intestinal mucosa remains largely unexplored.

Another major finding of this work is that C1q influences recovery following CDI and the severity of recurrent disease after vancomycin treatment. To our knowledge, a role for gut C1q in the clinical outcome of CDI and recurrence has not been previously reported. Our finding contrasts with the protective role described for luminal C3 during *Citrobacter rodentium* infection and suggests that components of the intestinal complement system exert different effects on enteric pathogen [42]. In our work, the absence of C1q did not alter the onset or initial severity of CDI, as wild-type and C1q-deficient mice exhibited comparable acute weight loss and diarrheal disease during the first days of infection. The main difference emerged during recovery and post-vancomycin treatment when C1q-deficient animals recovered body weight more rapidly and exhibited only minimal recurrent weight loss and negligible recurrent diarrhea compared with wild-type mice. These observations suggest that intestinal C1q may delay recovery and promote the severity of recurrent CDI. A plausible hypothesis is that C1q-dependent inflammatory signaling sustains mucosal injury or delays epithelial recovery following the acute inflammatory phase of the disease. Alternatively, C1q may influence the ability of *C. difficile* to establish a colonization that sustains subsequent persistence during vancomycin treatment and subsequent recolonization of the intestinal mucosa after vancomycin. Indeed, prior work suggests that C1q interacts with vegetative cells and may contribute to early colonization [38]. Moreover, toxin-dependent remodeling of intestinal C1q observed in our study further supports a potential role for C1q as part of a broader host response during epithelial injury.

However, a limitation of the present work is that we did not quantify intestinal *C. difficile* burden, spore persistence, or intestinal tissue-associated colonization in C1q-deficient and wild-type mice. Therefore, whether the attenuated recurrence observed in the absence of C1q is due to reduced inflammatory injury, enhanced epithelial recovery, decreased bacterial persistence, or a combination of these traits, remains unclear. Overall, these findings define, for the first time, a role for C1q in CDI pathogenesis and recurrent disease and warrant further work into how intestinal complement shapes *C. difficile* persistence and disease outcome.

A third major finding of this work is that *C. difficile* spores associate with C1q and C3 at the intestinal mucosal surface. We observed that spores associate with both C1q and C3 *in vivo* within the healthy ileal mucosa, demonstrating that complement–spore interactions can occur even in the absence of toxin-mediated epithelial injury. Although we did not test whether this association increases during *C. difficile* toxin-mediated intoxication of the intestinal mucosa, we observed that purified C1q and C3 directly interacted with *C. difficile* spores *in vitro* and localized predominantly to the outer exosporium. Indeed, the fac that both, C1q and C3, bind in a concentration dependent manner with high affinity Kd’s of ∼14 and 22, respectively, suggests that increased intestinal C1q and C3 will increase the extent of *C. difficile* spores associated with both molecules.

C1q and C3 bind differently to bacterial surfaces, while C1q commonly binds to a surface ligand first, followed by deposition of activated C3 fragments on the bacterial surface [57]. C1q and C3 ligands typically include bacterial surfaces, LPS, collagen-like proteins and surface protein adhesins [34, 57–61]. Here, we demonstrate that unlike with *B. anthracis* spores, where the collagen-like BclA protein mediates C1q and C3 deposition [34, 50], *C. difficile*’s collagen-like BclA1, BclA2 and BclA3 have no role in C1q and C3 binding. By contrast, the exosporium morphogenetic cysteine-rich protein, CdeM, seems to act as a C1q-and C3-spore ligand, as the affinity binding Kd was found to be ∼5 times lower than *C. difficile* spores. *C. difficile* strain deficient in CdeM has been previously described [47]. Inactivation of CdeM leads to absence of the entire exosporium layer, affecting abundance of BclA1, BclA2, BclA3, CdeC and coat proteins, including CotA and CotB. Give its defective surface, we avoided using this strain as a reagent to assess how absence of CdeM impacts C1q and C3 binding to *C. difficile* spores. An alternative approach that we are currently working is to use peptide microarray to map the sites where C1q and C3 bind to CdeM, as we recently described for E-cadherin [22]. Our far-western blots also suggested that CotE, a bifunctional spore coat enzyme that mediates intestinal adherence and colonization [62, 63]. CotE is a ∼81 kDa spore coat protein that possesses dual enzymatic activities (i.e., amino-terminal peroxiredoxin and a carboxy-terminal chitinase domain) [62]. In addition, we also demonstrated that CotE acts as a E-cadherin spore ligand [22], and it is plausible that CotE might also contribute to complement binding to *C. difficile* spores. Given that CdeM is unique to *C. difficile* and not present in other members of the Clostridia group [47], the binding mechanism is likely to be unique to *C. difficile*.

Internalization of *C. difficile* spores into intestinal epithelial cells has been shown to contribute to the reservoir of persistent spores that cause disease recurrence after vancomycin treatment [19]. Our results advance this concept to the complement system by demonstrating that, at least *in vitro*, presence of intact complement system contributes to internalization of *C. difficile* spores into intestinal epithelial cells. Previously we showed that fibronectin and vitronectin, both act as molecular bridges to facilitate spore entry in a BclA3-Fn/Vn-integrin dependent manner [19]. Work in *B. anthracis* demonstrated that spore entry is mediated by BclA and C1q interaction via the integrin receptor α2β1 in epithelial lung cells [37]. By contrast, C3a-opsonized bacteria become internalized into epithelial cells through CD46 [64]. These observations suggest that the reduced *C. difficile* spore entry into intestinal epithelial cells observed upon depletion of C1q and C3 could be through α2β1 integrin-and CD46-pathway mediated pathway. While our data suggests that BclA proteins had no role in C1q and C3 binding to *C. difficile* spores, it is tempting to speculate that CdeM may act as a spore ligand in the for C1q-and C3-mediated spore entry into intestinal epithelial cells. Another noteworthy observation is that absence of C1q or C3, internalization was almost completely abolished suggests that C1q and C3 have a higher role than serum Fn and Vn in spore entry into intestinal epithelial cells. Prior work in *B. anthracis* demonstrated that spore entry is mediated by BclA and C1q interaction [37]. These observations contrast with our results where individual removal of C1q and C3 affected spore entry, suggesting that active complement contributes to spore-entry. An intriguing question is whether the intestinal environment becomes enriched in C1q and C3, increasing levels of internalization of *C. difficile* spores into the intestinal mucosa, that lead to persistence and disease recurrence. Indeed, several reports support that during *C. difficile* infection, the intestinal barrier becomes permissible allowing influx of systemic complement [65–67]. Moreover, *C. difficile* toxins enhance intestinal expression of C3 up to 6-fold [33], and our results confirm that an increase of accessible C1q and C3 levels in murine intoxicated intestinal mucosa, suggesting that complement system may have a major role in spore internalization, and likely recurrence of CDI.

Our results also support the notion that CdeM may act as a novel spore-adherence and internalization determinant as observed on the ability of inclusion bodies of CdeM to attach to epithelial cells and internalize in the absence of serum components. The fact that the majority of inclusion bodies of CdeM attached to the adhered junctions, support our prior observation of CdeM as putative a E-cadherin spore-ligand [22], and that CdeM is a putative adhesion and internalization spore-ligand. Despite these insights, the impact of complement and CdeM in adhesion and internalization *in vivo* and its relevance to persistence and pathogenesis is unclear.

## Materials and Methods

### Bacterial strains and growth conditions

Wild-type *C. difficile* R20291, 630Δ*ermB* strain, and the interrupted mutants of *bclA1* (630Δ*erm* CT-*bclA1*), *bclA2* (630Δ*erm* CT-*bclA2*), and *bclA3* (630Δ*erm* CT-*bclA3*) in the NTD domain using CLOSTRON system (kindly donated by Dr. Simon Cutting of the University of London UK [87]) or in R20291, the wild-type (Δ*pyrE*/*pyrE^+^*), Δ*bclA3* deletion mutant using allelic exchange and the complemented Δ*bclA3/bclA3^+^* [44] (Table S1), were grown under anaerobic conditions at 37 °C in a Bactron III-2 chamber (Shellab, OR, USA) in BHIS medium; 3.7% brain-heart infusion broth (BD, USA) supplemented with 0.5% of yeast extract (BD, USA), and 0.1% L-cysteine (Merk, USA) liquid or on agar plated with 1.5% agar (BD, USA). Wild-type *B. subtilis* PY79 and *E. coli* Bl21 were cultured in aerobic conditions at 37°C in Luria-Bertani (LB) medium; 1% trypticase soy agar (BD, USA), 0.5% yeast extract (BD, USA), 1% NaCl (Merck, USA) liquid or on agar plates with 1.5% agar (BD, USA).

### Spore purification

For *C. difficile,* a 1:500 dilution of overnight culture in BHIS was spread into TY plates: 3% trypticase soy agar (BD, USA), 0.5% yeast extract (BD, USA) with 1.5% agar (BD, USA). Plates were incubated for 7 days at 37°C under anaerobic conditions. Then under aerobic conditions, the colonies were harvested with sterile ice-cold Milli-Q water, washed 5 times with water and centrifugation at 18,400×*g* for 5 min. Spores were purified with 50% Nicodenz (Axell, USA) and centrifuged at 18,400×*g* for 45 min previously described [40]. For *B. subtilis* 100µL of an overnight liquid culture in LB was sown into 2×SG agar plates; 0.6% brain heart infusion broth (BD, USA), 1% Bacto peptone (BD, USA), 0.05% MgSO_4_ × 7H_2_O (Merck, USA), pH was adjusted a 7.0 and 1.5% agar (BD, USA) was added and the medium was autoclaved. The following sterile components were added to a final concentration of 1mM Ca(NO_3_)_2_ (Merck, USA), 0.1mM MnCl_2_ × 4H_2_O (Merck, USA), 1 µM FeSO_4_ (Merck, USA), 1% glucose (Sigma−Aldrich, USA). Plates were incubated for 3 days at 37°C in aerobic conditions. The colonies were then harvested with sterile ice-cold Milli-Q water, and the bacterial culture was washed 5 times with water and centrifugation at 18,400×*g* for 5 min. The final layer of the pellet had mainly spore that was separated from the others by washing as was previously described [45, 46].

The spores were purified to obtain suspensions >99% free of vegetative cells, sporulation cells, and cell debris as analyzed by phase contrast microscopy. The number of spores per mL was quantified in Neubauer’s chamber, adjusted at 5 × 10^9^ spores/mL, and stored at −80°C until use.

### Mice used

6-8 weeks C57BL/6 (male or female) were obtained from the breeding colony at the Departamento de Ciencias Biológicas of the Universidad Andrés Bello derived from Jackson Laboratories. Mice were housed with *ad libitum* access to food and water. Bedding and cages were autoclaved, and mice had a 12-h cycle of light and darkness. Mice were housed at 20–24 °C with 40–60% of humidity. All procedures complied with all relevant ethical regulations for animal testing and research. All animal procedures were conducted in accordance with institutional and national guidelines for animal care and use and were approved by the Institutional Animal Care and Use Committee (IACUC) of Texas A&M University.

### Ligated ileal loop assay

C57BL/6 mice were anesthetized in an isoflurane chamber (RWD USA) with 4% isoflurane (Baxter, USA) and were maintained with 2% during the surgery administrated by air. The intestinal loop model was performed as previously described [44, 68]. Briefly, a midline laparotomy was performed, making 1–cm incision in the abdomen, 1.5 cm ileal, and proximal colon (at 1.0 – 1.5 cm from the cecum as a reference) were ligated with silk surgical suture. To evaluate the association of accessible C1q and C3 with *C. difficile* spores in ileum mucosa, the ileum of C57BL/6 were ligated and injected with 5 × 10^8^ *C. difficile* R20291 spores (*n* = 5). The intestine was returned to the abdomen, and the incision was closed. The animals were allowed to regain consciousness. Mice were kept for 5h and were euthanized. The ligated loops were removed and washed gently in PBS before immunostaining, as described below.

### Immunostaining of ileal mucosa

First, extracted ileal ligated loops were longitudinally cut, then washed 3 times by immersion in PBS at RT. For better visualization of the tissues, they were fixed flat at RT. To perform this, tissues were fixed over a filter paper imbibed with 30% sucrose (Winkler, Chile) in PBS–4% paraformaldehyde (Merck, USA) for at least 15 min. Tissues were transferred to a microcentrifuge tube with the same fixing solution and were incubated at 4 °C overnight. Since mucus fixation with cross-linking agents, such as paraformaldehyde, causes the colon’s mucus layer to collapse and shrink to a very tiny lining, the epithelia [69], we did not observe mucus layer in ileal loops. Before immunostaining, the ileum tissues were cut into ∼5 × 5 mm fragments.

On the other way, to visualize the association of *C. difficile* spore with C3 and C1q in the spore infected ileum mucosa, ileum fragments were incubated with primary antibody 1:150 rat monoclonal anti-C3 (#ab11862, Abcam, USA) or rat monoclonal anti-C1q (#ab11861, Abcam, USA) in PBS-3% BSA overnight at 4 °C and subsequently washed and incubated with 1:400 goat anti-rat IgG Alexa 568 (#A-11077, Thermo Fischer, USA) for 3h at RT. To stain *C. difficile* spores, tissues were made permeable by incubation with PBS-0.2% Triton X-100 for 2 h at RT and blocked with BSA-3% BSA for 3 h at RT, and were incubated with a primary polyclonal 1:1,000 anti-*C. difficile* spore IgY batch 7246 antibodies (Aveslab, USA) and to stain actin cytoskeleton with 1:200 phalloidin Alexa Fluor 647 (#A-22287, Thermo Fischer, USA) in PBS-3% BSA overnight at 4°C, then were washed 3 times with PBS and samples were incubated with goat 1:400 anti-chicken IgY secondary antibody Alexa-Fluor 488 (#ab 150173 Abcam USA) and 1:1,000 of Hoechst (ThermoFisher, USA) in PBS-3% BSA for 3h at RT.

To perform double immunostaining of accessible and total C1q and C3 in healthy ileum mucosa, the ileum of 2 independent, healthy C57BL/6 mice of 8 weeks old were removed, and mice were sacrificed. Next, tissues were washed by immersion 3 times in PBS at RT, and they were fixed flat with 30% sucrose in PBS–4% paraformaldehyde, as was described above. Subsequently, tissued cut into ∼5 × 5 mm fragments, then were blocked with PBS-3% BSA (Sigma–Aldrich, USA) for 3 h at RT.

Ileum tissued were incubated with primary antibody 1:150 rat anti-C3 (#ab11862, Abcam, USA) or rat anti-C1q (#ab11861, Abcam, USA) in PBS-3% BSA overnight at 4 °C and subsequently washed and incubated with chicken anti-rat IgG-Alexa Fluor 647 (#A-21472 ThermoFischer, USA) for 3h at RT. To stain the total protein, tissues were made permeable by incubation with PBS-0.2% Triton X-100 for 2 h at RT and blocked with BSA-3% BSA for 3 h at RT. Then tissues were incubated overnight at 4 °C with the same primary antibody used before (1:150). F-actin was stained with 1:150 fluorescently labeled phalloidin Alexa Fluor 647 (#A22287, ThermoFisher, USA). The next day, tissues were washed and incubated 1:400 chicken anti-rat IgG-Alexa Fluor 647 (#A-21472, ThermoFisher, USA) and 4.5µg/mL Hoechst 33342 for 3h at RT.

The aforementioned immune-stained tissues were subsequently mounted with the luminal side-up as previously was described [19, 44]. To do this, the intestinal villi were identified under light microscopy with 10× or 40× magnification and were oriented side-up towards the coverslip. The tissue segment was placed over 5µL of fluorescent mounting medium (Dako, Denmark) applied onto a glass slide. The tissue was covered with 15 µL fluorescent mounting medium and closed with a coverslip. Coverslips were affirmed to the glass slide with vinyl tape to hold the tissue sections in place and were allowed to cure for at least 24h before imaging.

### Confocal and Epifluorescence analysis of immunostained tissues

For confocal imaging, Leica SP8 was used with HPL APO CS2 40× oil, 1.30 numerical aperture. Signals, 3 PMT spectral detector PMT1 (410-483) DAPI PMT2 (505-550) Alexa-Fluor 488 PMT3 (587-726) Alexa-Fluor 555. Emitted fluorescence was split with dichroic mirrors DD488/552. To quantify the ileum mucosa cells with immunodetected accessible C1q and C3, confocal images (1,024×1,024 pixels) with a 2µm *z*-step size were filtered with Gaussian Blur 3D (sigma *x*: 0.6; *y*: 0.6; *z*:0.6) and quantified with cell counting plug-in of ImageJ.

To quantify the spore number associated with accessible C1q or C3, confocal images (1,024×1,024 pixels) with a 0.7 µm *z*-step were quantified with cell counting plug-in of ImageJ. Fl. Int. profiles were performed with the Plot Profile plug-in of ImageJ. Three-dimensional reconstructions of ileum mucosa were performed with a 3D projection plug-in of ImageJ software (NIH, U.S.A). Villi were visualized by Hoechst and phalloidin signals.

### CDI mouse model and fesses collection

To induce *C. difficile* susceptibility, an antibiotic cocktail containing 40 mg/kg kanamycin (Sigma–Aldrich, USA), 3.5 mg/kg gentamicin (Sigma–Aldrich, USA), 4.2 mg/kg colistin (Sigma–Aldrich, USA), 21.5 mg/kg metronidazole (Sigma–Aldrich, USA) and 4.5 mg/kg vancomycin (VAN) (Sigma–Aldrich, USA) was administered via gavage for 3 days (days -6 to -4 before the infection) as was previously described [70]. One day before the infection (day -1), 10 mg/kg clindamycin (Sigma–Aldrich, USA) was administered intraperitoneally to 3 independent C57BL/6 mice. On the next day (day 0) three fresh fesses were collected for each mouse and then were orally infected with 100µL containing 5 × 10^7^ *C. difficile* spores R20291. And 2 days post-infection other 3 independent fresh fesses were collected for each mouse and were stored at -80°C. to be tested for C3 and C1q using dot blot as is described below.

### Induction of CdeM inclusion bodies

The overexpression of CdeM-6×His and CdeC-6×His was performed as previously described [48]. Briefly, transformed strains containing the plasmid pARR21 with *cdeM* gene of *C. difficile* R20291 fused to 6×His (CdeM-6×His tag) was cultured overnight in LB-0.5% glucose (LBG) at 37°C with shaking at 1×*g*. 600mL of fresh LBG were inoculated with 5mL of the overnight culture and were evaluated frequently to an optical density 0.7 to 0.9 at 600nm. Then the cultures were induced with 0.5mM if isopropyl-β-D-thiogalactoside (IPTG) for 16h at 37°C with shaking at 1×*g*. The cells were precipitated by centrifugation at 5,853×*g* and the pellets were stored at -80°C.

Next, pellets were thawed and resuspended in 30mL of 10 mM EDTA, 0.1% Tween-20, 10 mg/mL of lysozyme in PBS for 1 h at 37°C. Then were sonicated 6 times at 12W for 15s with 3 min incubation on ice each time and centrifuged at 5,853×*g* for 45 min at 4°C. The supernatant was discarded, and the pellet was washed with 2% Triton X-100, 1mM phenylmethylsulfonyl fluoride (PMSF) 3 times in PBS by centrifugation at 5,853 ×*g* at 4°C. Then pellets were sonicated at 12W for 15s on ice and to separate the cells from the IBs, were purified through density gradient with 45% Nycodenz (Accurate Chemicals, U.S.A) by centrifugation at 5,853×*g* for 50 min×*g* at 4°C. Finally, the IBs were washed twice with PBS-1mM PMSF by centrifugation at 5,853×*g* for 20 min and were counted by Neubauer chamber and adjusted to 5×10^9^/mL and stored at −80 °C until use.

### Cell culture

Caco-2 cells were obtained from the ATCC (USA) and were grown at 37°C with 5% CO_2_ with DMEM High Glucose (Dubecco’s modified Eagle’s minimal essential medium) (Hyclone, USA) supplemented with 10% fetal bovine serum (HyClone, USA) and 100U/mL penicillin and 100µg/mL streptomycin (Hyclone, USA). For infection experiments, cells were plated over a glass coverslip in a 24-wells plate and cultured for 2-days post-confluence.

### Solid-phase binding assay

1.6 × 10^7^ *C. difficile* spores R20291, 630Δ*ermB*, 630Δ*erm* CT-*bclA1*, 630Δ*erm* CT-*bclA2*, 630Δ*erm* CT-*bclA3* or R20291 Δ*pyrE*/*pyrE*^+^, Δ*bclA3* or Δ*bclA*/*bclA3*^+^ or CdeM IBs were incubated in wells of a 96-well plate, or 1% BSA as control. Incubation was performed in PBS at 4°C overnight. The wells were washed 5 times with PBS to remove unbound spores and then incubated with TTBS-1% fish gelatin for 1h at RT. Then wells were incubated with increasing concentrations of 0.001, 0.01, 0.1, 1, 10% NHS, C1q-Dpl, C4-Dpl, FB-Dpl, or 0.005, 0.05, 0.5, 5, or 50 µg/mL of human purified C1q or C3 (Complement Technology, USA) or BSA (Sigma-Aldrich, USA), in GTTBS^++^ buffer for 45 min at 37°C and then were then washed 3 times with TTBS. Wells were incubated with 1:10,000 goat polyclonal antibodies anti-Human C1q, anti-Human C3, anti-human C4 (#A205), anti-human Factor B (#A235), (Complement Technology, USA), or with 1:2,500 mouse monoclonal antibody against BSA as indicated (#ab3781 Abcam, USA) in GTTBS^++^. Then were washed with TTBS three times and were incubated with 1:30000 of secondary antibody conjugated to HRP in GTTBS^++^ for 1h at RT (rabbit anti-goat IgG-HRP #605-4302 or goat anti-mouse IgG-HRP #610-1302, Rockland, USA). Wells were washed 5 times with TTBS. Finally, the plates were incubated with 50 μL 20 mg OPD substrate in 10mL H_2_O_2_ 5uL (Sigma, USA), for 10 min at RT, the reaction was stopped with 4N of H_2_SO_4_ and read in plate spectrophotometer Tecan Infinite^®^ F50 at 492nm.

### Evaluation of the binding of C3 and C1q to *C. difficile* spore and IBs by immunofluorescence

1 × 10^7^ *C. difficile* spores were incubated with 10 µg/mL of human purified C3 or C1q diluted in buffer GTTBS^++^ for 45 min at 37 °C. The spores were washed three times with TTBS and were added to glass covers previously poly-lysine-treated for 10 min (Sigma-Aldrich, USA), until dry at RT. Then, samples were fixed with 4% paraformaldehyde for 10 min and washed 3 times with TTBS and then blocked with TTBS 0.1% fish gelatin (GTTBS), for 1 h at RT. The covers were then incubated for 1 h at RT with polyclonal antibodies against anti-C1q or anti-C3, respectively (Complement Tech, USA). Subsequently, the covers were washed 3 times with TTBS and incubated for 1h at RT with 1:200 secondary antibody donkey anti-goat-CFL 488 (#SC362255 Santa Cruz Biotechnologies, USA) in TTBS. Finally, were washed 3 times with TTBS and one with sterile Milli-Q water and were incubated until dry at RT and then mounted using fluorescent mounting medium (Dako North America, Inc.). Finally, were sealed with clear nail polish.

In the case of inclusion bodies, 2.7 × 10^7^ *C. difficile* spores or CdeM IB were incubated with 10% NHS or 5% BSA for 1 h at 37°C and washed twice with PBS by centrifugation at 18,659×*g* for 5 min and resuspended with sterile PBS. IB and spores were fixed to a poly-L-lysine pre-treated cover glass for 10 min at RT. Once mounted, samples were dried at 37 °C for 5 min and fixed with PBS-4% PFA for 10 min at RT, washed twice with PBS and blocked with PBS-1% BSA for 1h at RT and incubated with 1:1,000 goat polyclonal antibodies anti-human C1q (#A200), or 1:1,000 anti-human C3 (#A213) and then washed 3 times with PBS and incubated with 1:500 chicken anti-mouse IgG-Alexa Fluor 488 (#A-21200, Invitrogen, USA), or Donkey anti-goat IgG-CFL 488 (#SC362255 Santa Cruz Biotechnologies, USA). Finally, they were mounted on with fluorescent mounting medium (Dako, USA) and sealed with transparent nail polish.

### Evaluation of CdeM IBs internalization by immunofluorescence

Caco 2 cells were grown on coverslips in 24-wells tissue culture plate until they reached a 2 days post confluence monolayer. The cells were incubated with DMEM without FBS 1h at 37°C, simultaneously *C. difficile* spores and CdeM inclusion bodies (IB) at MOI 10 were incubated with 20 uL of NHS 1h at 37°C. Then the spores and CdeM IB were suspended in 180 uL of DMEM that was added to each well and were incubated for for 5 h at 37°C. Infected cells were washed three times with PBS to remove the spores and CdeM IB that were not bond to the cells and were fixed with 4% PFA for 10 min at room temperature. The fixed cells were washed 2 times with PBS, blocked with 1% BSA (Sigma) overnight at 4°C mouse anti-6xHis (Rockland 800-6567625) for CdeM IB detection or 1:50 anti *C. difficile* spore goat serum for 1 h at room temperature. The coverslips were washed twice with PBS and incubated with 1:500 Donkey anti-mouse Alexa Fluor 488 (Abcam ab15109) or 1:400 Donkey anti-goat IgG CF 568 (Sigma SAB4600074) for 1h at room temperature. The cells were permeabilized with Triton 0.2% 10 min, blocked with 1% BSA 1h, incubated with 1:400 mouse anti-6xHis (Rockland 800-6567625) for CdeM IB detection or 1:50 anti *C. difficile* spore goat serum for 1 h at room temperature and incubated with 1:500 Donkey anti-mouse Alexa Fluor 568 (Abcam ab175700) or 1:400 Donkey anti-goat IgG Texas Red (Santa Cruz SC3856) for 1h at room temperature. The coverslips were washed twice with PBS, once with MiliQ water, dried and mounted using Dako Fluorescence Mounting medium and sealed with nail polish. Samples were analyzed in Olympus fluorescence microscope.

### Epifluorescent imaging acquisition and analysis

Epifluorescence images were acquired using an Olympus BX53 fluorescence microscope with UPLFN 100 × oil objective (1.30 numerical aperture). Images were captured using the Qimaging R6 Retiga camera. To quantify the fluorescence intensity of C1q and C3 for *C. difficile* spores and CdeC, CdeM IBs, the contour of the spores was draw in the phase contrast image using the tool "freehand selection" of ImageJ (NIH, USA) and then the fluorescence intensity (Fl. Int.) and the area were measured for 150 particles. Also, the Fl. Int.of the background was measured. The [(Fl. Int._particle_/area_particle_) – (Fl. Int._Background_/area _Background_)] is shown.

### Spore preparation for transmission electron microscopy and immunogold electron microscopy

The binding of the C1q and C3 to *C. difficile* spore was realized as was described before with modifications [44]. Briefly, 4 × 10^7^ *C. difficile* spores were incubated with 10 µg/mL of human purified C3 or C1q for 45 min at 37 °C in PBS-0.2% BSA. The spores were then washed with PBS and centrifugation at 18,400×*g* for 5 min at RT three times. Then the spores were pelleted by one cycle of centrifugation at 18,400 × *g* for 10 min. Pellets were then blocked in PBS-1% BSA, for 30 min at RT, and sedimented at 18,400 × *g* for 10 min at RT. Then spores were incubated with primary antibody goat anti-human C3 or anti-human C1q (Complement tech, USA), in PBS-1% BSA or 1% BSA as control for 1h at RT. The excess of antibody was eliminated by three cycles of centrifugation at 18,400×*g* for 5 min at RT and resuspension in PBS-0.1% BSA. Spore suspensions were then incubated for 1 h with 1:20 donkey anti-goat IgG antibody coupled to 12 nm gold particles (#ab105269, Abcam, USA), in 1% BSA-PBS for 1h at RT. And were washed with PBS at 18,400×*g* for 5 min three times. Subsequently, samples were fixed with freshly prepared with 2.5% glutaraldehyde 1% paraformaldehyde in 0.1 M cacodylate buffer (pH 7.2) at 4 °C overnight, rinsed in cacodylate buffer, and stained with 1% tannic acid for 30 min. Then samples were dehydrated with 30% acetone (with or without 2% uranyl acetate) for 20 min, 50% for 20 min, 75% for 20 min, 90% for 20 min, and twice with 100% for 20 min, embedded in spurs resin at ratio acetone: spurs of 3:1, 1:1, and 1:3 for 40 min each and then resuspended in spurs for 4h and baked overnight at 65 °C, and prepared for transmission electron microscopy. 90 nm sections were obtained with a microtome and placed on glow discharge carbon-coated grids for negative staining and double lead stained with 2% uranyl acetate and lead citrate. Grids were visualized with a Philips Tecnai 12 Biotwin electron microscope of the Universidad Católica de Chile. Positively associated spores were quantified and aggrouped by the morphotype of thin or thick exosporium.

### Sample preparing for immunoblotting

To evaluate the co-precipitation of C3 protein, 2.5 × 10^7^ spores or vegetative cells of *C. difficile* R20291, 630Δ*ermB* or *B. subtilis*. were incubated with 10% NHS (#NHS, Complement Technology, Inc, USA), 10% C3-Dpl, (#A314, Complement Technology, Inc, USA). Meanwhile to evaluate the co-precipitation of C1q spores or vegetative cells were incubated with 10% NHS, 10% C1q-Dpl (#A300, Complement Technology, Inc, USA). To evaluate the co-precipitation of C3 and C1q to CdeM IB, 5 × 10^7^ CdeM inclusion bodies were incubated with 10% NHS, 5% BSA or DMEM for 1 h at 37°C and washed twice with PBS by centrifugation at 18,659×*g* for 5 min and resuspended with PBS. As a negative control, samples were incubated with serum-free medium (SFM), in GTTBS^++^ buffer [15 mM Tris (OmniPur®, USA), 150 mM NaCl (Merck, USA), 0.5 mM MgCl_2_ (Winkler, Chile), 0.15 mM CaCl_2_ (Winkler, Chile), 0.1% Tween-20 (Merck,USA), and 1% fish gelatin (Sigma-Aldrich, USA), pH 7.3] for 45 min at 37 °C. This buffer is an adaptation of GVS^++^ buffer used in complement activation assays that contains Mg^++^ that is a cofactor for the C3 and C5 convertases and Ca^++^ for C1 complex activation in the classic via of the complement. The veronal buffer was changed for TBS because is a less controlled substance and have a low chelating activity for the aforementioned cofactors also the saline concentration was adjusted to physiological ion strength using 150 mM of NaCl. Fish gelatin was used as a blocking solution and to prevent the absorption of protein to plastic materials. The spores were then washed 3 times in 0.1% Tween-20 in Tris-Buffered saline (20 mM Tris, 500 mM NaCl, pH 7.3; TTBS), before anti-C3 or C1q immunoblotting as described below. In the case of *C. difficile* spores R20291 and 630Δ*ermB* incubated with C3, after incubation, samples were centrifugated at 18,400×*g* and the supernatant was collected and, pellets were washed and were tested for anti C3 by immunoblotting as described below. As positive control 0.2 µL of NHS or 1μg of purified C3 or C1q as appropriate were loaded in the gel. To evaluate the presence of C3a in the serum stocks, 10µL of NHS, C3-Dpl, C1q-Dpl, C4-Dpl, FB-Dpl, were processed for immunoblotting anti-C3a as is described below.

### Immunoblotting

Once the samples were prepared as was described above, were suspended in 2× SDS-page sample loading buffer (Bio-rad, USA), boiled, and electrophoresed on 4% and 12% acrylamide SDS-PAGE gels. For CdeM IBs we used a 15% acrylamide SDS-PAGE. Then proteins were transferred to a nitrocellulose membrane (Bio-rad, USA). Membranes were blocked with TTBS-3% BSA (Winkler, Chile), for 1h at room temperature (RT). Then were incubated with 1:5,000 goat polyclonal antibodies anti-human C1q (#A200), 1:5,000 anti-human C3 (#A213), or 1:5000 rabbit anti-human C3a (#A218), prepared in TTBS with 3% BSA for 1 h at RT (all antibodies were purchased from Complement Technology, USA). Then membranes were washed three times for 5 min with TTBS and incubated with 1:10,000 of secondary antibody rabbit anti-goat IgG conjugated to horseradish peroxidase (HRP; Rockland, USA) or goat anti-rabbit IgG-HRP (Abcam, USA) diluted in TTBS-3% BSA. The membranes were washed three times with TTBS and once with TBS for 5 min each for posterior incubation with chemiluminescent substrate (Clarity Western ECL, Biorad, USA) Images were acquired using FOTO/Analyst® FX (FOTODYNE Inc., USA). Immunoblot band intensities were quantified by densitometry using ImageJ software (NIH, USA). For each membrane, regions of interest of identical dimensions were drawn around the bands of interest, and the integrated density was measured. Background signal was determined from an adjacent region of the membrane lacking detectable bands and subtracted from the corresponding band intensity. Background-corrected densitometric values were used for comparisons among experimental conditions and, when indicated, were normalized to the corresponding control condition.

### Two-dimensional gels and far-western blot

1 × 10^9^ spores of *C. difficile* R20291 were treated with a solution of 8M Urea, 50 mM DTT, 1% SDS in 50 mM Tris Buffer pH 8.0 for 2 h at 37 °C. Spores were sonicated at 35% of frequency 5 times. The spores were then centrifuged for 7min at 18,400×*g* and the supernatant was dialyzed in water to promote protein precipitation. The samples were then rehydrated with 8 M urea, 2 M thiourea, 65 mM DTT, 0.2% ampholytes, bromophenol blue (traces), in 4% CHAPS buffer for 1 h at 37°C. The samples were centrifuged at 14.000×*g* for 10 min. The supernatant was loaded to isoelectric focusing in a 7cm strip (ReadyStrip IPG, Bio-Rad, USA) that were placed on the sample and were incubated at RT for 1h. The rails were covered with mineral oil and incubated for overnight at RT. The isoelectric focusing was performed in Hoefer IEF100 (Hoefer, Inc Germany), at 0.1 W constant per 1h, 0.5 W constant per 3 h, 1,000 V for 1h in 1.5 M Tris equilibrium buffer pH 8.8, 29.3% glycerol, 2% SDS and bromophenol blue (traces). Proteins were separate by size in a 12% SDS-PAGE and ran at 20 mA/gel (fixed) for 3 h. For ligand identification by far-western blot, two gels were treated in pairs. One gel was stained with Coomassie Blue G-250 (Bio-Rad). The proteins of the other were transferred to nitrocellulose membrane (Bio-Rad, USA). The membrane was blocked with TTBS-3% BSA in and then incubated with 5 µg/mL of human purified C1q or C3 (Complement-tech, USA), in 1% BSA and 0.1% Tween-20 in 50 mM TBS, for 16h at 4 °C. Subsequently, the membranes were washed 3 times for 10 min with TTBS. The membranes were incubated with 1:10,000 goat polyclonal antibodies anti-human C1q or anti-human C3 (Complement Tech, USA) in TTBS-3% BSA for 1 h at RT. Then were washed 3 times for 5 min with TTBS and then were incubated with 1:10,000 polyclonal rabbit anti-goat HRP for 1h at RT (#ab6741 Rockland, USA). After incubation, the membranes were washed three times with TTBS and once with TBS for 5 min each for posterior incubation with Chemiluminescent substrate (Clarity Western ECL, Biorad, USA)-Images were acquired using FOTO/Analyst® FX (FOTODYNE Inc., USA). mmunoblot band intensities were quantified by densitometry using ImageJ software (NIH, USA). For each membrane, regions of interest of identical dimensions were drawn around the bands of interest, and the integrated density was measured. Background signal was determined from an adjacent region of the membrane lacking detectable bands and subtracted from the corresponding band intensity. Background-corrected densitometric values were used for comparisons among experimental conditions and, when indicated, were normalized to the corresponding control condition.

### Protein identification by mass spectrometry

Immunoreactive proteins were identified using duplicate 2-DE gels. Proteins from one gel were transferred to a nitrocellulose membrane at 220 mA for 90 min. The membrane was blocked overnight in 3% BSA (Sigma) and subsequently incubated for 1 h at room temperature with goat polyclonal antiserum raised against purified *C. difficile* spores. After washing, the membrane was incubated for 1 h at room temperature with an HRP-conjugated anti-goat secondary antibody, and immunoreactive spots were visualized using standard chemiluminescent procedures. Corresponding protein spots were excised from the duplicate gel for identification by mass spectrometry. Excised gel spots were subjected to in-gel digestion with sequencing-grade chymotrypsin (Promega, Madison, WI, USA), followed by nanoRPLC in-line desalting as previously described [71, 72]. Peptides were analyzed by nanoUPLC-ESI-MS/MS. Samples were desalted and concentrated using a Symmetry C18 RP-Trap column (5 µm, 180 µm i.d. × 20 mm; Waters Corp., Milford, MA, USA) and separated on a BEH130 C18 reversed-phase nanoAcquity UPLC column (1.7 µm, 100 µm i.d. × 100 mm; Waters Corp.). The chromatographic gradient consisted of 3% mobile phase B from 0 to 2 min, followed by a linear increase from 3% to 80% B between 2 and 40 min. Mobile phase A consisted of water/formic acid (99.9:0.1, v/v), whereas mobile phase B consisted of acetonitrile/formic acid (99.9:0.1, v/v). The flow rate was maintained at 400 nL/min using a nanoAcquity UPLC system (Waters Corp.). Eluting peptides were analyzed by nano-ESI using a Q-Tof Synapt G1 HDMS mass spectrometer (Waters, Milford, MA, USA) operated in positive-ion mode. Data acquisition consisted of one full precursor MS scan over an m/z range of 400–1500, followed by four MS/MS scans of the most abundant precursor ions under dynamic-exclusion conditions. Spectra were deconvoluted and processed using MassLynx v4.1 software (Micromass, UK), and peak lists were generated in PKL format to identify singly and multiply charged precursor ions from the raw mass-spectrometry files. Instrument calibration in MS/MS mode was performed using 100 fmol of human [Glu1]-fibrinopeptide B, yielding a root mean square residual of 7.484 × 10^−4 amu, corresponding to 7.76 × 10^−1 ppm. The parent-ion and fragment-ion mass ranges were 400–1500 Da and 65–1500 Da, respectively.

Protein identification was performed using Mascot Server v2.5.1 (Matrix Science, UK) in MS/MS ion-search mode against the *Peptoclostridium difficile* strain R20291 database obtained from UniProt (version 20171130; 3,507 sequences; 1,134,223 residues). Carbamidomethylation of cysteine was specified as a fixed modification, whereas deamidation of asparagine and glutamine and oxidation of methionine were included as variable modifications. Searches allowed one missed cleavage, used monoisotopic masses, and applied precursor-and fragment-mass tolerances of 20 ppm and 0.3 Da, respectively. The ion-score or expected-value cutoff was set at 5. MS/MS spectra were searched using a 95% confidence threshold (P < 0.05), corresponding to a minimum Mascot score of 17 for peptide identification. Mascot error-tolerant searches were additionally performed to identify potential peptides not assigned during the initial search. When identified peptides matched multiple protein entries equally well, only proteins detected in at least two independent replicates were retained in the final identification list. Additional analyses were performed using an Ultimate 3000RS UHPLC system (Thermo Scientific, Bremen, Germany) coupled through a nanoelectrospray ion source to a Q Exactive Plus quadrupole-Orbitrap mass spectrometer (Thermo Scientific), following previously described procedures [73, 74]. Mascot Distiller v2.6.2.0 (Matrix Science) and Proteome Discoverer v2.1 (Thermo Scientific) were used to generate Mascot Generic Format (MGF) peak lists from the original raw files for downstream database searching.

The spots in the SDS gel were identified using the spots marked in the membranes, and spots in gel were cut, and protein were in-gel digested with chymotrypsin (ThermoFisher, U.S.A) Generated peptides were separated by nanoUPLC followed by MS/MS using a Quad-TOF MS, and corroborated by Ultimate 3000 RS nano UHPLC coupled to Q Exactive Plus Orbitrap MS [73, 75] in the SD-BRIN Proteomics Facility of the University of South Dakota, USA. The peptide sequences obtained by MS/MS were contrasted with *Clostridium difficile* database R20291_refseq_20171130 using Mascot algorithm using MS/MS ion search and list of score tables were analyzed for better coverage of the putative sequences for spore proteins (See Supplementary Information: S1_File_Mascot_MS-MS_Peptide_Reports.pdf and Table S2 MS MS data per spot summary).

### Dot Blots

200mg of mouse fecal samples were resuspended in 750uL of ethanol 50% and the 3µL were spotted in a nitrocellulose membrane and let it dry for 30 min at RT. Meanwhile, to test the presence of C3 or C1q in the used serum, dilution of NHS, C1q-Dpl, C3-Dpl, C4-Dpl, FB-Dpl were made at 2% for C3 detection and 10% for C1q detection. Then 3μL were spotted and allowed to dry in a nitrocellulose membrane as described before. Then the membranes were blocked with TTBS-3% BSA for 1h at RT and incubated with primary antibodies anti-mouse C1q or anti-mouse C3 prepared 1:5,000 with TTBS-3% BSA for 1h at RT. The membranes were washed 3 times for 5 min with TTBS and then incubated with 1:10000 dilution of secondary antibody conjugated with HRP for 1h at RT. The membranes were washed three times with TTBS and once with TBS for 5 min each for posterior incubation with chemiluminescent substrate (Clarity Western ECL, Biorad, U.S.A.), and the capture of the images using the imaging equipment FOTO/Analyst® FX (FOTODYNE Inc., USA). In the other hand, to quantify the optical density of dot blot membranes, a custom circle with the same size it was drawn around each dot in the non-inverted RGB images using the "oval" tool of ImageJ and the Integrated Density was measured. For each circle, the [(Fl. Int_circle_/area_circle_) is shown.

### Infection and immunofluorescence assay

Caco-2 cells cultured over cover glass in a 24-well plate were washed three times with PBS and *C. difficile* spores were incubated separately in normal human serum (NHS), C1q depleted human serum (C1q-Dpl), C1q-Dpl + 70 µg/mL of purified C1q, C3 depleted serum (C3-Dpl), C3-Dpl + 70 µg/mL of purified C3, or serum free medium (SFM), for 45 minutes at 37°C (All sera are purchased from Complementtech, U.S.A). Before infecting the spores, three times. The infection was carried out at an MOI of 10 and was synchronized by centrifugation at 604 g for 5 minutes then incubating at 37°C for 4 hours. After the time has elapsed the wells were washed 3 times to remove the spores that were not they brought the cells together. To continue with the immunofluorescence the cells were fixed at 4% of paraformaldehyde for 15 minutes at RT and washed 3 times with PBS. Then non-specific epitopes were blocked with 1% BSA on PBS Tween-20 0.05% overnight at 4°C and incubated with primary antibody raised in goat against *C. difficile* spores in dilution 1:200 for 1 hour at RT. After washing, it was incubated with secondary antibody conjugated to fluorophore CFL488 at a dilution of 1:500 for 1 hour at RT. The coverslips were washed 3 times and incubated with Hoechst 4 mg/mL dilution 1:1.000 for 10 minutes at RT. The coveralls were washed 3 times with PBS, were dried, assembled and sealed at RT. The cover plates with the samples were analyzed under the OLYMPUS epifluorescence microscope. The adhesion was estimated by counting the number of spores in the cells and estimating the percentage of the condition with 100% of human serum. The internalization was estimated by subtracting the extracellular spores marked in green when total spores were identified in phase contrast and estimated relative to the condition with full human serum (100%).

## Acknowledgments

We acknowledge Scarlet Troncoso-Cotal and Alba Romero for their support in the purification of CdeM inclusion bodies. We also Acknowledge Dr. Jon Skare from Texas A&M University for sharing C1q deficient mice. This work was also supported by grants from Awards from ANID -Millennium Science Initiative Program -NCN17_093, Texas A&M University Start Up funds, and 5R01AI177842 from the National Institute of Allergy and Infectious Diseases, all to D.P-S.

## Authoŕs Contribution

**Conceptual Design** (Formulation of hypothesis, development of study objectives, defining experimental, statistical and analytical procedures:

**MPG, CBS, PCC, DPS,**

**Data acquisition** (labwork, theoretical calculations, literature searches)

**MPG, CBS, PCC, NMB, EC**

**Analysis and interpretation of the data** (making sense of and presenting the results, data analysis (statistical or other)

**MPG, CBS, PCC, NMB, EC, FG, DPS**

**Writing publication** (creating all or substantial part of the scientific and technical product)

**MPG, PCC, FG, DPS**

**Critical Revision of Publication** (reworking all the scientific and technical product for intellectual content before submission (not just spelling and grammar checking)

**MPG, PCC, FG, DPS**

**Approval of final publication** (providing approval of the final product to be published)

**DPS**

**Supervision** (oversight and responsibility for the study, general supervision of the research group)

**DPS**

**Resources** (funding, equipment, facilities, personnel vital to the project)

**DPS**

---

**Figure 12 | C1q deficiency alters the clinical course of *C. difficile* infection in mice.** Wild-type (WT) and C1q-deficient mice were infected with *Clostridioides difficile* strain R20291 and monitored for 13 days. Mock-treated WT and C1q-deficient mice were included as uninfected controls. **A,** Changes in body weight during the experimental period, expressed as percentage of the initial body weight. **B,** Daily weight-loss clinical score for R20291-infected WT mice, R20291-infected C1q-deficient mice, and mock-treated controls. Symbols represent individual animals and bars indicate the mean ± SEM. **C,** Cumulative percentage of R20291-infected WT and C1q-deficient mice that developed diarrhea during the early phase of infection. **D,** Cumulative percentage of mice exhibiting diarrhea during the later phase of the experiment in mock-treated controls and R20291-infected WT and C1q-deficient mice. Data represent **[n = X mice/group]** from **[X independent experiments]**. Error bars in A and B indicate mean ± SEM.

## Supplementary Figures

**Supplementary Fig. 1.**
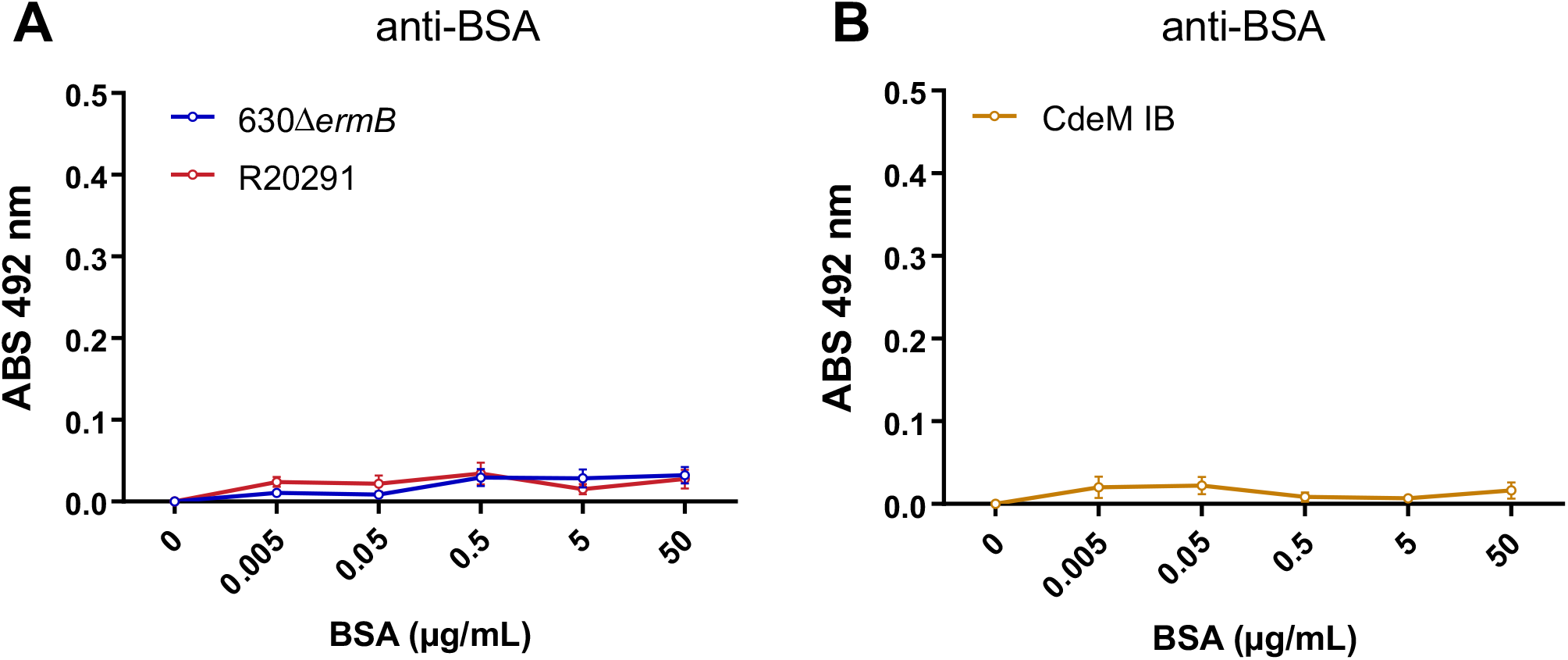
BSA does not associate with *C. difficile* spores or CdeM inclusion bodies. Solid-phase binding assays were performed using BSA as a negative-control protein. Wells coated with **A,** *C. difficile* spores of strains 630Δ*ermB* or R20291, or **B,** CdeM inclusion bodies (IBs), were incubated with increasing concentrations of BSA (0– 50 μg/mL). After washing, associated BSA was detected by immunoassay using anti-BSA antibodies, and absorbance was measured at 492 nm. Minimal signal was detected across all BSA concentrations for either *C. difficile* spores or CdeM IBs, supporting the specificity of the interactions observed with complement proteins. Data represent the mean ± SEM of six wells collected from three independent experiments.

**Supplementary Fig. 2.**
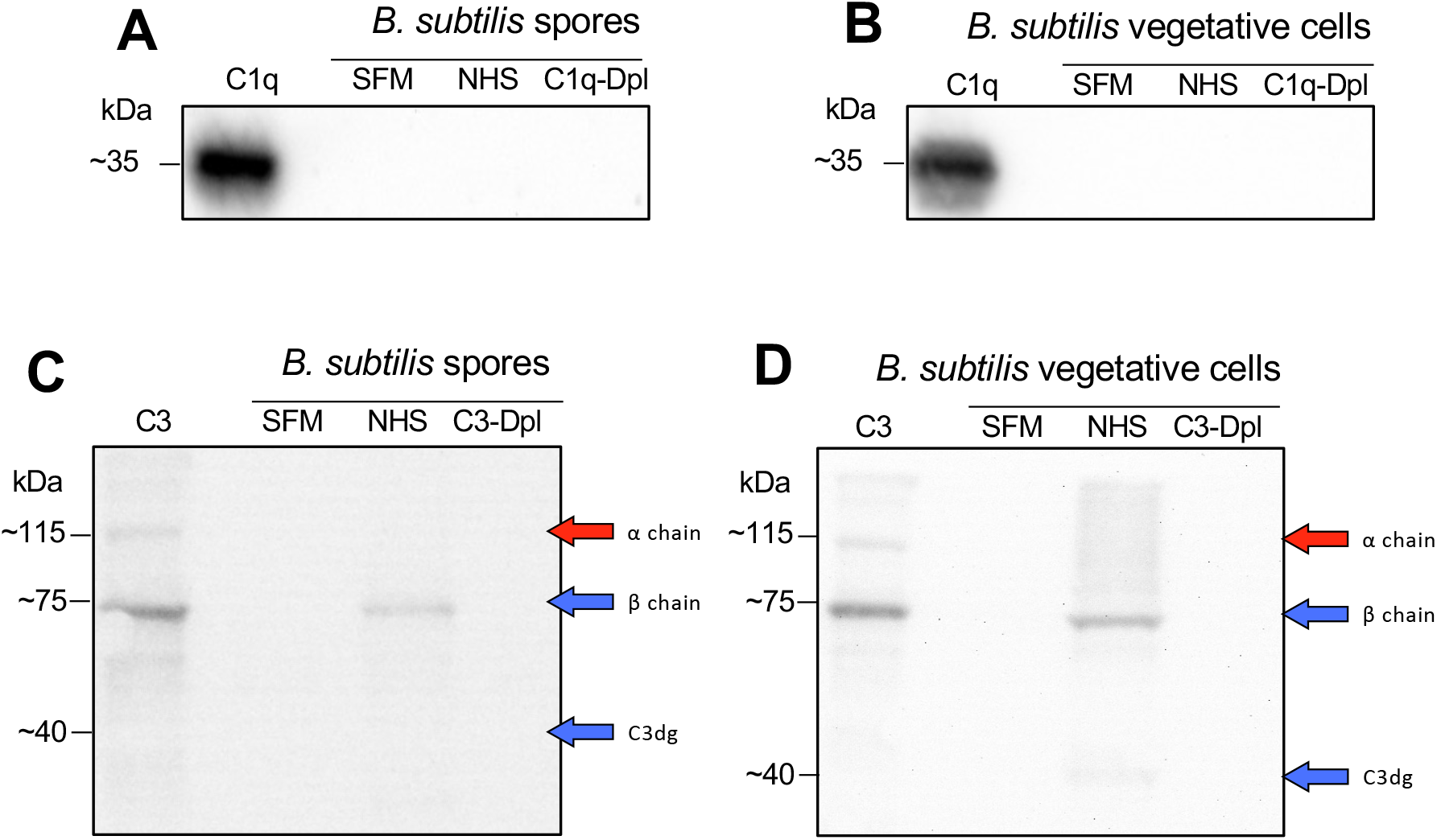
C3, but not C1q, associates with *B. subtilis* spores and vegetative cells. *Bacillus subtilis* PY79 spores or vegetative cells (2.5 × 10^7) were incubated with 10% normal human serum (NHS), the corresponding complement-depleted serum, or serum-free medium (SFM), washed, and analyzed by immunoblotting. **A, B,** Detection of C1q associated with **A,** *B. subtilis* spores or **B,** vegetative cells following incubation with NHS, C1q-depleted serum (C1q-Dpl), or SFM. Purified human C1q was included as a positive control. No detectable C1q-associated signal was observed in either spores or vegetative cells. **C, D,** Detection of C3 associated with **C,** *B. subtilis* spores or **D,** vegetative cells following incubation with NHS, C3-depleted serum (C3-Dpl), or SFM. Purified human C3 was included as a positive control. A C3-immunoreactive band corresponding to the β chain (∼75 kDa) was detected in NHS-incubated spores and vegetative cells, whereas the α chain (∼115 kDa) and C3dg (∼40 kDa) were not detected in the associated fractions. The positions of the C3 α chain, β chain, and C3dg are indicated. Each panel is representative of two independent experiments.

**Supplementary Fig. 3.**
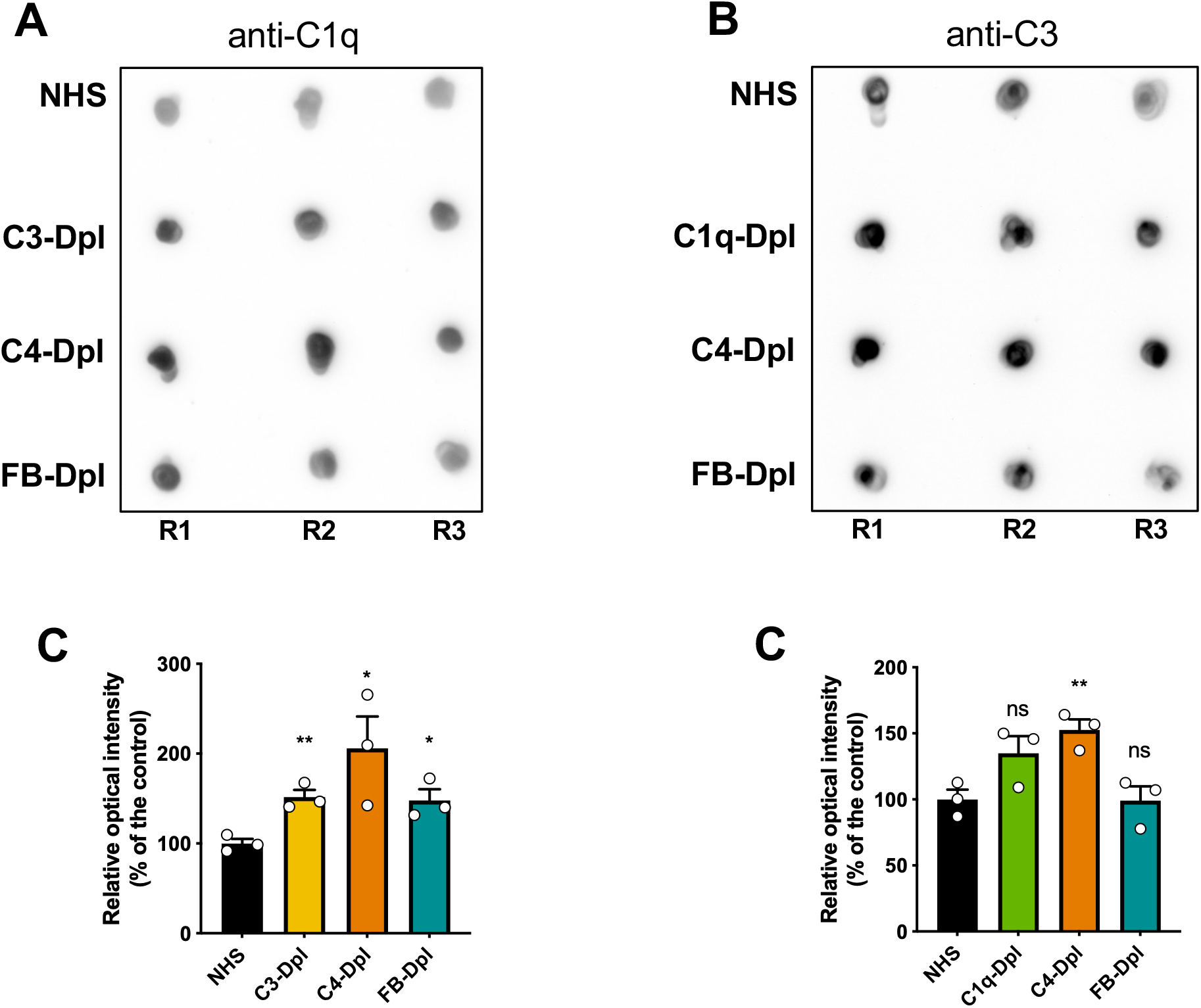
C1q and C3 are present in the complement-depleted sera used in this study. Dot-blot analysis was performed to verify the presence of C1q and C3 in the depleted human sera used throughout the study. **A,** Detection of C1q in normal human serum (NHS), C3-depleted serum (C3-Dpl), C4-depleted serum (C4-Dpl), and factor B-depleted serum (FB-Dpl). Two microliters of 20% serum were spotted onto nitrocellulose membranes and probed with anti-C1q antibodies. **B,** Detection of C3 in NHS, C1q-depleted serum (C1q-Dpl), C4-Dpl, and FB-Dpl. Two microliters of 2% serum were spotted onto nitrocellulose membranes and probed with anti-C3 antibodies. R1, R2, and R3 represent technical replicates. **C,** Quantification of C1q dot-blot signal intensity in the sera shown in **A**, expressed relative to NHS, which was set to 100%. **D,** Quantification of C3 dot-blot signal intensity in the sera shown in **B**, expressed relative to NHS, which was set to 100%. Individual points represent technical replicates, and bars indicate the mean ± SEM. Statistical significance was determined using a two-tailed unpaired Student’s *t*-test relative to NHS. ns, not significant; *, *P* < 0.05; **, *P* < 0.01.

**Supplementary Fig. 4.**
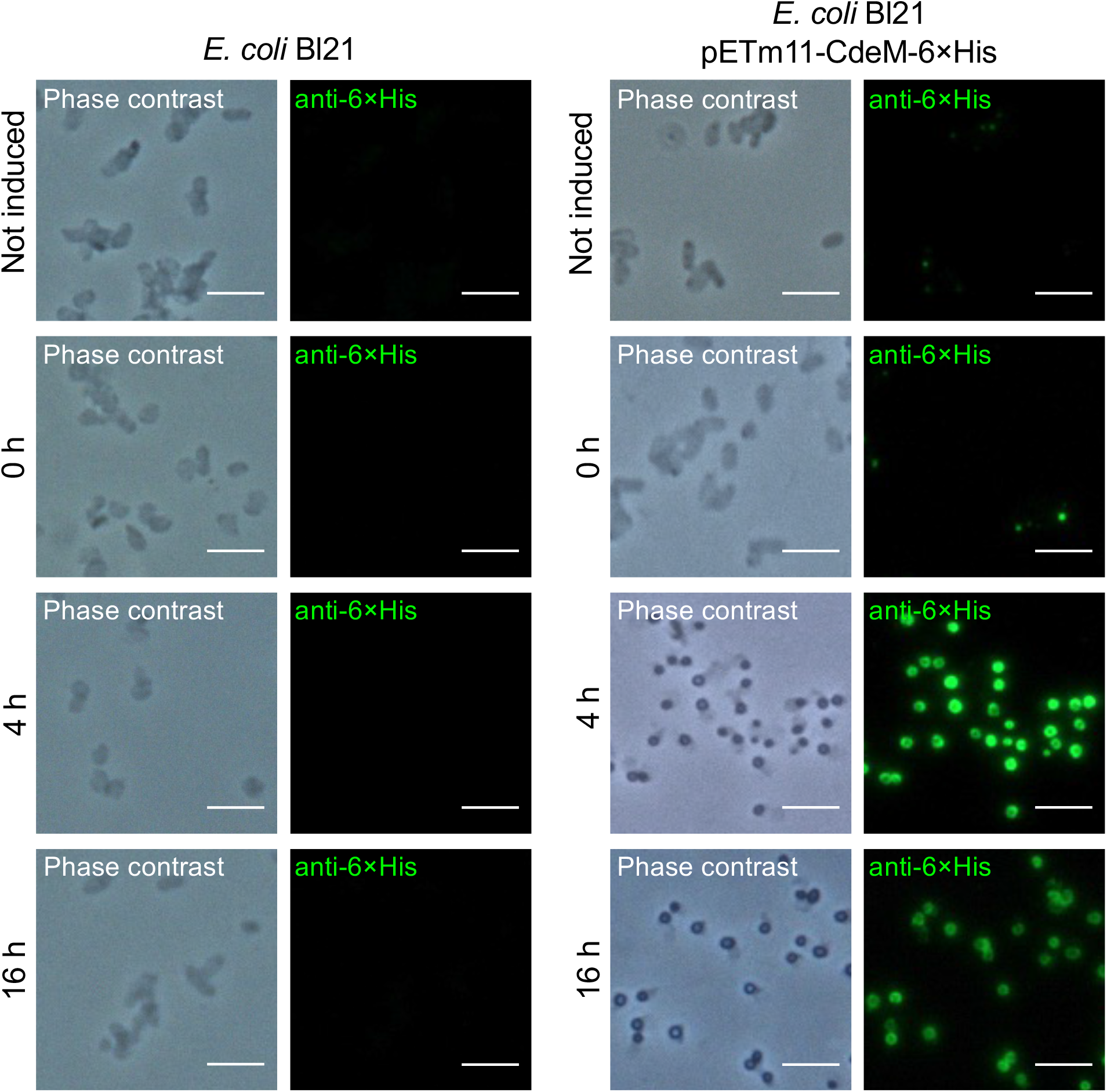
Time-dependent formation of CdeM-6×His inclusion bodies in *E. coli* BL21 cells. Representative phase-contrast and anti-6×His immunofluorescence images of control *Escherichia coli* BL21 cells and *E. coli* BL21 harboring the CdeM-6×His expression construct. Cells were examined under non-induced conditions and at 0, 4, and 16 h following induction. Control BL21 cells lacking the CdeM-6×His construct showed no detectable anti-6×His fluorescence at any time point. In contrast, BL21 cells expressing CdeM-6×His developed discrete intracellular anti-6×His-positive structures following induction, consistent with the formation of CdeM inclusion bodies. Little or no fluorescence was detected under non-induced conditions, whereas prominent intracellular anti-6×His-positive structures were observed at 4 and 16 h after induction. Phase-contrast images are shown alongside the corresponding anti-6×His fluorescence images. Scale bars are indicated in the micrographs.

## References

1. Evans CT, Safdar N. Current Trends in the Epidemiology and Outcomes of Clostridium difficile Infection. Clin Infect Dis. 2015;60 Suppl 2:S66–71. Epub 2015/04/30. doi: 10.1093/cid/civ140. PubMed PMID: 25922403.

2. Cohen SH, Gerding DN, Johnson S, Kelly CP, Loo VG, McDonald LC, et al. Clinical practice guidelines for Clostridium difficile infection in adults: 2010 update by the society for healthcare epidemiology of America (SHEA) and the infectious diseases society of America (IDSA). Infect Control Hosp Epidemiol. 2010;31(5):431–55. Epub 2010/03/24. doi: 10.1086/651706. PubMed PMID: 20307191.

3. Carroll KC, Bartlett JG. Biology of Clostridium difficile: implications for epidemiology and diagnosis. Annu Rev Microbiol. 2011;65:501–21. doi: 10.1146/annurev-micro-090110-102824. PubMed PMID: 21682645.

4. Lyras D, Adams V, Lucet I, Rood JI. The large resolvase TnpX is the only transposon-encoded protein required for transposition of the Tn4451/3 family of integrative mobilizable elements. Molecular microbiology. 2004;51(6):1787–800. PubMed PMID: 15009902.

5. Hussain HA, Roberts AP, Mullany P. Generation of an erythromycin-sensitive derivative of Clostridium difficile strain 630 (630Deltaerm) and demonstration that the conjugative transposon Tn916DeltaE enters the genome of this strain at multiple sites. Journal of medical microbiology. 2005;54(Pt 2):137–41. doi: 10.1099/jmm.0.45790-0. PubMed PMID: 15673506.

6. Sebaihia M, Wren BW, Mullany P, Fairweather NF, Minton N, Stabler R, et al. The multidrug-resistant human pathogen Clostridium difficile has a highly mobile, mosaic genome. Nat Genet. 2006;38(7):779–86. doi: 10.1038/ng1830. PubMed PMID: 16804543.

7. Rupnik M, Wilcox MH, Gerding DN. Clostridium difficile infection: new developments in epidemiology and pathogenesis. Nat Rev Microbiol. 2009;7(7):526–36. doi: 10.1038/nrmicro2164. PubMed PMID: 19528959.

8. Dawson LF, Valiente E, Wren BW. Clostridium difficile--a continually evolving and problematic pathogen. Infect Genet Evol. 2009;9(6):1410–7. doi: 10.1016/j.meegid.2009.06.005. PubMed PMID: 19539054.

9. Bouza E. Consequences of Clostridium difficile infection: understanding the healthcare burden. Clin Microbiol Infect. 2012;18 Suppl 6:5–12. Epub 2012/11/21. doi: 10.1111/1469-0691.12064. PubMed PMID: 23121549.

10. Bakken JS, Polgreen PM, Beekmann SE, Riedo FX, Streit JA. Treatment approaches including fecal microbiota transplantation for recurrent Clostridium difficile infection (RCDI) among infectious disease physicians. Anaerobe. 2013;24:20–4. Epub 2013/09/10. doi: 10.1016/j.anaerobe.2013.08.007. PubMed PMID: 24012687.

11. Aktories K, Papatheodorou P, Schwan C. Binary Clostridium difficile toxin (CDT) -A virulence factor disturbing the cytoskeleton. Anaerobe. 2018. Epub 2018/03/11. doi: 10.1016/j.anaerobe.2018.03.001. PubMed PMID: 29524654.

12. Brito GA, Carneiro-Filho B, Oria RB, Destura RV, Lima AA, Guerrant RL. Clostridium difficile toxin A induces intestinal epithelial cell apoptosis and damage: role of Gln and Ala-Gln in toxin A effects. Dig Dis Sci. 2005;50(7):1271–8. Epub 2005/07/29. doi: 10.1007/s10620-005-2771-x. PubMed PMID: 16047471.

13. Zemljic M, Rupnik M, Scarpa M, Anderluh G, Palu G, Castagliuolo I. Repetitive domain of Clostridium difficile toxin B exhibits cytotoxic effects on human intestinal epithelial cells and decreases epithelial barrier function. Anaerobe. 2010;16(5):527–32. Epub 2010/07/14. doi: 10.1016/j.anaerobe.2010.06.010. PubMed PMID: 20620216.

14. Carter GP, Chakravorty A, Pham Nguyen TA, Mileto S, Schreiber F, Li L, et al. Defining the Roles of TcdA and TcdB in Localized Gastrointestinal Disease, Systemic Organ Damage, and the Host Response during Clostridium difficile Infections. mBio. 2015;6(3):e00551. Epub 2015/06/04. doi: 10.1128/mBio.00551-15. PubMed PMID: 26037121; PubMed Central PMCID: PMCPMC4453007.

15. Deakin LJ, Clare S, Fagan RP, Dawson LF, Pickard DJ, West MR, et al. The Clostridium difficile spo0A gene is a persistence and transmission factor. Infection and immunity. 2012;80(8):2704–11. Epub 2012/05/23. doi: 10.1128/IAI.00147-12. PubMed PMID: 22615253; PubMed Central PMCID: PMCPMC3434595.

16. Barra-Carrasco J, Hernandez-Rocha C, Ibanez P, Guzman-Duran AM, Alvarez-Lobos M, Paredes-Sabja D. [Clostridium difficile spores and its relevance in the persistence and transmission of the infection]. Rev Chilena Infectol. 2014;31(6):694–703. Epub 2015/02/14. doi: 10.4067/S0716-10182014000600010. PubMed PMID: 25679927.

17. Srikhanta YN, Hutton ML, Awad MM, Drinkwater N, Singleton J, Day SL, et al. Cephamycins inhibit pathogen sporulation and effectively treat recurrent Clostridioides difficile infection. Nat Microbiol. 2019;4(12):2237–45. Epub 2019/08/14. doi: 10.1038/s41564-019-0519-1. PubMed PMID: 31406331.

18. Pizarro-Guajardo M, Diaz-Gonzalez F, Alvarez-Lobos M, Paredes-Sabja D. Characterization of Chicken IgY Specific to Clostridium difficile R20291 Spores and the Effect of Oral Administration in Mouse Models of Initiation and Recurrent Disease. Front Cell Infect Microbiol. 2017;7:365. Epub 2017/09/01. doi: 10.3389/fcimb.2017.00365. PubMed PMID: 28856119; PubMed Central PMCID: PMCPMC5557795.

19. Castro-Córdova P, Mora-Uribe P, Reyes-Ramírez R, Cofré-Araneda G, Orozco-Aguilar J, Brito-Silva C, et al. Entry of spores into intestinal epithelial cells contributes to recurrence of *Clostridioides difficile* infection. Nat Commun. 2021;12(1):1140. Epub 2021/02/18. doi: 10.1038/s41467-021-21355-5. PubMed PMID: 33602902.

20. Paredes-Sabja D, Sarker MR. Adherence of Clostridium difficile spores to Caco-2 cells in culture. Journal of medical microbiology. 2012;61(Pt 9):1208–18. Epub 2012/05/19. doi: 10.1099/jmm.0.043687-0. PubMed PMID: 22595914.

21. Mora-Uribe P, Miranda-Cardenas C, Castro-Cordova P, Gil F, Calderon I, Fuentes JA, et al. Characterization of the Adherence of Clostridium difficile Spores: The Integrity of the Outermost Layer Affects Adherence Properties of Spores of the Epidemic Strain R20291 to Components of the Intestinal Mucosa. Front Cell Infect Microbiol. 2016;6:99. Epub 2016/10/08. doi: 10.3389/fcimb.2016.00099. PubMed PMID: 27713865; PubMed Central PMCID: PMCPMC5031699.

22. Lopez-Garcia OK, Pizarro-Guajardo M, Kocurek KI, Rezenom YH, Paredes-Sabja D. Coupling Far-Western Blotting with Peptide Microarrays Reveals Novel E-Cadherin Spore-Surface Ligands in Clostridioides difficile. J Proteome Res. 2026;25(7):3487–506. Epub 20260616. doi: 10.1021/acs.jproteome.5c01166. PubMed PMID: 42299067; PubMed Central PMCID: PMCPMC13339774.

23. Castro-Córdova P, Otto-Medina M, Montes-Bravo N, Brito-Silva C, Lacy DB, Paredes-Sabja D. Redistribution of the Novel *Clostridioides difficile* Spore Adherence Receptor E-Cadherin by TcdA and TcdB Increases Spore Binding to Adherens Junctions. Infect Immun. 2023;91(1):e0047622. Epub 20221130. doi: 10.1128/iai.00476-22. PubMed PMID: 36448839; PubMed Central PMCID: PMCPMC9872679.

24. Hovingh ES, van den Broek B, Jongerius I. Hijacking Complement Regulatory Proteins for Bacterial Immune Evasion. Front Microbiol. 2016;7:2004. Epub 20161220. doi: 10.3389/fmicb.2016.02004. PubMed PMID: 28066340; PubMed Central PMCID: PMCPMC5167704.

25. Kunz N, Kemper C. Complement Has Brains-Do Intracellular Complement and Immunometabolism Cooperate in Tissue Homeostasis and Behavior? Front Immunol. 2021;12:629986. Epub 20210225. doi: 10.3389/fimmu.2021.629986. PubMed PMID: 33717157; PubMed Central PMCID: PMCPMC7946832.

26. Pratt JR, Abe K, Miyazaki M, Zhou W, Sacks SH. In situ localization of C3 synthesis in experimental acute renal allograft rejection. Am J Pathol. 2000;157(3):825–31. doi: 10.1016/s0002-9440(10)64596-8. PubMed PMID: 10980122; PubMed Central PMCID: PMCPMC1885894.

27. Liszewski MK, Kolev M, Le Friec G, Leung M, Bertram PG, Fara AF, et al. Intracellular complement activation sustains T cell homeostasis and mediates effector differentiation. Immunity. 2013;39(6):1143–57. Epub 20131205. doi: 10.1016/j.immuni.2013.10.018. PubMed PMID: 24315997; PubMed Central PMCID: PMCPMC3865363.

28. Lubbers R, van Essen MF, van Kooten C, Trouw LA. Production of complement components by cells of the immune system. Clin Exp Immunol. 2017;188(2):183–94. Epub 20170324. doi: 10.1111/cei.12952. PubMed PMID: 28249350; PubMed Central PMCID: PMCPMC5383442.

29. Kulkarni HS, Elvington ML, Perng YC, Liszewski MK, Byers DE, Farkouh C, et al. Intracellular C3 Protects Human Airway Epithelial Cells from Stress-associated Cell Death. Am J Respir Cell Mol Biol. 2019;60(2):144–57. doi: 10.1165/rcmb.2017-0405OC. PubMed PMID: 30156437; PubMed Central PMCID: PMCPMC6376412.

30. King BC, Kulak K, Krus U, Rosberg R, Golec E, Wozniak K, et al. Complement Component C3 Is Highly Expressed in Human Pancreatic Islets and Prevents β Cell Death via ATG16L1 Interaction and Autophagy Regulation. Cell Metab. 2019;29(1):202–10.e6. Epub 20181004. doi: 10.1016/j.cmet.2018.09.009. PubMed PMID: 30293775.

31. Kolev M, West EE, Kunz N, Chauss D, Moseman EA, Rahman J, et al. Diapedesis-Induced Integrin Signaling via LFA-1 Facilitates Tissue Immunity by Inducing Intrinsic Complement C3 Expression in Immune Cells. Immunity. 2020;52(3):513–27.e8. doi: 10.1016/j.immuni.2020.02.006. PubMed PMID: 32187519; PubMed Central PMCID: PMCPMC7111494.

32. Lloyd-Price J, Arze C, Ananthakrishnan AN, Schirmer M, Avila-Pacheco J, Poon TW, et al. Multi-omics of the gut microbial ecosystem in inflammatory bowel diseases. Nature. 2019;569(7758):655–62. Epub 20190529. doi: 10.1038/s41586-019-1237-9. PubMed PMID: 31142855; PubMed Central PMCID: PMCPMC6650278.

33. D’Auria KM, Kolling GL, Donato GM, Warren CA, Gray MC, Hewlett EL, et al. In vivo physiological and transcriptional profiling reveals host responses to Clostridium difficile toxin A and toxin B. Infection and immunity. 2013;81(10):3814–24. doi: 10.1128/IAI.00869-13. PubMed PMID: 23897615; PubMed Central PMCID: PMCPMC3811747.

34. Gu C, Jenkins SA, Xue Q, Xu Y. Activation of the classical complement pathway by Bacillus anthracis is the primary mechanism for spore phagocytosis and involves the spore surface protein BclA. J Immunol. 2012;188(9):4421–31. Epub 2012/03/24. doi: 10.4049/jimmunol.1102092. PubMed PMID: 22442442; PubMed Central PMCID: PMC3331890.

35. Premanandan C, Storozuk CA, Clay CD, Lairmore MD, Schlesinger LS, Phipps AJ. Complement protein C3 binding to Bacillus anthracis spores enhances phagocytosis by human macrophages. Microbial pathogenesis. 2009;46(6):306–14. Epub 2009/03/31. doi: 10.1016/j.micpath.2009.03.004. PubMed PMID: 19328844.

36. Chung MC, Tonry JH, Narayanan A, Manes NP, Mackie RS, Gutting B, et al. Bacillus anthracis interacts with plasmin(ogen) to evade C3b-dependent innate immunity. PloS one. 2011;6(3):e18119. Epub 2011/04/06. doi: 10.1371/journal.pone.0018119. PubMed PMID: 21464960; PubMed Central PMCID: PMCPMC3064659.

37. Xue Q, Gu C, Rivera J, Hook M, Chen X, Pozzi A, et al. Entry of *Bacillus anthracis* spores into epithelial cells is mediated by the spore surface protein BclA, integrin alpha2beta1 and complement component C1q. Cell Microbiol. 2011;13(4):620–34. Epub 2010/12/08. doi: 10.1111/j.1462-5822.2010.01558.x. PubMed PMID: 21134100.

38. Arato V, Gasperini G, Giusti F, Ferlenghi I, Scarselli M, Leuzzi R. Dual role of the colonization factor CD2831 in Clostridium difficile pathogenesis. Sci Rep. 2019;9(1):5554. Epub 2019/04/05. doi: 10.1038/s41598-019-42000-8. PubMed PMID: 30944377; PubMed Central PMCID: PMCPMC6447587.

39. Gunji N, Katakura K, Abe K, Kawashima K, Fujiwara T, Onizawa M, et al. Upregulation of complement C1q reflects mucosal regeneration in a mouse model of colitis. Medical molecular morphology. 2020. doi: 10.1007/s00795-020-00266-2.

40. Abt MC, McKenney PT, Pamer EG. Clostridium difficile colitis: pathogenesis and host defence. Nat Rev Microbiol. 2016;14(10):609–20. doi: 10.1038/nrmicro.2016.108. PubMed PMID: 27573580; PubMed Central PMCID: PMCPMC5109054.

41. Sharma S, Bhatnagar R, Gaur D. Complement Evasion Strategies of Human Pathogenic Bacteria. Indian J Microbiol. 2020;60(3):283–96. Epub 20200424. doi: 10.1007/s12088-020-00872-9. PubMed PMID: 32655196; PubMed Central PMCID: PMCPMC7329968.

42. Wu M, Zheng W, Song X, Bao B, Wang Y, Ramanan D, et al. Gut complement induced by the microbiota combats pathogens and spares commensals. Cell. 2024;187(4):897–913.e18. Epub 20240126. doi: 10.1016/j.cell.2023.12.036. PubMed PMID: 38280374; PubMed Central PMCID: PMCPMC10922926.

43. Andoh A, Fujiyama Y, Sakumoto H, Uchihara H, Kimura T, Koyama S, et al. Detection of complement C3 and factor B gene expression in normal colorectal mucosa, adenomas and carcinomas. Clin Exp Immunol. 1998;111(3):477–83. PubMed PMID: 9528886; PubMed Central PMCID: PMCPMC1904873.

44. Castro-Cordova P, Mora-Uribe P, Reyes-Ramirez R, Cofre-Araneda G, Orozco-Aguilar J, Brito-Silva C, et al. Entry of spores into intestinal epithelial cells contributes to recurrence of Clostridioides difficile infection. Nat Commun. 2021;12(1):1140. Epub 2021/02/20. doi: 10.1038/s41467-021-21355-5. PubMed PMID: 33602902.

45. Pizarro-Guajardo M, Calderon-Romero P, Castro-Cordova P, Mora-Uribe P, Paredes-Sabja D. Ultrastructural Variability of the Exosporium Layer of *Clostridium difficile* Spores. Appl Environ Microbiol. 2016;82(7):2202–9. Epub 2016/02/07. doi: 10.1128/AEM.03410-15. PubMed PMID: 26850296; PubMed Central PMCID: PMCPMC4807528.

46. Pizarro-Guajardo M, Calderón-Romero P, Paredes-Sabja D. Ultrastructure Variability of the Exosporium Layer of *Clostridium difficile* Spores from Sporulating Cultures and Biofilms. Appl Environ Microbiol. 2016;82(19):5892–8. Epub 20160916. doi: 10.1128/aem.01463-16. PubMed PMID: 27474709; PubMed Central PMCID: PMCPMC5038037.

47. Calderon-Romero P, Castro-Cordova P, Reyes-Ramirez R, Milano-Cespedes M, Guerrero-Araya E, Pizarro-Guajardo M, et al. Clostridium difficile exosporium cysteine-rich proteins are essential for the morphogenesis of the exosporium layer, spore resistance, and affect C. difficile pathogenesis. PLoS Pathog. 2018;14(8):e1007199. Epub 2018/08/09. doi: 10.1371/journal.ppat.1007199. PubMed PMID: 30089172; PubMed Central PMCID: PMCPMC6101409.

48. Romero-Rodriguez A, Troncoso-Cotal S, Guerrero-Araya E, Paredes-Sabja D. The Clostridioides difficile Cysteine-Rich Exosporium Morphogenetic Protein, CdeC, Exhibits Self-Assembly Properties That Lead to Organized Inclusion Bodies in Escherichia coli. mSphere. 2020;5(6). Epub 2020/11/20. doi: 10.1128/mSphere.01065-20. PubMed PMID: 33208520; PubMed Central PMCID: PMCPMC7677010.

49. Brito-Silva C, Pizarro-Cerda J, Gil F, Paredes-Sabja D. Identification of Escherichia coli strains for the heterologous overexpression of soluble Clostridium difficile exosporium proteins. Journal of microbiological methods. 2018;154:46–51. Epub 2018/10/07. doi: 10.1016/j.mimet.2018.10.002. PubMed PMID: 30291882.

50. Wang Y, Jenkins SA, Gu C, Shree A, Martinez-Moczygemba M, Herold J, et al. Bacillus anthracis Spore Surface Protein BclA Mediates Complement Factor H Binding to Spores and Promotes Spore Persistence. PLoS Pathog. 2016;12(6):e1005678. Epub 20160615. doi: 10.1371/journal.ppat.1005678. PubMed PMID: 27304426; PubMed Central PMCID: PMCPMC4909234.

51. Lessa FC, Mu Y, Bamberg WM, Beldavs ZG, Dumyati GK, Dunn JR, et al. Burden of *Clostridium difficile* infection in the United States. N Engl J Med. 2015;372(9):825–34. Epub 2015/02/26. doi: 10.1056/NEJMoa1408913. PubMed PMID: 25714160.

52. Kelly CP. Can we identify patients at high risk of recurrent *Clostridium difficile* infection? Clin Microbiol Infect. 2012;18(6):21–7. doi: 10.1111/1469-0691.12046. PubMed PMID: 23121551.

53. Zhang S, Palazuelos-Munoz S, Balsells EM, Nair H, Chit A, Kyaw MH. Cost of hospital management of Clostridium difficile infection in United States-a meta-analysis and modelling study. BMC Infect Dis. 2016;16(1):447. Epub 20160825. doi: 10.1186/s12879-016-1786-6. PubMed PMID: 27562241; PubMed Central PMCID: PMCPMC5000548.

54. Dong D, Su T, Chen W, Wang D, Xue Y, Lu Q, et al. *Clostridioides difficile* aggravates dextran sulfate solution (DSS)-induced colitis by shaping the gut microbiota and promoting neutrophil recruitment. Gut Microbes. 2023;15(1):2192478. doi: 10.1080/19490976.2023.2192478. PubMed PMID: 36951545; PubMed Central PMCID: PMCPMC10038061.

55. Pruss KM, Sonnenburg JL. *C. difficile* exploits a host metabolite produced during toxin-mediated disease. Nature. 2021;593(7858):261–5. Epub 20210428. doi: 10.1038/s41586-021-03502-6. PubMed PMID: 33911281.

56. Fletcher JR, Pike CM, Parsons RJ, Rivera AJ, Foley MH, McLaren MR, et al. *Clostridioides difficile* exploits toxin-mediated inflammation to alter the host nutritional landscape and exclude competitors from the gut microbiota. Nat Commun. 2021;12(1):462. Epub 20210119. doi: 10.1038/s41467-020-20746-4. PubMed PMID: 33469019; PubMed Central PMCID: PMCPMC7815924.

57. Albertí S, Marqués G, Camprubí S, Merino S, Tomás JM, Vivanco F, et al. C1q binding and activation of the complement classical pathway by Klebsiella pneumoniae outer membrane proteins. Infect Immun. 1993;61(3):852–60. doi: 10.1128/iai.61.3.852-860.1993. PubMed PMID: 8432605; PubMed Central PMCID: PMCPMC302811.

58. Roumenina LT, Popov KT, Bureeva SV, Kojouharova M, Gadjeva M, Rabheru S, et al. Interaction of the globular domain of human C1q with Salmonella typhimurium lipopolysaccharide. Biochim Biophys Acta. 2008;1784(9):1271–6. Epub 20080510. doi: 10.1016/j.bbapap.2008.04.029. PubMed PMID: 18513495.

59. Terrasse R, Amoroso A, Vernet T, Di Guilmi AM. Streptococcus pneumoniae GAPDH Is Released by Cell Lysis and Interacts with Peptidoglycan. PLoS One. 2015;10(4):e0125377. Epub 20150430. doi: 10.1371/journal.pone.0125377. PubMed PMID: 25927608; PubMed Central PMCID: PMCPMC4415926.

60. De Gaetano GV, Coppolino F, Lentini G, Famà A, Cullotta C, Raffaele I, et al. Streptococcus pneumoniae binds collagens and C1q via the SSURE repeats of the PfbB adhesin. Mol Microbiol. 2022;117(6):1479–92. Epub 20220530. doi: 10.1111/mmi.14920. PubMed PMID: 35570359; PubMed Central PMCID: PMCPMC9328315.

61. Valotteau C, Prystopiuk V, Pietrocola G, Rindi S, Peterle D, De Filippis V, et al. Single-Cell and Single-Molecule Analysis Unravels the Multifunctionality of the Staphylococcus aureus Collagen-Binding Protein Cna. ACS Nano. 2017;11(2):2160–70. Epub 20170202. doi: 10.1021/acsnano.6b08404. PubMed PMID: 28151647.

62. Hong HA, Ferreira WT, Hosseini S, Anwar S, Hitri K, Wilkinson AJ, et al. The Spore Coat Protein CotE Facilitates Host Colonization by *Clostridium difficile*. J Infect Dis. 2017;216(11):1452–9. Epub 2017/10/03. doi: 10.1093/infdis/jix488. PubMed PMID: 28968845; PubMed Central PMCID: PMCPMC5853579.

63. Permpoonpattana P, Tolls EH, Nadem R, Tan S, Brisson A, Cutting SM. Surface layers of *Clostridium difficile* endospores. J Bacteriol. 2011;193(23):6461–70. Epub 20110923. doi: 10.1128/jb.05182-11. PubMed PMID: 21949071; PubMed Central PMCID: PMCPMC3232898.

64. Li K, Zhou W, Hong Y, Sacks SH, Sheerin NS. Synergy between type 1 fimbriae expression and C3 opsonisation increases internalisation of E. coli by human tubular epithelial cells. BMC Microbiol. 2009;9:64. Epub 20090331. doi: 10.1186/1471-2180-9-64. PubMed PMID: 19335887; PubMed Central PMCID: PMCPMC2670304.

65. Papatheodorou P, Barth H, Minton N, Aktories K. Cellular Uptake and Mode-of-Action of Clostridium difficile Toxins. Advances in experimental medicine and biology. 2018;1050:77–96. Epub 2018/02/01. doi: 10.1007/978-3-319-72799-8_6. PubMed PMID: 29383665.

66. Voth DE, Ballard JD. Clostridium difficile toxins: mechanism of action and role in disease. Clin Microbiol Rev. 2005;18(2):247–63. doi: 10.1128/CMR.18.2.247-263.2005. PubMed PMID: 15831824; PubMed Central PMCID: PMCPMC1082799.

67. Bartlett JG, Chang TW, Gurwith M, Gorbach SL, Onderdonk AB. Antibiotic-associated pseudomembranous colitis due to toxin-producing clostridia. N Engl J Med. 1978;298(10):531–4. doi: 10.1056/NEJM197803092981003. PubMed PMID: 625309.

68. Castro-Córdova P, Mendoza-León MJ, Paredes-Sabja D. Using a ligate intestinal loop mouse model to investigate *Clostridioides difficile* adherence to the intestinal mucosa in aged mice. PLoS One. 2021;16(12):e0261081. Epub 20211222. doi: 10.1371/journal.pone.0261081. PubMed PMID: 34936648; PubMed Central PMCID: PMCPMC8694449.

69. Johansson ME, Larsson JM, Hansson GC. The two mucus layers of colon are organized by the MUC2 mucin, whereas the outer layer is a legislator of host-microbial interactions. Proceedings of the National Academy of Sciences of the United States of America. 2011;108 Suppl 1:4659–65. Epub 2010/07/10. doi: 10.1073/pnas.1006451107. PubMed PMID: 20615996; PubMed Central PMCID: PMCPMC3063600.

70. Chen X, Katchar K, Goldsmith JD, Nanthakumar N, Cheknis A, Gerding DN, et al. A mouse model of Clostridium difficile-associated disease. Gastroenterology. 2008;135(6):1984–92. Epub 2008/10/14. doi: 10.1053/j.gastro.2008.09.002. PubMed PMID: 18848941.

71. Pizarro-Guajardo M, Ravanal MC, Paez MD, Callegari E, Paredes-Sabja D. Identification of *Clostridium difficile* Immunoreactive Spore Proteins of the Epidemic Strain R20291. Proteomics Clin Appl. 2018:e1700182. Epub 2018/03/25. doi: 10.1002/prca.201700182. PubMed PMID: 29573213.

72. Diaz-Gonzalez F, Milano M, Olguin-Araneda V, Pizarro-Cerda J, Castro-Cordova P, Tzeng SC, et al. Protein composition of the outermost exosporium-like layer of *Clostridium difficile* 630 spores. J Proteomics. 2015;123:1–13. Epub 2015/04/08. doi: 10.1016/j.jprot.2015.03.035. PubMed PMID: 25849250.

73. Kelstrup CD, Young C, Lavallee R, Nielsen ML, Olsen JV. Optimized fast and sensitive acquisition methods for shotgun proteomics on a quadrupole orbitrap mass spectrometer. J Proteome Res. 2012;11(6):3487–97. Epub 20120510. doi: 10.1021/pr3000249. PubMed PMID: 22537090.

74. Sun L, Zhu G, Dovichi NJ. Comparison of the LTQ-Orbitrap Velos and the Q-Exactive for proteomic analysis of 1-1000 ng RAW 264.7 cell lysate digests. Rapid Commun Mass Spectrom. 2013;27(1):157–62. doi: 10.1002/rcm.6437. PubMed PMID: 23239329; PubMed Central PMCID: PMCPMC3673017.

75. Sun L, Zhu G, Dovichi NJ. Comparison of the LTQ-Orbitrap Velos and the Q-Exactive for proteomic analysis of 1-1000 ng RAW 264.7 cell lysate digests. Rapid communications in mass spectrometry : RCM. 2013;27(1):157–62. Epub 2012/12/15. doi: 10.1002/rcm.6437. PubMed PMID: 23239329; PubMed Central PMCID: PMCPMC3673017.

