## Supplementary figures and images for "Gut complement C1q and C3 interact with *Clostridioides difficile* spores via CdeM and contribute to pathogenesis"

### Figure Supplemental

#### Supplementary Figure 1

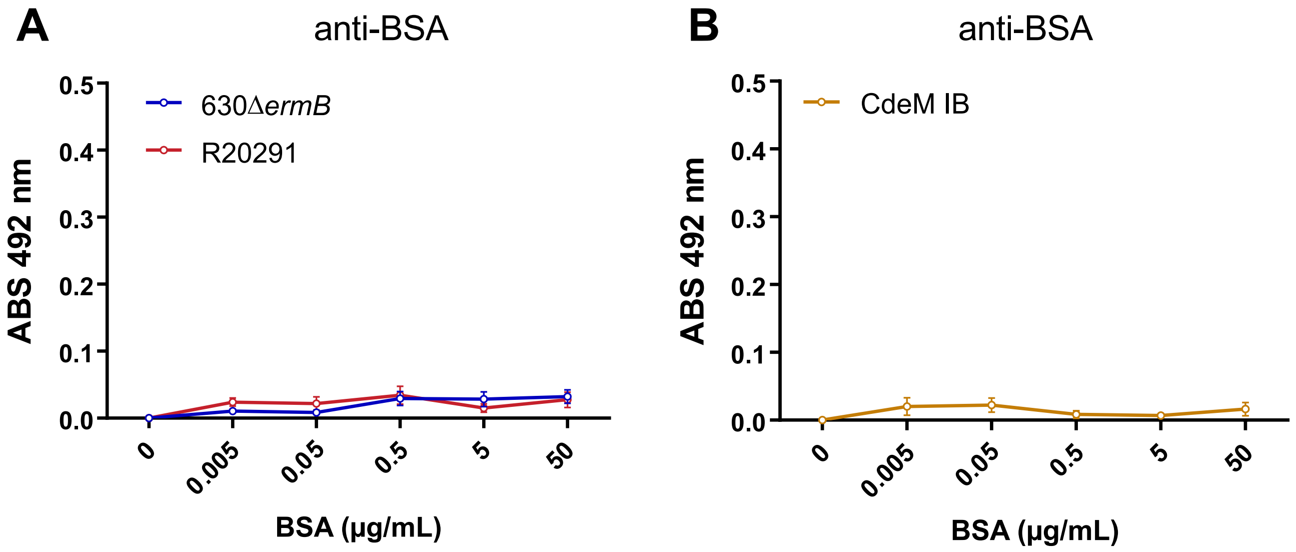

#### Supplementary Figure 2

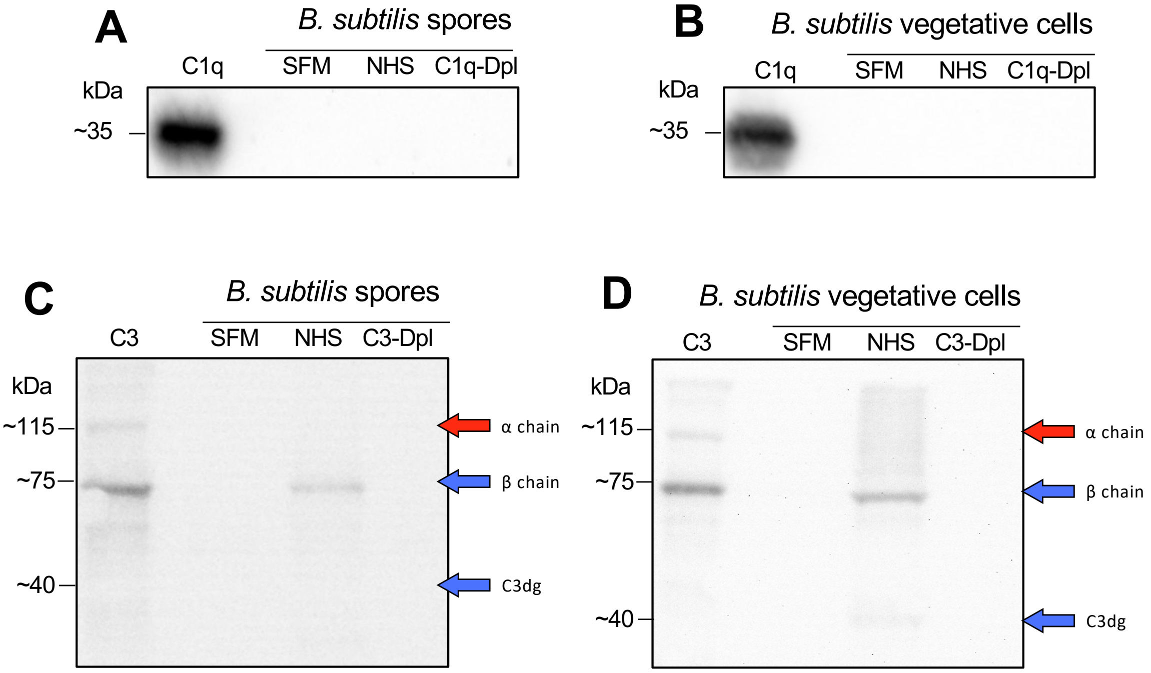

#### Supplementary Figure 3

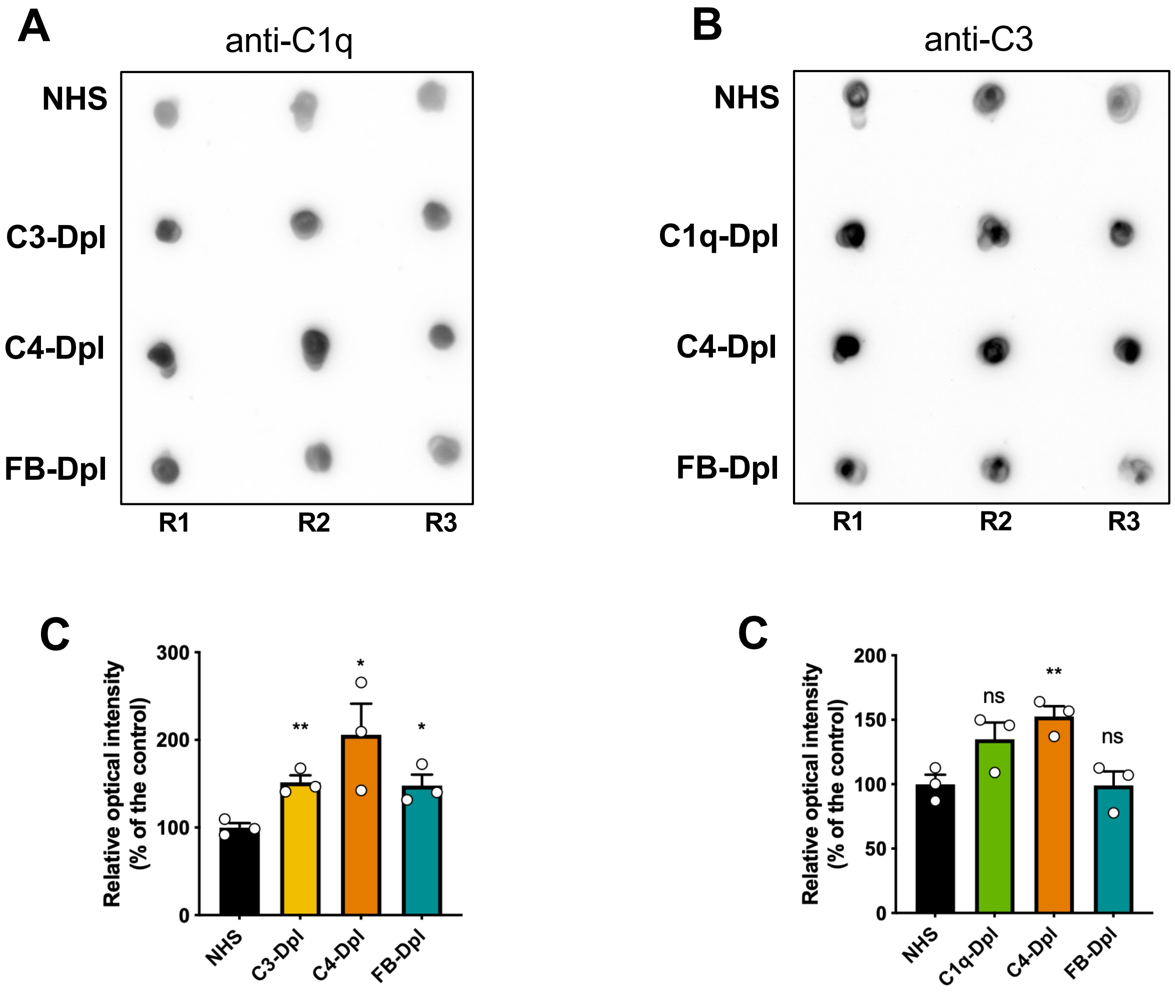

#### Supplementary Figure 4

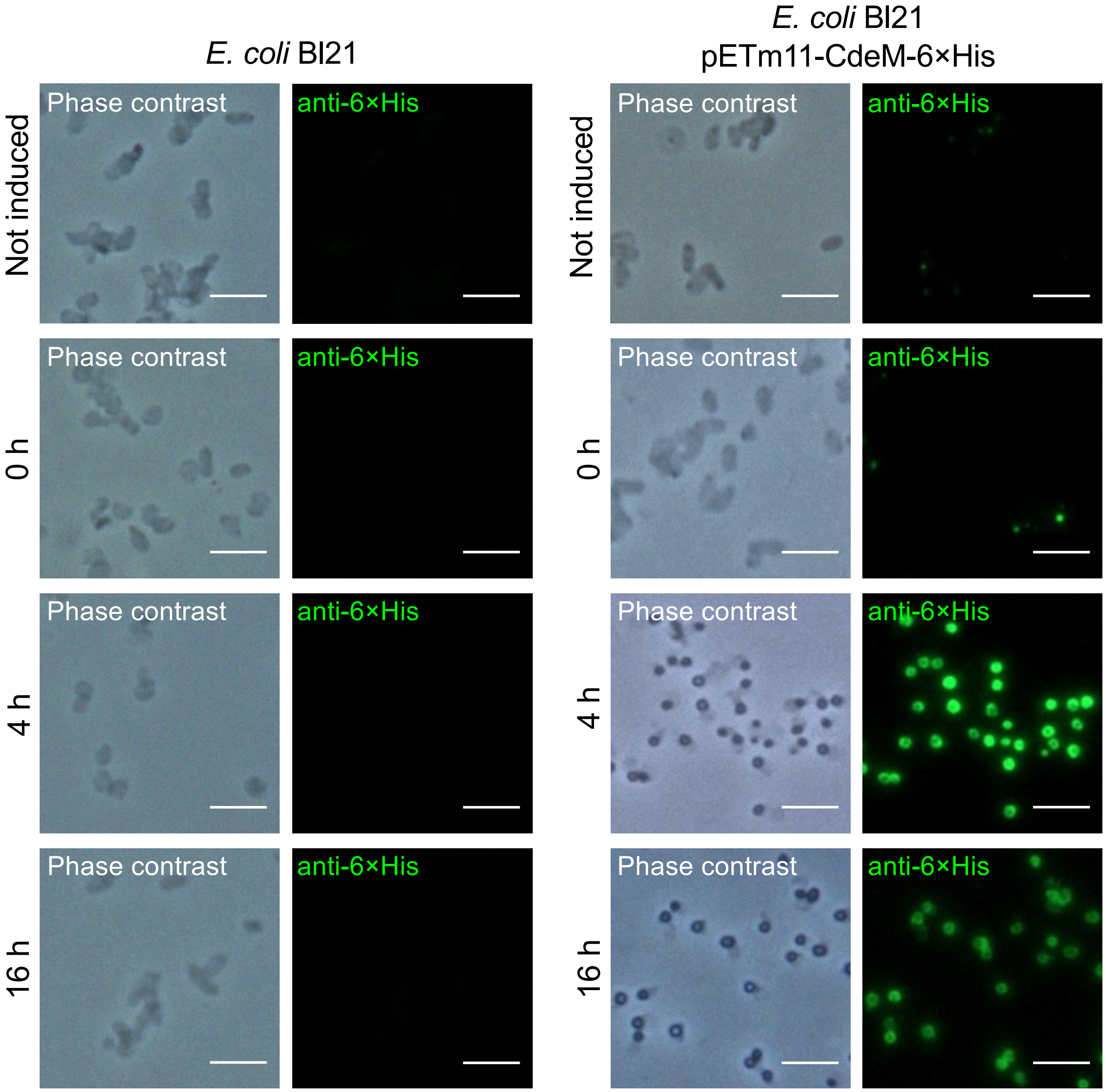
