## Supplementary material for "Gut complement C1q and C3 interact with *Clostridioides difficile* spores via CdeM and contribute to pathogenesis": Table S1

| TABLE S1. Bacterial strains and plasmids used | | |
| --- | --- | --- |
| **Strain** | **Relevant characteristic** | **Source/Reference** |
| *C. difficile* 630∆*ermB* | A laboratory strain with erythromycin sensitive derivative of *C. difficile* strain 630 | ^1^ |
| *C. difficile* R20291 | Ribotype 027, epidemically relevant strain | ^2^ |
| *C. difficile* 630∆*ermB*-*bclA1::CT1050a* | ClosTron insertional mutant in *bclA1* | ^3^ |
| *C. difficile* 630∆*ermB*-*bclA2::CT150a* | ClosTron insertional mutant in *bclA2* | ^3^ |
| *C. difficile* 630∆*ermB*-*bclA3::CT125s* | ClosTron insertional mutant in *bclA3* | ^3^ |
| R20291 Δ*pyrE/pyrE^+^* | R20291 isogenic *pyrE* mutant complemented with wild-type *pyrE* into the *pyrE* loci | ^4^ |
| R20291 Δ*bclA3* | R20291 isogenic *bclA3* mutant | ^4^ |
| R20291 Δ*bclA3/bclA3^+^* | R20291 isogenic *bclA3* mutant complemented with wild-type *bclA3* in the *pyrE* loci | ^4^ |
| *Bacillus subtilis* PY79 | Wild-type *Bacillus subtilis* |  |
| *Escherichia coli Bl21*Bl21 (DE3) | Bl21 strain carrying a plasmid that contain extra copies of the argU, ileY, and leuW tRNA genes (rare tRNAs codons) | Agilent, U.S.A. |
| ***Plasmids*** |  |  |
| pARR21 | Expression plasmid carring cdeM from R20291 cloned into NcoI and XhoI sites of pETM11, giving a CdeM-6xHis tag fusion. | ^5^ |

**References**

1	Hussain, H. A., Roberts, A. P. & Mullany, P. Generation of an erythromycin-sensitive derivative of *Clostridium difficile* strain 630 (630Deltaerm) and demonstration that the conjugative transposon Tn916DeltaE enters the genome of this strain at multiple sites. *J Med Microbiol* **54**, 137-141, doi:10.1099/jmm.0.45790-0 (2005).

2	McEllistrem, M. C., Carman, R. J., Gerding, D. N., Genheimer, C. W. & Zheng, L. A hospital outbreak of *Clostridium difficile* disease associated with isolates carrying binary toxin genes. *Clin Infect Dis* **40**, 265-272, doi:10.1086/427113 (2005).

3	Phetcharaburanin, J. *et al.* The spore-associated protein BclA1 affects the susceptibility of animals to colonization and infection by *Clostridium difficile*. *Mol Microbiol* **92**, 1025-1038, doi:10.1111/mmi.12611 (2014).

4	Castro-Córdova, P. *et al.* Entry of spores into intestinal epithelial cells contributes to recurrence of *Clostridioides difficile* infection. *Nat Commun* **12**, 1140, doi:10.1038/s41467-021-21355-5 (2021).

5	Romero-Rodríguez, A., Troncoso-Cotal, S., Guerrero-Araya, E. & Paredes-Sabja, D. The *Clostridioides difficile* Cysteine-Rich Exosporium Morphogenetic Protein, CdeC, Exhibits Self-Assembly Properties That Lead to Organized Inclusion Bodies in Escherichia coli. *mSphere* **5**, doi:10.1128/mSphere.01065-20 (2020).
